# Systematic comparative analysis of single cell RNA-sequencing methods

**DOI:** 10.1101/632216

**Authors:** Jiarui Ding, Xian Adiconis, Sean K. Simmons, Monika S. Kowalczyk, Cynthia C. Hession, Nemanja D. Marjanovic, Travis K. Hughes, Marc H. Wadsworth, Tyler Burks, Lan T. Nguyen, John Y. H. Kwon, Boaz Barak, William Ge, Amanda J. Kedaigle, Shaina Carroll, Shuqiang Li, Nir Hacohen, Orit Rozenblatt-Rosen, Alex K. Shalek, Alexandra-Chloé Villani, Aviv Regev, Joshua Z. Levin

## Abstract

A multitude of single-cell RNA sequencing methods have been developed in recent years, with dramatic advances in scale and power, and enabling major discoveries and large scale cell mapping efforts. However, these methods have not been systematically and comprehensively benchmarked. Here, we directly compare seven methods for single cell and/or single nucleus profiling from three types of samples – cell lines, peripheral blood mononuclear cells and brain tissue – generating 36 libraries in six separate experiments in a single center. To analyze these datasets, we developed and applied scumi, a flexible computational pipeline that can be used for any scRNA-seq method. We evaluated the methods for both basic performance and for their ability to recover known biological information in the samples. Our study will help guide experiments with the methods in this study as well as serve as a benchmark for future studies and for computational algorithm development.

---

Single-cell RNA sequencing (scRNA-seq) has emerged as a central tool for identifying and characterizing cell types, states, lineages, and circuitry^1–3^. The rapid growth in the scale and robustness of lab protocols and associated computational tools has opened the way to substantial scientific discoveries and to an international initiative, the Human Cell Atlas (HCA), to build comprehensive reference maps of all human cells^4^.

ScRNA-seq methods differ in how they tag transcripts for their cell-of-origin and generate libraries for sequencing. Low-throughput, plate-based methods^5, 6^ sort a cell into a well of a multi-well plate; high-throughput, bead-based methods distribute a cell suspension into tiny droplets^7–9^ or wells^10, 11^ containing reagents and barcoded beads to produce a single droplet or well with one cell and one bead that is used to mark all the cDNA generated from that cell; and scalable, combinatorial indexing methods reverse transcribe and barcode mRNAs *in situ* inside each cell or nucleus, without physically isolating single cells^12–14^ (**Supplementary Fig. 1**).

ScRNA-seq remains a rapidly evolving field^15^ with continued development of new methods and improvement of existing ones. There is thus an urgent need to provide comparison and benchmarking information to help guide users make informed choices based on each method’s capabilities and limitations, compare newly proposed methods to existing ones, and identify shared weaknesses as targets for experimental improvement. Prior comparisons of scRNA-seq methods^16–21^, though useful, have several shortcomings. Many are outdated given the fast-paced field, or are incomplete, inapplicable (e.g., not actually performed with single cells), or insufficiently controlled (e.g., performed using different samples for comparisons); others limit their assessment to basic technical factors, but do not assess the key benchmark of ability to recover meaningful biological information, such as population heterogeneity and structure. In particular, comparisons often focused on cultured cell lines, even though most scRNA-seq studies focus in practice on tissues and primary cells. An updated comparison of scRNA-seq methods is needed to help scRNA-seq method developers better understand the advantages and disadvantages of different methods, researchers to choose appropriate experimental methods to answer their biological questions, and computational method developers to create new data and software packages for data processing.

Here, we systematically and directly compared seven methods (**Fig. 1**, **Supplementary Fig. 1**), including two low-throughput plate-based methods (Smart-seq2^5^ and CEL-Seq2^6^) and five high-throughput methods (10x Chromium^9^, Drop-seq^8^, Seq-Well^10^, inDrops^7^, and sci-RNA-seq^12^), producing expression profiles from ∼92,000 cells overall. We analyzed at either the single cell or single nucleus level three sample types – a mixture of human and mouse cell lines, human peripheral blood mononuclear cells (PBMCs), and mouse cortex, each sample with two replicates – to generate a total of 36 different scRNA-seq libraries. For mouse cortex we tested four single nucleus RNA-seq methods^9, 12, 22, 23^, in what is, to the best of our knowledge, the first such comparison. For each sample type, we characterized performance with basic metrics, and for PBMC and cortex libraries, we further examined how well the methods capture biological information. This is a critical part of most scRNA-seq studies, but has not been evaluated in other comparison studies that used only relatively homogeneous cell lines^16, 20^. Our study provides both immediate guidance on each method’s relative performance, and an experimental and computational benchmarking framework to assess future techniques. For the low-throughput methods, Smart-seq2 and CEL-Seq2 performed similarly, though the latter may be affected more by contaminating reads from other cells. Among the high-throughput methods, 10x Chromium was the top performer, both for single cell and single nucleus data.

**Figure 1.**
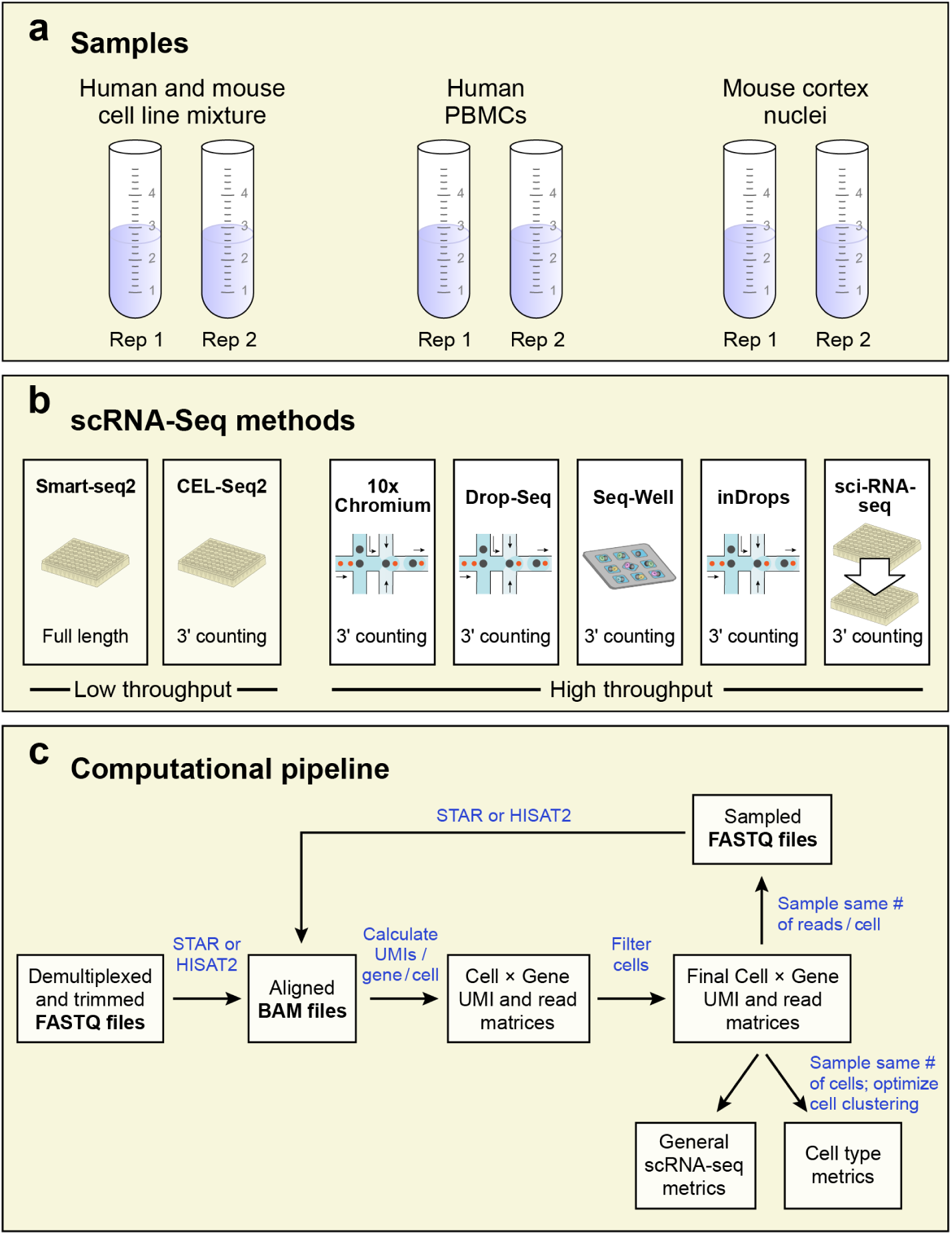
Description of methods & experimental design. (**a**) samples, (**b**) scRNA-seq methods, (**c**) computational pipeline summary. Cell line mixtures tested with all methods. PBMCs tested with all methods except sci-RNA-seq. Cortex nuclei tested with Smart-seq2, 10x Chromium, Drop-seq (aka DroNc-seq for nuclei), and sci-RNA-seq. Additional details can be found in **Supplementary Figs. 1** and **2**.

## RESULTS

### A comparison of scRNA-seq methods

We selected seven scRNA-seq methods for comparison and tested each with up to three sample types: a mixture of mouse and human cell lines, frozen human PBMCs, and mouse cortex nuclei (**Fig. 1**, **Supplementary Figs. 1** and **2**). We chose to profile a cell line mixture with 50% human HEK293 and 50% mouse NIH3T3 cells (“mixture”) because (**1**) these cells are a common test^8, 9, 12, 24^ for samples with relatively high amounts of RNA per cell and (**2**) multiplets can be detected when cell barcodes have a substantial fraction of reads from both species. In mixture experiments, we grew HEK293 and NIH3T3 cell lines separately, mixed together equal number of cells for each cell line, and processed aliquots with seven scRNA-seq methods in parallel. We profiled frozen aliquots of human PBMCs because (**1**) they are a heterogeneous mixture of cells, particularly with respect to their amount of RNA per cell, yet they do not require dissociation (a separate technical challenge), (**2**) their cell types and associated expression patterns are well-studied. (We do not include data for sci-RNA-seq with PBMCs because we detected very few genes (e.g., <10 genes) per cell in several experiments (data not shown).) We wanted to extend our study of single cells to single nuclei as such samples have distinct properties including lower RNA input amounts. We profiled the mouse cortex because brain tissue is a major example of a tissue type commonly analyzed, by necessity, through single nucleus RNA-seq. We thus used four methods, which had been previously applied to nuclei^9, 12, 22, 25^, to test with nuclei from mouse cerebral cortex. Each sample type was tested in two experiments (Mixture1 and Mixture2, PBMC1 and PBMC2, Cortex1 and Cortex2) run on different days to assess reproducibility.

In each comparison experiment, we started with one sample with processing of aliquots starting at the same time for each method. The only exceptions were for Seq-Well in PBMC1, in which we thawed an identical PBMC aliquot a second time to obtain a Seq-Well dataset with sufficient cells profiled for PBMCs, and for 10x Chromium in PBMC1, in which we thawed an identical aliquot to directly compare version 2 (v2; designated as “B”) with version 3 (v3), which was not available when our original comparisons were done. In each experiment, we aimed to collect data from ∼384 cells for the low-throughput methods and ∼3,000 cells for the high-throughput methods. In each experiment, we also used an aliquot of cells to generate a bulk RNA-seq library as a control.

We sequenced all libraries together in an attempt to avoid batch effects due to varying sequence quality among Illumina flowcell lanes, with the following exceptions. The inDrops libraries were sequenced separately because they have an opposite read structure from those generated from the other methods with the cDNA in read 1 needing a long read and indexing information in read 2 needing a short read (**Online Methods**). We performed additional sequencing for some libraries in an attempt to sequence similar numbers of reads per cell for each low or high throughput method (**Online Methods**). We aimed for 50,000 to 100,000 reads per cell for high-throughput methods and 750,000 to 1,000,000 reads per cell for low-throughput methods.

### scumi computational pipeline allows unified analysis across any scRNA-seq method

Because each method had its own standard computational pipeline, we developed a new “universal” pipeline, to permit direct comparison of all the experimental methods, and remove processing differences introduced by these existing pipelines (**Supplementary Fig. 2**). First, we developed the scumi software package (<u>s</u>ingle-<u>c</u>ell RNA-sequencing with <u>UMI,</u> **Supplementary Fig. 2a**), which starts from FASTQ files as input and generates gene-cell expression count matrices for downstream analyses. We used scumi to analyze and compare the data from all the methods tested.

Second, we addressed the major pre-processing challenge of filtering out low-quality cells prior to downstream analysis (**Supplementary Fig. 2b**). This is particularly important when comparing methods, to ensure that our approach is fair to all methods and less subjective. In particular, when selecting the top cell barcodes with the largest number of reads or UMIs assigned to them, the challenge is to decide what threshold to choose for excluding lower quality cells or barcodes likely reflecting ambient RNA rather than real cells (or nuclei). For the mixture experiments we removed cells based on their complexity (UMIs or reads per cell). For the more complex PBMCs and cortex samples, consisting of different cells with different characteristics, such a simple approach could bias against the recovery of cell types with relatively small amounts of RNA. Instead, we first looked at more cell barcodes than we expected to truly recover from experiments, did an initial clustering, looked for differential gene expression to find cluster specific marker genes, and removed cells in clusters likely to be low quality (**Supplementary Fig. 2b**, **Online Methods**).

Third, prior to calculating metrics that potentially show improvements with greater sequencing depths, such as the number of genes per cell or ability to detect known cell types, we sampled the same number of reads per cell for all the methods of the same type, either low- or high-throughput, in a given experiment (**Online Methods, Supplementary Fig. 2c**). This leads to better relative performance for methods that have a higher fraction of informative reads, *i.e.* aligning to genes and present in cells used for analysis. Note that because for most experiments we sequenced the poly(T) sequences that follow the cell barcode and UMI sequences in all methods except SMART-seq2, we tracked and removed reads without poly(T) at the expected positions.

Finally, we assessed the methods by several key metrics spanning (**1**) the structure and alignment of reads to the nuclear and mitochondrial genomes; (**2**) sensitivity in capturing RNA molecules; (**3**) extent of multiplets (assessed in mixture experiments); (**4**) their technical precision/reproducibility with respect to expression estimates; and (**5**) the ability to recover meaningful biological distinctions in cell types (for the PBMC and cortex experiments). We describe each of these assessments next.

### Read structure and alignment reveal efficiency differences among methods

First, we characterized the methods by the distribution of reads from each library with respect to their structure and alignment with the genome (**Supplementary Fig. 3**). These metrics inform about the “efficiency” of methods in generating useful reads for downstream analysis. In the mixture experiments, we considered only uniquely mapped reads to minimize the effects of multi-mapped reads on calculating cell multiplet rates and other metrics.

The methods varied in the fraction of reads without poly(T) at the expected positions. Methods that do not use beads to capture mRNA (CEL-Seq2 and sci-RNA-seq) generally had poly(T) detected at the expected positions (**Supplementary Fig. 3**, **Supplementary Tables 1,2**). The 10x Chromium data also had a high fraction of reads with poly(T). By contrast, Drop-seq, Seq-Well, and inDrops showed high percentages of reads without poly(T). Interestingly, although Drop-seq and Seq-Well used the same batch of beads, the percentages of reads without poly(T) from the two methods were different, with 6.5% and 7.5% for Drop-seq, but 10.8% and 16.8% for Seq-Well in the mixture experiments. Similarly, in the PMBC experiments, 10x Chromium (v2 and v3) had the lowest fraction of reads lacking poly(T) (0.7% to 2.1%) and Seq-Well had the highest (41.2%; **Supplementary Table 2**, **Supplementary Fig. 3b**). These results suggest that some aspect of the methods independent of the beads contributed to the proportion of with reads without poly(T). In the nuclei experiments, both 10x Chromium (v2) and DroNc-Seq had a higher fraction of reads lacking poly(T) with nuclei than in the other experiments (**Supplementary Tables 1-3**, **Supplementary Fig. 3**).

We next considered the distributions of reads across these categories: exonic, intronic, intergenic, overlapping different genes (ambiguous), multi-mapped, and unmapped. Exonic reads are typically the only reads used in scRNA-seq studies of cells, whereas intronic reads are used for studies with nuclei. In the mixture experiment, both replicates of Smart-seq2 and one replicate of inDrops had the highest fraction of exonic reads (51.0%, 53.7%, and 56.9%, respectively), with sci-RNA-seq performing worst (28.7% and 29.4%, **Supplementary Fig. 3a**). Overall, the PBMC samples had a lower fraction of reads aligned to exons than the mixture samples (**Supplementary Fig. 3a**,**b**), with one replicate of inDrops having the highest fraction of exonic reads (46%), and Seq-Well having the lowest (20%, **Supplementary Fig. 3b**).

To determine the extent to which existing annotation limits recovery of reads aligning to genes^26^, we used the PBMC1 and PBMC2 bulk RNA-seq to create a matched transcriptome and new annotation (**Online Methods**). The new transcriptome annotations led to very few (<2%) additional reads aligning (**Supplementary Table 4**).

While the relative performance of each method was generally similar between the cortex nuclei and the other experiments, there was a higher ratio of intron-aligning reads to exon-aligning reads in nuclei than in cells (**Supplementary Fig. 3**) as expected because nuclei contain a higher proportion of unspliced transcripts than whole cells^27^. We assessed whether reads aligned in the sense or antisense orientation for each method, except Smart-seq2, which is not strand-specific. Interestingly, 10x Chromium (v2) had the highest fraction of antisense reads (33% and 29%), followed by DroNc-seq (10% and 12%), and sci-RNA-seq (6% and 9%; **Supplementary Fig. 3c**). We did not observe this level of antisense reads in bulk cortex nuclei nor in PBMC scRNA-seq datasets (data not shown). We excluded these antisense reads from our subsequent analysis.

### Fraction of mitochondrial reads varies, but is at or below level in bulk RNA-seq

We determined the fraction of reads aligning to mitochondrial genes, as these can be associated with lower quality scRNA-seq samples^28^ on the one hand, but can also be useful for inferring lineage relations between cells based on mtDNA mutations^29^ on the other hand.

In the mixture experiments, CEL-Seq2 had the highest frequency of such reads at 8.7% (**Supplementary Fig. 4a**,**b**), but this corresponds to the levels of mitochondrial transcripts in bulk RNA-seq for the cell line samples: 12.6% and 12.3% for NIH3T3 cells, and 8.5% and 9.1% for HEK293 cells, suggesting this may be an accurate reflection of the cells’ transcriptome. In PBMCs, inDrops and CEL-Seq2 had the highest mitochondrial ratios at ∼11% (**Supplementary Fig. 4b**). This may reflect the fact that both methods use *in vitro* transcription amplification rather than PCR. In addition, 10x Chromium (v3) had a high mitochondrial ratio at 11.2% (**Supplementary Fig. 4b**). Bulk RNA-seq of PBMC1 and PBMC2 had 12.6% and 12.3% mitochondrial transcripts, respectively, again suggesting this may be an accurate reflection of the transcriptome. In cortex nuclei, all methods had low mitochondrial ratios (**Supplementary Fig. 4c**), possibly because the isolation of nuclei successfully removed most mitochondria from the samples. This is consistent with the low (1.9% and 2.2%) fraction of mitochondrial transcripts in the matched bulk RNA-seq samples.

### Similar relative ranking of method sensitivity across experiments

As scRNA-seq methods start with limited RNA inputs, a key quality metric is the sensitivity, or the ability to capture RNA molecules. We assessed the sensitivity of each method by measuring the number of detected UMIs or genes per cell in datasets sampled to the same number of reads per cell (**Online Methods**; **Supplementary Table 5**). The only exception was Seq-Well PBMC1 with ∼46,000 reads per cell compared to ∼69,000 reads per cell for the other high-throughput methods in PBMC1 (**Supplementary Table 5**). For the mixture experiments, we report the results for mouse and human cells separately as the number of UMIs and genes per cell in the two cell types differs, such that differences in the ratio of human to mouse cells among the libraries (**Supplementary Table 1**) could skew the results, but the overall ranking of the methods is the same for both human and mouse cells (**Fig. 2a,b**, **Supplementary Fig. 5a**).

**Figure 2.**
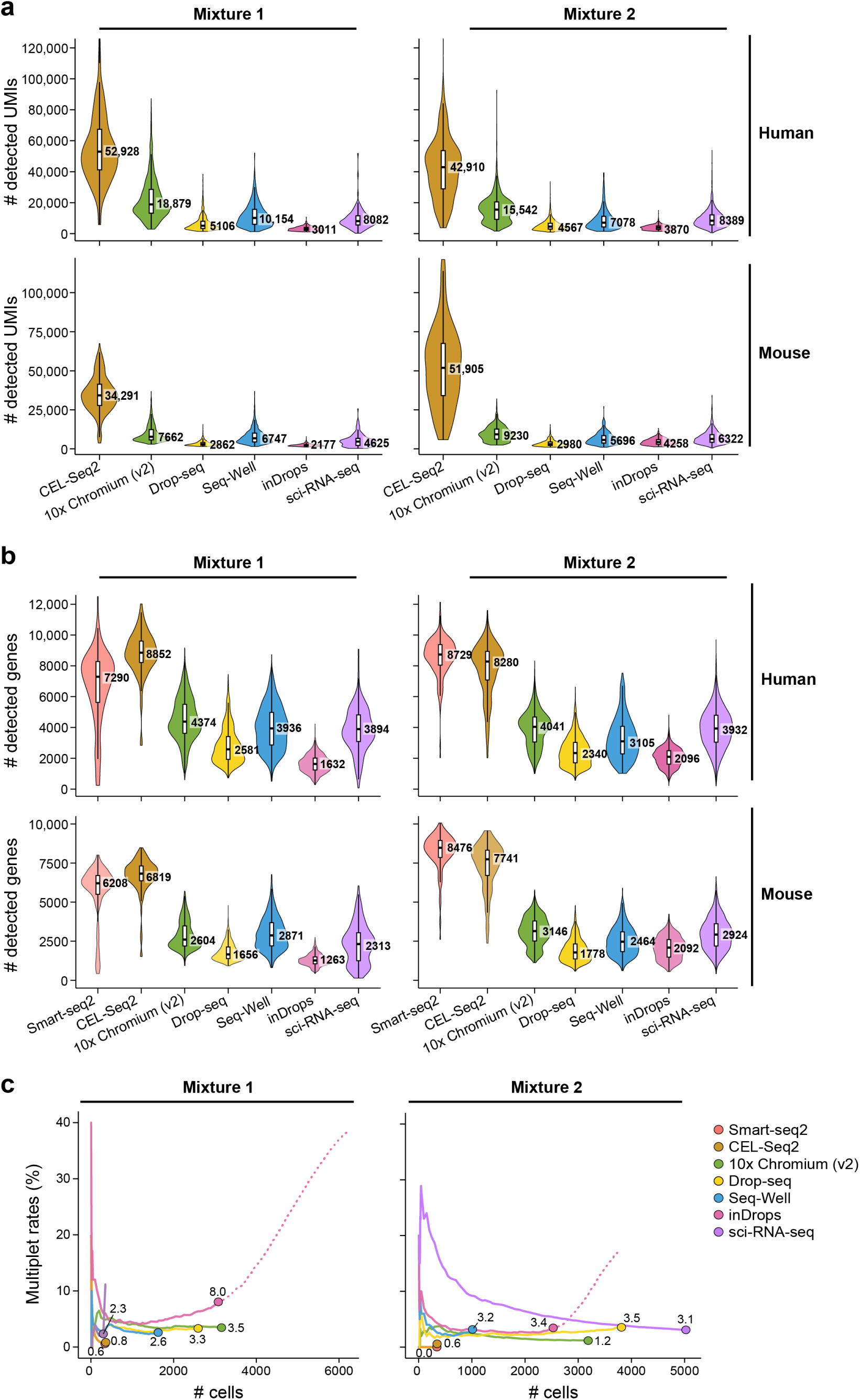
Performance metrics for mixture experiments. (**a**) UMIs per cell for each method in the two experiments. (**b**) Genes per cell for each method in the two experiments. For these plots, the top row shows data for human cells, and the bottom row data for mouse cells; the first column show data for replicate one, and the second column figures for replicate two. For (**a**) and (**b**), median and box plots were based on all the cells, but a few outlier cells were omitted in drawing the violin plots. Boxplots denote the medians (labeled on the right) and the interquartile ranges (IQRs). The whiskers of each boxplot are the lowest datum still within 1.5 IQR of the lower quartile and the highest datum still within 1.5 IQR of the upper quartile. (**c**) Multiplet frequency. We ordered cells based on the number of UMIs detected, from highest to lowest. For a given x value, the plot shows the percent of the top x cells (so cells with higher number of UMIs were further to the left on the *x*-axis) that are multiplets. The dotted line for inDrops shows the multiplet rate including low-quality cells that were not included in subsequent analysis.

Overall, low-throughput methods Smart-seq2 and CEL-Seq2 had the highest sensitivities, whereas among high-throughput methods, 10x Chromium detected the most UMIs and genes per cell. In the mixture experiments, inDrops had the lowest sensitivity and Seq-Well detected fewer genes per cell compared to 10x Chromium (v2) and sci-RNA-seq, but more genes per cell compared to Drop-seq and inDrops. The relative ranking of the methods was generally consistent when comparing the median number of detected UMIs per cell (**Fig. 2a**), detected genes per cell (**Fig. 2b**), or mean detected reads per cell (**Supplementary Fig. 5a**). Similarly, in PMBCs low-throughput methods detected more UMIs and genes per cell than the high-throughput methods. (**Fig. 3**, **Supplementary Fig. 5b**), with similar performance of Smart-seq2 (2406 and 2632 median number of genes detected) and CEL-Seq2 (2717 and 2545; **Fig. 3b**). Among the high-throughput methods, 10x Chromium (v3) had the largest median number of UMIs (4494) and genes per cell (1482; **Fig. 3**), and inDrops (366 and 1118 UMIs; 256 and 568 genes) and Seq-Well (844 and 577 UMIs; 513 and 372 genes) had the lowest (**Fig. 3**). In cortex nuclei, Smart-seq2 was the only low-throughput method tested and we sequenced to a slightly higher depth than for the other samples (**Supplementary Tables 1**-**3**) and used all the reads. As expected, Smart-seq2 detected more genes per cell than the high-throughput methods. (**Fig. 4**, **Supplementary Fig. 5c**).

**Figure 3.**
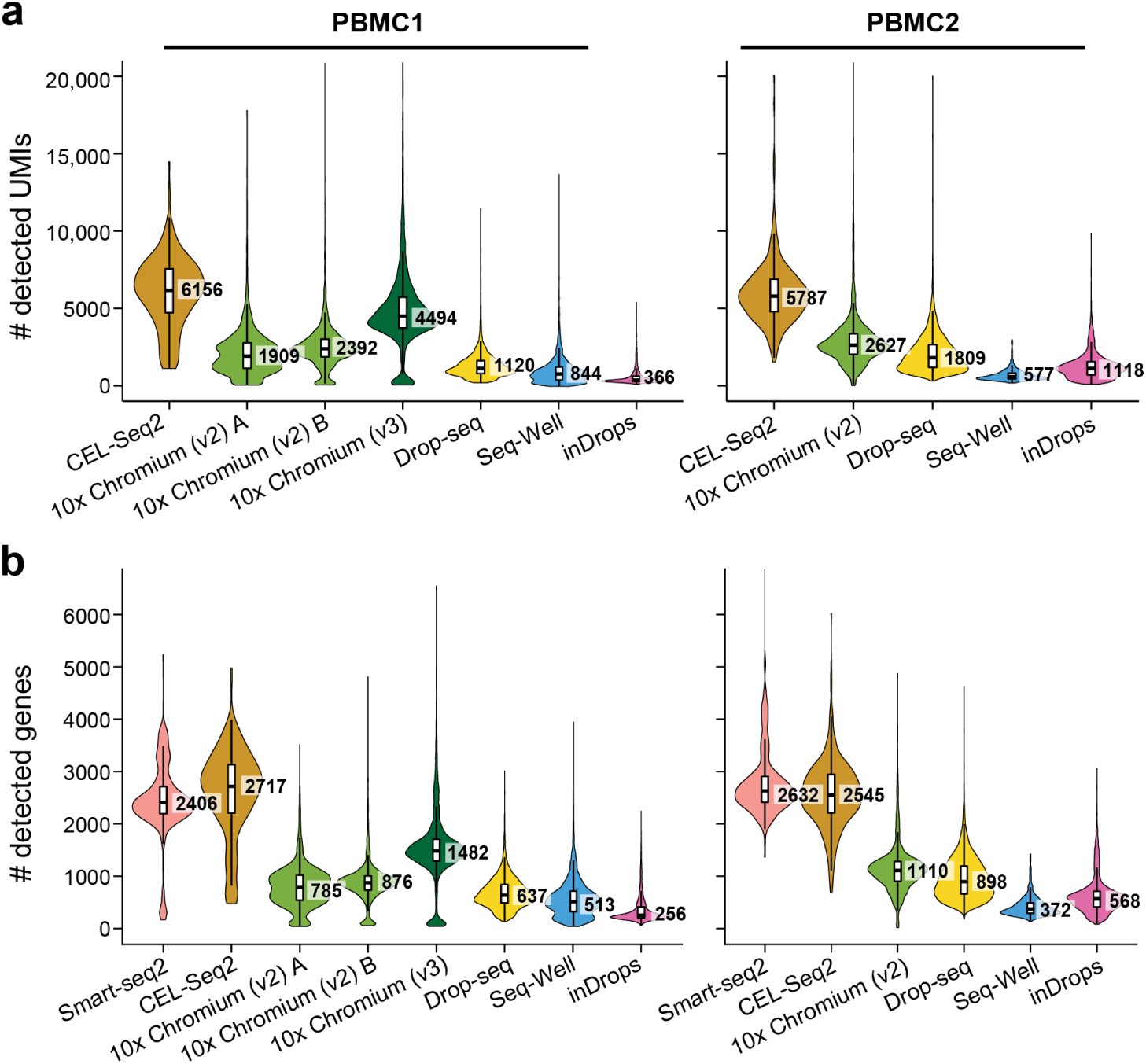
PBMCs sensitivity. (**a**) UMIs per cell for each method in the two experiments. (**b**) Genes per cell for each method in the two experiments.

**Figure 4.**
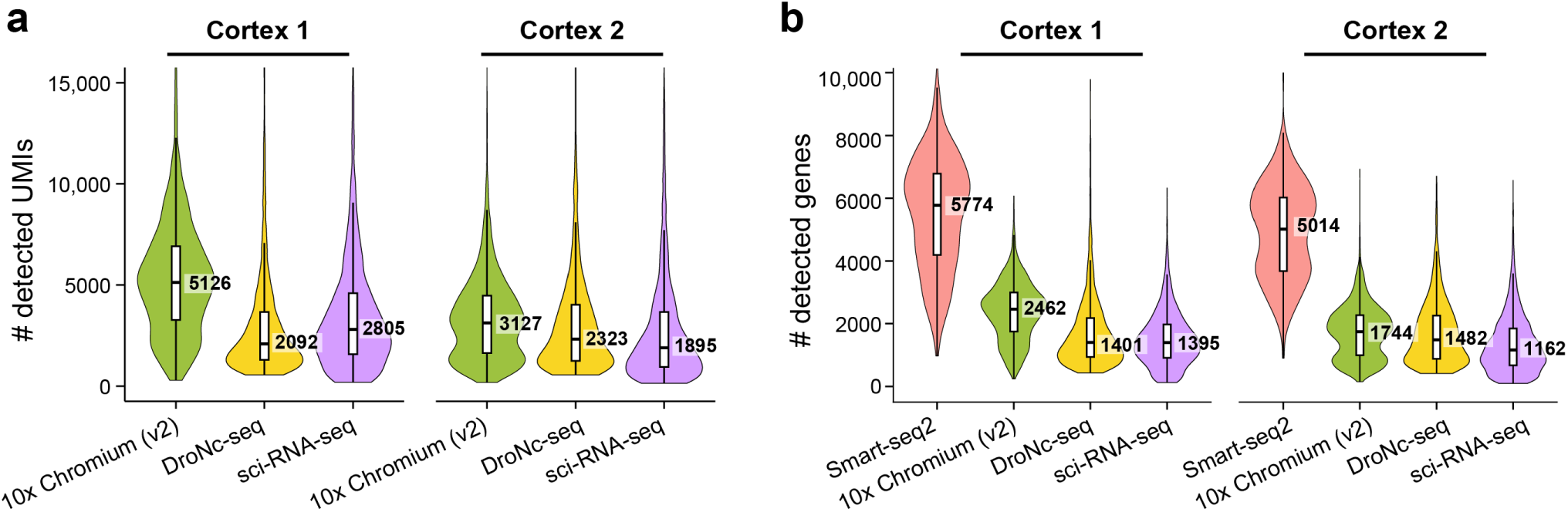
Cortex nuclei sensitivity. (**a**) UMIs per cell for each method in the two experiments. (**b**) Genes per cell for each method in the two experiments.

Among the high-throughput methods, 10x Chromium (v2) had the highest median number of UMIs (5126 and 3127) and genes per cell (2462 and 1744) (**Fig. 4**, **Supplementary Fig. 5c**).

Although it is not possible to directly compare these metrics to the mixture and PBMC experiments as each set of cells differ in many ways, including the amount of RNA per cell or nucleus, it is possible to compare the relative ranking of different methods. Although sci-RNA-seq had better sensitivity than Drop-seq with cell lines (**Fig. 2**), the two methods performed similarly in Cortex1 and DroNc-seq was more sensitive in Cortex2 (**Fig. 4**).

To explore how sensitivity varied with sequencing depth, we sampled fewer reads per cell from each method in the PBMC datasets. For each dataset, the relative ranking of the methods with respect to the median number of genes per cell detected remained the same at all sequencing depths (**Supplementary Fig. 6**). In addition, it appears that the number of genes detected may not have saturated at these sequencing depths, except for Seq-Well PBMC2.

### Mixture experiments enable detection of multiplets and reads from other cells

In the mixture experiment, we were able to assess the frequency of multiplets, two or more cells being sequenced together and assigned one cell barcode because we started with a mixture of human and mouse cells. The observed multiplet rates were less than 3.5% for all seven tested methods (**Fig. 2c**), except for the first inDrops experiment, which also had a high fraction of reads without poly(T) (**Supplementary Table 1**). The multiplet rate depends on the number of cells used in each experiment^9^ and the ratio of mouse to human cells, but it was not possible to sequence exactly the same number of cells nor the same ratio of mouse and human cells with each method in our experiments (**Supplemental Table 1**). The multiplet rates of low-throughput methods (Smart-seq2 and CEL-Seq2) were the lowest (<1%) of all the methods, as expected as FACS was used to place a single cell in each well of a plate (**Fig. 2c**).

We also examined how the estimated multiplet rate varied with the number of detected UMIs per cell. Generally, the multiplet rates were high in cells with the largest number of UMIs (**Fig. 2c**), as expected because multiplets are expected to have more RNA input. While most cells with intermediate quantities of UMIs were not multiplets, cells with lowest number of UMIs in some cases had higher rates suggesting that these cells might be low-quality or have more contributions from cell-free ambient RNA (**Fig. 2c**). In sci-RNA-seq experiment 2, the rate of multiplets decreased more gradually than for other methods for unknown reasons (**Fig. 2c**).

We also used the mixture experiments to ask whether the genes detected in a cell were actually from that cell instead of “contamination” from other cells. As sequencing depth increased, more genes were detected from the “wrong” species (**Supplementary Fig. 7a,b**), as reflected by the slope of a regression line along the cell barcodes adjacent to each axis (**Online Methods**), such that the best performing methods have the lowest slope. For the low-throughput methods, Smart-seq2 performed much better than CEL-Seq2 – perhaps related to the pooling of cells or their barcoding during library construction in CEL-Seq2. Among the high-throughput methods, inDrops had the lowest slope and Seq-Well had the highest slope.

### Reproducibility, technical precision, and accuracy in gene expression quantification

We compared the reproducibility of expression quantifications from replicates, among the different methods. In the mixture experiments, we calculated the Pearson correlation coefficients between pseudo-bulk data of each of the scRNA-seq mixture human and mouse datasets (**Online Methods**). As expected, replicates were almost always more correlated with each other than with pseudo-bulks from different methods (**Supplementary Fig. 8**). The pseudo-bulks from sci-RNA-seq, although correlated well with each other from the two replicates, were not as correlated with the pseudo-bulks from other methods. The pseudo-bulks from the other methods were highly correlated with each other (**Supplementary Fig. 8**), suggesting that all methods have high accuracies in quantifying the gene expression, consistent with previous comparison studies that demonstrated scRNA-seq methods had high accuracies based on measured ERCC spike-ins and their annotated molarities^16, 18^. Notably, pairs of methods with similar molecular biology – CEL-Seq2 with inDrops and Drop-seq with Seq-Well – were more highly correlated with each other (**Supplementary Fig. 8**).

To assess technical precision in the mixture experiment, which consisted of two homogenous cell lines grown in controlled conditions in culture, we also compared the variation in scRNA-seq data, which we expect to be primarily driven in this case by technical variation. Previous studies have demonstrated that the technical variation generally follows Poisson distributions^16, 30, 31^. CEL-Seq2, inDrops, and Drop-seq consistently had relatively low extra Poisson CVs (**Supplementary Fig. 9**). Consistent with previous findings, Smart-seq2 data had the highest extra Poisson CV, most likely because no UMIs were used (**Supplementary Fig. 9**).

To examine how well the scRNA-seq methods reflect gene expression captured in bulk experiments, we compared them in the PBMC datasets. For each library, we combined each of the single cell datasets, sampled to the same number of reads per cell, into a pseudo-bulk dataset and compared each of them to the matched bulk (**Online Methods**). Most libraries showed a strong correlation with the bulk with Smart-seq2, 10x Chromium, and inDrops having the highest correlations (**Supplementary Fig. 10**).

### Individual methods vary in their ability to distinguish and recover cell types

A key consideration for a scientist in choosing a scRNA-Seq experimental method is its ability to uncover the underlying biology of interest. Among the many biological features studied by scRNA-seq, one of the most prominent use cases is the identification and recovery of distinct cell types by clustering of scRNA-Seq profiles. Both the PBMC and mouse cortex datasets consist of diverse cell types, and were chosen to allow us to compare methods in the context of this use case. To this end, we processed the data with the goal of a fair and optimal assessment of each method. Not only did we sample the same number of reads per cell for each low- and high-throughput method in each experiment as above for the sensitivity metrics, we also performed another round of sampling to use the same number of *cells* from each low- and high-throughput method in each experiment (**Online Methods**). The only exceptions were Seq-Well PBMC2, which had fewer cells (**Supplementary Tables 2**, **5**) because we used only one microwell array for that experiment, while we used two arrays for Seq-Well PBMC1, and DroNc-seq Cortex2, which had fewer cells for unknown reasons (**Supplementary Tables 3**, **5**).

For each PBMC or mouse cortex dataset, we clustered the cells or nuclei based on their gene expression profiles to assess how well they detected the known cell types and their associated transcriptional profiles. For clustering, the Louvain algorithm’s performance largely depends on the parameter settings. For each dataset, we searched a range of parameters to select the optimal clustering to recover each of the expected cell types (**Online Methods**). For each cluster, we assigned a cell type identity based on known marker genes (**Online Methods**). Next, to quantify the quality of the clusters at separating cell types, we scored the expression of each cell for each cell type signature generated from known marker genes and calculated the area under the curve (AUC) for each cluster to measure how well the cells in a cluster score for each cell type (**Online Methods**). The AUC summarizes the performance of the gene signature scores in separating a cluster of cells from the rest of the cells, with AUC =1 for all cell types as the ideal outcome.

For PMBCs, methods varied in the ability to distinguish cell types, as well as in the proportion of cell types recovered and in some cases to recover certain cell types altogether. As expected, methods had more difficulty in distinguishing transcriptionally related cell types, such as CD4^+^ T cells, cytotoxic T cells, and natural killer (NK) cells (**Fig. 5a,b**, **Supplementary Fig. 11**). From the t-SNE plots for PBMC2, we observed that 10x Chromium and inDrops performed well (**Fig. 5a**, **Supplementary Fig. 11b**). Given that all the libraries for each experiment were generated from the same sample, we compared the fraction of cells assigned to each cell type within an experiment to see whether they were consistent (**Fig. 5b**). Generally, most methods were able to successfully recover the abundant cell types in PBMCs, i.e., CD4^+^ T helper cells, cytotoxic T cells, NK cells, B cells, and CD14^+^ monocytes, but varied in the relative abundance of cell types. Methods also varied in whether cell types were detected, particularly for the rarer cell types, such as megakaryocytes (**Fig. 5b**). For the low-throughput methods, we did not profile a sufficient number of cells to recover the rarer cell types (**Fig. 5b**). In PBMC1 among the high-throughput methods, 10x Chromium (v2) showed the best quality for both the number of cell types identified and the average AUCs across cell types, followed by Drop-seq and 10x Chromium (v3), with Seq-Well and inDrops not identifying two cell types (**Fig. 5c**). In PBMC2, 10x Chromium (v2) and inDrops performed well – identifying all the cell types (**Fig. 5c**). For Seq-Well PBMC2, the poor performance was strongly influenced by the low number of cells recovered in the experiment (**Fig. 5c**). The two low-throughput methods performed similarly for the AUC measurements and did not have sufficient cells to capture and identify clusters for the rarer cell types (**Fig. 5c**).

**Figure 5.**
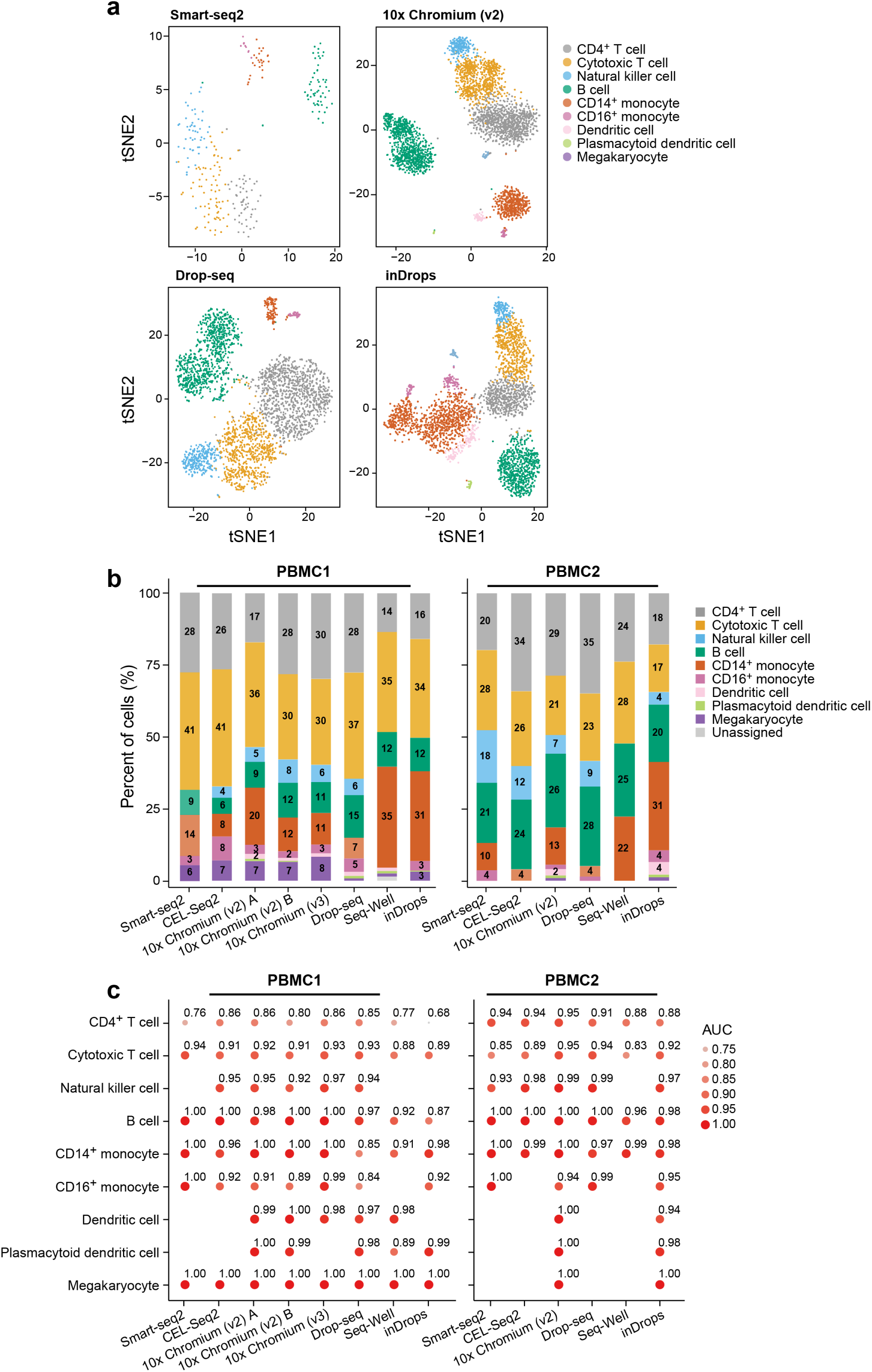
PBMCs biological metrics. (**a**) t-SNEs for representative PBMC2 libraries, (**b**) Proportion of each cell type detected with different methods. Those not labeled with a number rounded to one or less. Sum does not always add to 100 due to this and rounding. (**c**) AUC. The AUC of each cluster from classifying the cell type the cluster was assigned to. The size and color gradient of each dot encoded the AUC, which was also labeled beside each dot. The AUCs of cell clusters from PBMC1, and PBMC2. See **Supplementary Table 5** for the numbers of cells used.

Similar to PBMCs, the mouse cortex also has well-defined cell types, including excitatory and inhibitory neurons, astrocytes, oligodendrocytes, oligodendrocyte progenitor cells (OPCs), microglia, endothelial cells, and pericytes^32^. In both experiments for all the methods, apart from for sci-RNA-seq, we identified all these cell types, except pericytes, a rare cell type only found in DroNc-seq in Cortex1 (**Fig. 6**; **Supplementary Fig. 12**). In the sci-RNA-seq datasets, we also could not find OPCs and microglia (**Fig. 6**). In the AUC analysis, Smart-seq2, 10x Chromium (v2), and DroNc-seq all had high AUCs, though their relative ability to detect the expected cells varied by cell type (**Fig. 6c**). Notably, even the small number of cells in the Smart-seq2 datasets (295 and 349) sufficed to find these cell types, in contrast to the PBMC datasets (**Fig. 5**). There were some clusters of cells in the sci-RNA-seq datasets for which we could not confidently assign cell types (7% and 4% of cells; **Fig. 6a**,**b**). These cells were not used in calculating the AUCs.

**Figure 6.**
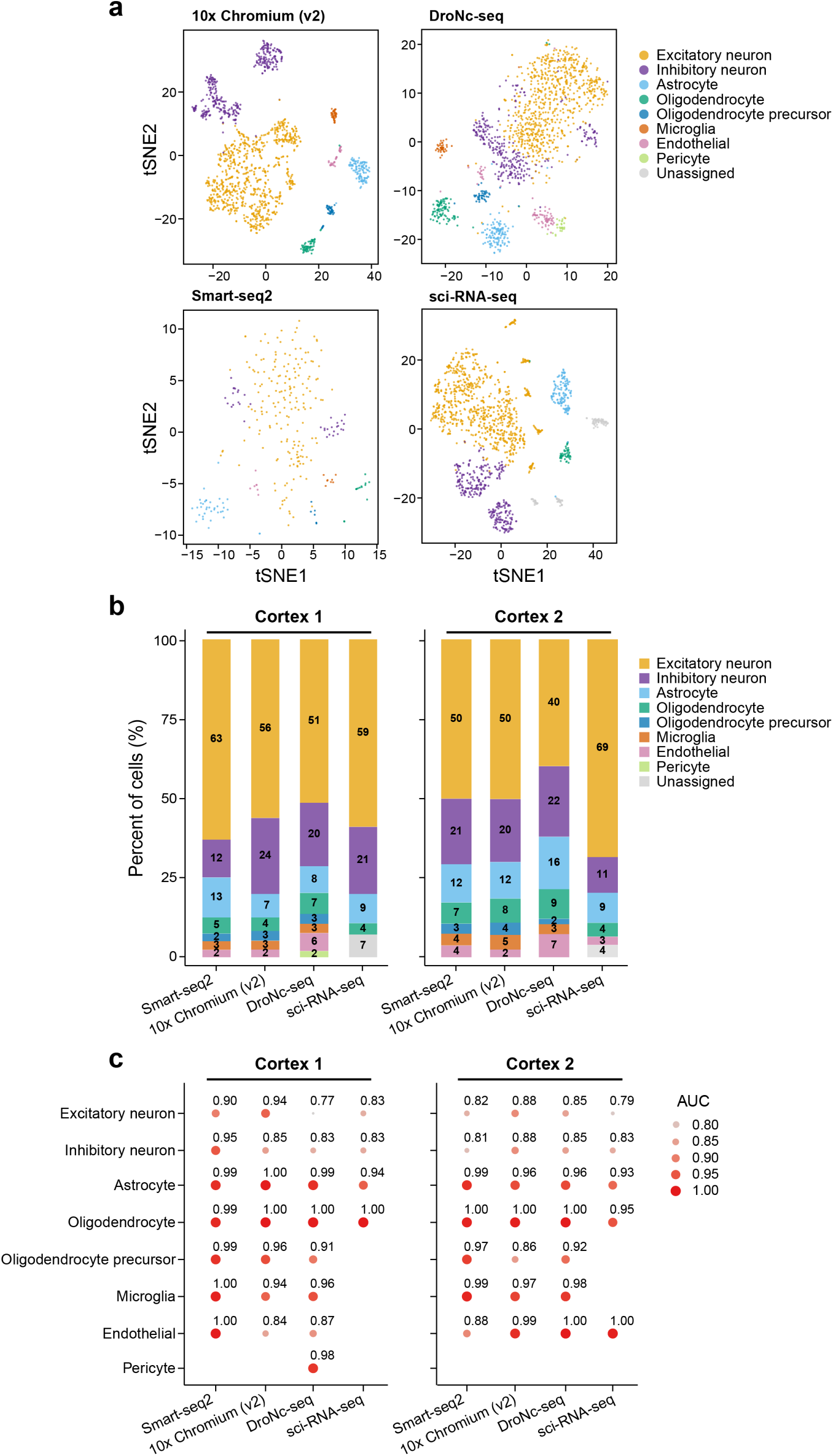
Cortex nuclei – biological metrics. (**a**) t-SNEs plots for Cortex1. (**b**) Proportion of each cell type detected with different methods. (**c**) AUC for Cortex1 and Cortex2. The AUC of each cell type in separating the cells from that cell type from the rest cells. The size and color gradient of each dot encoded the AUC, which was also labeled beside each dot. See **Supplementary Table 5** for the numbers of cells used. We could not confidently assign cell types to some clusters of cells from sci-RNA-seq and these cells were not used in calculating the AUCs.

### Pooled data analysis across methods enhances biological signal and consistency

Two general reasons may underlie the failure to detect certain cell types in the PBMC and cortex experiments: (1) libraries did not contain cDNAs from these cell types due to experimental issues; or (2) data quality from these cells may not have been insufficient to identify them, given this depth of sequencing and number of cells. To distinguish between these possibilities, we combined for each PBMC experiment all the sampled data together using Harmony^33^, re-clustered the cells (**Supplementary Fig. 13a**), and repeated our analysis. Following this analysis, all cell-types were detected in each library, supporting the second possibility and showing the power of accruing data across methods (**Supplementary Fig. 13b-d**). Moreover, we were able to determine in which cell type these missing cell types were originally assigned (**Supplementary Fig. 13c,d**). Although most of the combined and individual cell type assignments agree, some cell types seemed to be harder to distinguish. For example, in several libraries such as Smart-seq2 and CEL-Seq2, the undetected dendritic cells were grouped with the CD14^+^ or CD16^+^ monocytes (**Supplementary Fig. 13c**,**d**). Overall, 10x Chromium (v2) was the most consistent between the combined and individual level clustering, followed closely by 10x Chromium (v3), and others having fairly high but variable levels of consistency.

To check the cell types assigned by the combined analysis, we examined the cells assigned to cell types missing in our original analysis of each library separately. In most cases (20/25), we found that these cells could be assigned the same identity using our original AUC method (**Online Methods**), with some exceptions for rare cell types with only ∼1-2% of cells in a cluster (**Supplementary Table 6**). Thus, the failure to identify all the relevant cell types was due, as least in part, to data quality issues such as reads that could not be used in the analysis (**Supplemental Table 2**), with the possible exception of two rarer cell types in our datasets, megakaryocytes and plasmacytoid dendritic cells, which may not have been present in some datasets. Another possible explanation is that low quality cells were included that prevented identification of distinct cell types – this points to the difficulty in finding an optimal filtering threshold as well.

The combined analysis of mouse cortex nuclei (**Supplementary Fig. 14a**) allowed us to identify all the cell types in each dataset, except for pericytes, which clustered with endothelial cells (**Supplementary Fig. 14b**,**c**). In most cases, the combined and individual assignments agree (**Fig. 6**, **Supplementary Fig. 14c**). The 10x Chromium (v2) was the most consistent between the joint and individual level clustering, while sci-RNA-seq was the least (**Supplementary Fig. 14c**). In particular, some cell types were harder to distinguish—for example, oligodendrocytes and oligodendrocyte precursors, and microglia and excitatory neurons. All the cell clusters assigned new cell types by Harmony were confirmed when cells from individual libraries were checked with our AUC method (**Online Methods**).

### Comparison of scumi with standard computational pipelines

Although we developed and used the scumi computational pipeline in this study to analyze each method’s datasets in as similar a manner as possible, we also processed each of the datasets with its original method-specific pipeline for comparative purposes. There are significant differences in how the individual pipelines work. For example, the Drop-seq pipeline^8^ only considers primary alignments and discards secondary ones. For the reads that can be equally mapped to multiple positions, the primary alignment is typically randomly chosen from these alignments. On the other hand, Cell Ranger^9^ (for 10x data) only counts those multi-mapped reads if all the alignments of a read overlap a single gene.

Generally, there was good agreement between the basic metrics generated by the method-specific and scumi pipelines in terms of the median number of UMIs and genes per cell for the PBMC datasets (**Supplementary Table 2**). In particular, the relative ranking of all the methods for median genes per cell is the same between the scumi and method-specific computational pipelines (**Supplementary Table 2**). One of the most challenging steps in the analysis is choosing the correct number of cells in a dataset and most of these method-specific pipelines do not include this as a formal part of their pipeline or it does not work that well in practice, so that we followed standard practices and manually chose the number of cells based on the number of genes per cell in most cases (**Online Methods**). In addition, we examined the Pearson correlation of the gene expression for the two pipelines for each PBMC dataset. We found very high correlations for all datasets for both UMIs and genes (**Supplementary Fig. 15**). The genes per cell output agrees more closely than that of the UMIs per cell (**Supplementary Table 2**), perhaps because of the differences in how the method-specific pipelines process the data.

## DISCUSSION

In this study, we systematically benchmarked seven methods across three major categories: plate-based, bead-based, and combinatorial index-based methods. We selected representative methods that are more widely used and for which we had the expertise and resources to prepare the libraries. Our results with two replicates each for three different sample types – mixture of cell lines, human PBMCs and mouse cortex – were generally consistent in their ranking of the methods for sensitivity (**Figs. 2-4**), reproducibility (**Supplementary Fig. 8**), technical precision (**Supplementary Fig. 9**), and capturing biological information about cell types (**Figs. 5**, **6**, **Supplementary Figs. 13**, **14**).

All of the methods were able to generate useful data, but overall we found that 10x Chromium had the strongest consistent performance. In our limited testing of 10 Chromium (v3), which was introduced recently, we note that it had higher sensitivity (**Fig. 3**), but we did not detect improved cell type identification (**Fig. 5**) and had a higher fraction of reads aligned to mitochondrial genes (**Supplementary Fig. 4**). sci-RNA-seq, which has shown great scaling to larger numbers of single cells^13^, may require optimization for use with some samples, such as PBMCs. We note that we used the original version with two rounds of indexing^12^. Moreover, its performance with cortex nuclei was not ideal as it could not assign an identity to some cells and did not detect all the cell types present (**Fig. 6**, **Supplementary Fig. 14**). For the low-throughput methods, Smart-seq2 and CEL-Seq2 performed similarly without a consistent pattern for which was better (**Figs. 2-5**). For studies that require the highest sensitivity, these two methods are clearly better than the high-throughput methods (**Figs. 2-4**). Smart-seq2 has inherent advantages for genetic variant detection and studying RNA splicing isoforms because its sequencing is not limited to the 3’ end of genes – along with the disadvantage of lacking UMIs. Note however that in CEL-Seq2 we cannot rule out the issue of contaminating reads from other cells (**Supplementary Fig. 7**)^34^.

Looking beyond performance, we compared the time and reagent costs for each method as performed in this study (**Supplementary Table 7**). Drop-seq, inDrops, and Seq-Well had the lowest costs – noting that they are not fully packaged commercial products at this time – and Smart-seq2 was the most expensive, primarily because there is no pooling during library preparation. Many of the methods, particularly sci-RNA-seq, would be more cost effective with larger numbers of single cells or nuclei^13^. The 10x Chromium method required the least time and Smart-seq2, CEL-Seq2, and inDrops took the most time. We did not utilize automation, but it could decrease hands-on time and affect cost.

Analyzing single nuclei rather than single cells is an important strategy, which addresses tissues that cannot be readily dissociated into a single cell suspension (such as brain, skeletal muscle or adipose) and frozen samples, as well minimizes the alteration of gene expression which may be caused by dissociation^35, 36^. As in previous studies^37, 38^, we found that single nucleus RNA-seq generally performed well for sensitivity (**Fig. 4**) and classification of cell types (**Fig. 6**). Even with the inclusion of intron-aligning reads in our analysis, a higher fraction reads for 10x Chromium, and to a lesser extent for DroNc-seq, could not be analyzed because of the absence of a poly(T) sequence or aligning in an antisense orientation (**Supplementary Fig. 3**).

To systematically analyze and compare data generated from different scRNA-seq protocols and avoid introducing potential biases from method-specific computational pipelines, we developed the scumi software package. Although other scRNA-Seq processing pipelines^8, 9, 18, 39–46^ are available, some are specifically designed to process data from one or a few protocols^8, 9, 43–46^ and others can process data from different protocols, but are not scalable to large datasets^18, 40^ or are harder to use due to dependency and maintenance issues^42^. scumi is flexible to process reads from different protocols, works with different aligners, is scalable to billions of reads in a single run, has functions for detecting and correcting cell-barcodes, collapsing and filtering UMIs, and efficiently sampling reads to compare different methods, and, most importantly, is relatively easy to use. It takes FASTQ files as inputs and generates gene-cell expression count matrices. Except for removing the reads lacking poly(T) sequences, scumi keeps all other reads in BAM files that can be used for visualization or to pinpoint potential problems in the data. A significant challenge in scRNA-seq analysis, particularly when comparing across methods, is to decide which cells to exclude as low-quality. For samples such as PBMCs and cortex, heterogeneity in the RNA content of the different cell types makes a simple threshold of UMIs or genes per cell inappropriate as it introduces biases against cells with less RNA. Our pipeline provides multiple, customizable solutions to the problem of how to set threshold to filter out low-quality cells, including a mixture model or filtering of initially clustered datasets. Although scumi is not optimized for data generated from any one specific platform, we showed that it produced results that had high concordance with those from method-specific pipelines (**Supplementary Table 2**, **Supplementary Fig. 15**). We foresee that scumi could be useful for analyzing scRNA-seq data from new platforms and for future benchmarking studies of scRNA-seq methods.

Our study, the scumi pipeline, data and approaches will be a resource for future research in many fields where scRNA-seq methods are applied, and provides important guidance. First, using a coherently and reproducibly collected set of data, spanning three sample types, it provides direct guidance on key methods by a rich set of parameters and considerations – from technical to biological. It spans key and popular methods, including the first comparison of single nucleus RNA-seq methods. Second, the results presented here for each method could be used to further optimize and improve the existing scRNA-seq methods. Third, our use of representative and easily accessible sample types – common cell lines, human PBMCs, and mouse cortex from the C57BL/6 strain– should allow future studies, particularly those introducing new or improved methods, to make direct comparisons to this benchmark study. Indeed, all data sets were collected in a manner that allows open sharing, including of the human PBMC data. Finally, we expect that our datasets will be valuable for computational method developers to benchmark algorithms, e.g., batch correction algorithms, and build pipelines for efforts such as the Human Cell Atlas, the BRAIN Initiative, the Cancer Moonshot Human Tumor Atlas Network (HTAN), and other efforts to map cells in disease.

## ADDENDUM

This study complements a study entitled “Benchmarking Single-Cell RNA Sequencing Protocols for Cell Atlas Projects” by Mereu et al. which applied a complementary design (BIORXIV/2019/630087).

## ACKNOWLEDGEMENTS

We especially thank Mandovi Chatterjee, Alex Ratner, and Sarah A. Boswell of the Single Cell Core at Harvard Medical School for performing the inDrops experiments. We are grateful to A. Neumann, J. Lee, D. Dionne, and N. Sharif for assistance with project coordination, A. Klein for helpful discussions and suggestions, R. Kirchner for advice on inDrops data analysis, D. Leib for advice on CEL-Seq2 data analysis, B. Li for advice on PBMC data analysis, K. Shekhar for precision analysis in cell line mixture data, M. Cuoco for sample transportation, Broad Flow Cytometry Facility for cell sorting, Broad Genomics Platform for sequencing, and L. Gaffney for assistance with figures. Work was supported by the Klarman Cell Observatory, the Manton Foundation, and the BRAIN Initiative. A.R. is an Investigator of the Howard Hughes Medical Institute.

## AUTHOR CONTRIBUTIONS

J.L, A.S., O.R., and A.R. conceived the research. X.A., C.H., N.M., T.H., M.W., T.B., L.N., J.K., S.C., and S.L. performed the scRNA-seq experiments. X.A. and C.H. organized the sequencing. X.A. prepared the bulk RNA-seq libraries. J.D. created the scumi pipeline. J.D., S.S., A-C.V., A.K., and J.L. analyzed the data. M.K. contributed an optimized Smart-seq2 protocol. J.K. prepared the cell lines. A-C.V. prepared the PBMCs. B.B. prepared the mouse cortex. J.L., N.H. O.R., A.S., A-C.V., and A.R. provided supervisory guidance. J.D., X.A., S.S., C.H., T.H., M.W., T.B., J.K., A-C.V., A.R. and J.L. wrote the paper. All authors assisted in editing the paper.

## METHODS

### Cell preparation for the mixture experiment

We purchased human HEK293 (ATCC, #CRL-1573) and murine NIH3T3 (ATCC, #CRL-1658) cell lines and cultured them in Eagle’s Minimum Essential Medium (ATCC, #30-2003) supplemented with 10% Heat-Inactivated Fetal Bovine Serum (Thermo Fisher Scientific, #16140071) and in Dulbecco’s Modified Eagle’s Medium (ATCC, #30-2002) supplemented with 10% Iron Fortified Bovine Calf Serum (ATCC, #30-2030), respectively. We tested the cells for mycoplasma using a PCR-based assay (ATCC, #30-1012K) three days prior to each RNA-seq experiment. We maintained the cell lines at 37 °C, 5% CO_2,_ following the supplier’s recommendations.

We grew HEK293 and NIH3T3 cells to 50-60% confluence, detached them with TrypLE Express Enzyme (Thermo Fisher Scientific, #12604013), quenched them with complete growth medium, and spun them down at 250 x g for 5 min. We washed the cells twice with 1X Phosphate Buffered Saline (PBS), re-suspended them in 1X PBS containing 0.01% Bovine Serum Albumin (BSA, Sigma, #A8806), filtered them through a 40 µm cell strainer (Falcon, #21008-949), and counted them manually. We mixed the HEK293 and NIH3T3 cells at a 1:1 ratio, aliquoted cells for each scRNA-seq method, spun them down again and re-suspended them in a method-appropriate buffer (**Supplementary Table 8**). The passage number and time in culture for the cells are shown in **Supplementary Table 9**.

### PMBC preparation

All biospecimens were collected with informed consent by a commercial vendor. Use of all de-identified biospecimens for sequencing at the Broad Institute was further approved by the Broad’s Office of Research Subject Protection (ORSP), which determined that the research did not involve human subjects according to U.S. federal regulations (45CFR46.102f) – determination ORSP-3635. This study complied with all relevant ethical regulations.

We acquired commercially-available PBMCs (AllCells; Lot #PB006F-A5865) as frozen aliquots with 25 million cells for the PBMC1 experiment or isolated them from fresh blood (AllCells) within 4 h of collection using Ficoll-Paque density gradient centrifugation^47^ followed by cryopreservation (25 million cells/aliquot) using CryoStor CS10 (StemCell, #07930) for the PBMC2 experiment. On the day of each experiment, we rapidly thawed a PBMC aliquot and transferred the cells to a 50 ml Falcon tube containing 45 ml of RPMI medium without phenol (Thermo Fisher Scientific, #11835-055) containing 2% human AB serum (Corning, #35-060). We spun down the cells at 300 x g for 10 min at 4 °C and resuspended them in the same medium described above at a concentration of 1 x 10^8^ cells/ml prior to removing dead cells using the EasySep™ Dead Cell Removal (Annexin V) Kit (StemCell, #17899). We counted the live cells using trypan blue solution (Thermo Fisher Scientific, #15250061) and adjusted the cell concentration to 1000 cells/μl prior to distribution for downstream processes.

### Mouse whole cortex dissection

All animal-related work was performed under the guidelines of the Division of Comparative Medicine, with the protocol (# 0416-050-1) approved by the Committee for Animal Care of the Massachusetts Institute of Technology, and was consistent with the Guide for Care and Use of Laboratory Animals (1996 edition). Each cage contained 2–5 mice regardless of genotype; mice were housed at a constant 23 °C in a 12-h light–dark cycle (lights on at 7:00, lights off at 19:00) with ad libitum food and water. We obtained whole cortex samples by euthanizing 1 month old C57BL/6 male mice by cervical dislocation. We then dissected the whole cortex out of the first hemisphere by a midsagittal cut and a cut made between the cerebellum and the hemisphere. The hippocampus, olfactory bulb, and all basal ganglia were dissected out, and the cortex was placed in 500 µl RNAlater (Thermo Fisher Scientific) in a 1.5 ml tube, followed by incubation overnight at 4 °C before storage at −80 °C. Then second half of the cortex was processed similarly and placed in a different tube. We took only 2 min to harvest the first half of the cortex after the cervical dislocation step and another minute for the second half of the cortex.

### Single cell or nucleus experimental design

We performed two experiments with each single cell method for the mixed cell lines and PBMCs, except as noted. To generate data for Seq-Well, we performed a second PBMC1 experiment on a different day with an aliquot identical to the one used in the main PBMC1 experiment. Similarly, we performed a third PBMC1 experiment with 10x Chromium (v2) and (v3) on a different day. In addition, we performed two experiments with four methods for the mouse whole cortex nuclei. In all cases, each lab method was started at the same time by different researchers, so that the results would be directly comparable without any confounding due to the time cells or nuclei waited to start the experiment.

### Cell sorting

We passed each cell suspension through a 35 µm filter (Falcon) and added 20 µl of 1 µM TO-PRO®-3 Iodide (Thermo Fisher Scientific) per ml to stain dead cells. We used a MoFlo Astrios EQ cell sorter (Beckman Coulter) and set fluorescence activated cell sorting (FACS) gating on forward scatter plot, side scatter plot and on fluorescent channels to pick TO-PRO®-3 negative (live cells) while excluding debris and doublets. We used a 100 µm nozzle to sort single cells into 96-well plates containing 5 µl TCL buffer (Qiagen) with 1% beta-mercaptoethanol for Smart-seq2 and 384-well plates containing 0.6 µl 1% NP40 (Thermo Fisher Scientific) for CEL-Seq2.

### Nuclei isolation and sorting

We isolated single nuclei as previously described^23^ with the following modifications. We processed the left and right cortical hemispheres of one mouse separately through Dounce homogenization. We then combined the nuclei suspensions during the washing step. We aliquoted ⅓ volume in Tube A and ⅔ volume in Tube B. We spun down the aliquots and re-suspended the pellets in 1 ml DroNc-seq resuspension buffer (1X PBS, 0.01% BSA, 0.16 U/μl RNase Inhibitor (Clontech/TaKaRa, #2313A)) for Tube A and 1 ml 10X Nuclei resuspension buffer (1X PBS, 1% BSA, 0.2 U/μl RNase Inhibitor) for Tube B. We passed the nuclei suspensions through 20 µm strainers (Celltrics, Sysmex). We counted the nuclei and made the final aliquots for DroNc-seq and sci-RNA-seq from Tubes A and B at a concentration of 300,000 nuclei/ml with the DroNc-seq and 10x resuspension buffers (RBs), respectively. We preserved the rest of Tube A for later bulk RNA extraction and stained the rest of Tube B with Vybrant DyeCycle Violet Stain (Thermo Fisher Scientific) diluted to a final concentration of 10 µM.

We used FACS settings similar to those for single cells described above except for picking Violet positive (for nuclei) to sort 10,000 nuclei into 20 μl 10x RB buffer for 10x Chromium (10x Genomics). We also sorted single nuclei into 96-well plates as described above for Smart-seq2.

### Smart-seq2

We prepared RNA-seq libraries from single cells and single nuclei sorted into 96-well plates using a modified Smart-seq2 method^48^. Briefly, we cleaned up single cell RNA by adding 11 µl of RNAClean SPRI beads (Beckman Coulter Genomics) into each well of 5 µl of cell or nucleus lysate. After binding and washing twice with 80% ethanol, we eluted the RNA by adding 4 µl of master mix containing 1 µl of 10 mM dNTPs (New England Biolabs), 0.1 µl of 100 µM 3’ SMART reverse transcriptase (RT) oligo as 5’-AAGCAGTGGTATCAACGCAGAGTACT(30)VN, 0.1 µl of 40 U/µl RNase Inhibitor (Clontech/TaKaRa), 2.7 µl of 1 M Trehalose (Sigma), and 0.1 µl of 1/2500000x diluted ERCC ExFold RNA Spike-In Mixes (Thermo Fisher Scientific, #4456739). We incubated this mixture at 72 °C for 3 min for denaturing followed by a quick chill on ice and adding the remaining 6 µl reverse transcription mix that contained 2 µl of 5x First Strand Buffer (Thermo Fisher Scientific), 0.1 µl of 100 µM SMARTer II oligo (5’-AAGCAGTGGTATCAACGCAGAGTACATrGrG+G where “+” indicates a locked nucleic acid (LNA) base), 0.25 µl of 40 U/µl RNase Inhibitor, 0.1 µl of 200 U/µl Maxima H Minus Reverse Transcriptase (Thermo Fisher Scientific), 3.45 µl of 1 M Trehalose, 0.1 µl of 1 M MgCl_2_. We incubated the RT reaction at 50 °C for 90 min followed by 85 °C for 5 min. We then added 15 µl PCR master mix containing 12.5 µl KAPA HiFi HotStart ReadyMix (KAPA Biosystems)) and 0.5 µl 10 µM PCR primer (5’-AAGCAGTGGTATCAACGCAGAGT). We amplified for 21 PCR cycles with conditions as described^48^. We purified PCR products twice with 0.7x AMPure XP SPRI beads (Beckman Coulter Genomics) and eluted in 20 µl EB buffer (Qiagen). We quantified the cDNAs with the Quant-iT™ PicoGreen® dsDNA Assay Kit (Thermo Fisher Scientific) using an EnVision 2104 Multi-label Reader (Perkin Elmer). We normalized the cDNA concentrations and used 0.075 ng of each in a quarter volume of a NexteraXT (Illumina) library construction reaction. We pooled equal volumes from each well and purified with 0.6x AMPure XP SPRI beads.

### CEL-Seq2

We prepared CEL-Seq2 libraries from single cells sorted into 384-well plates following the published CEL-Seq2 method^6^ with these modifications. 1) To each RT reaction, we added 0.6 µl primer and dNTP mix that contained 4.28 ng RT primer and 1 nmol dNTPs. 2) We used 0.6 µl of NEBNext Second Strand Synthesis Enzyme Mix with 1x NEBNext Second Strand Synthesis Reaction Buffer (New England Biolabs) in a total volume of 12 µl for the 2^nd^ strand cDNA synthesis. 3) We pooled a total of 1152 µl (12 µl from each well) double stranded cDNA for each 96-well quadrant and used 0.8x volume of SPRI beads mix that contained 76.6 µl of the original RNAClean XP SPRI beads (Beckman Coulter Genomics) and 383 µl 2.5M Sodium Chloride, 20% PEG Solution (Teknova). 4) We used 2.75 µl 10X RNA Fragmentation Buffer (New England Biolabs) in a total 27.5 µl fragmentation reaction volume and later on added 2.75 µl 10x RNA Fragmentation Stop Solution. 5) We performed 15 cycles of PCR and used 0.8x volumes of AMPure XP SPRI beads for the first round of cleanup.

### 10x Chromium

We loaded 7000 single cells or nuclei onto a Chromium Single Cell 3’ Chip A (10x Genomics, PN-120236) and processed them through the Chromium Controller to generate GEMs (Gel Beads in Emulsion). We prepared RNA-Seq libraries with the Chromium Single Cell 3’ Library & Gel Bead Kit v2 (10x Genomics, PN-120237) following the manufacturer’s protocol.

In an additional experiment, we loaded one channel of PBMC1 cells side by side on Chromium™ Single Cell 3’ Chip A and Chip B (10x Genomics, PN-1000073) respectively aiming to capture 4000 cells each with Gel Bead Kit v2 and v3 (10x Genomics, PN-1000075) for comparison. We prepared the libraries following the manufacturer’s protocols.

### Drop-seq

We performed Drop-seq experiments with single cells according to the published paper^8^ with the following details. 1) We used a Drop-seq device that generated droplets 125 µm in diameter. 2) We used 100 cells/µl and 120 beads/µl. 3) We flowed cell and bead suspensions at 4 ml/hour, while flowing the oil at 16 ml/hour. 4) We collected the emulsion into a 50 ml Falcon tube for 15 min for each collection and did 2 collections for each experiment. We incubated the collections for up to 45 min at room temperature before breaking the emulsion. 5) We counted the total number of beads collected and made enough PCR reaction mix to have 5000 beads per 50 µl PCR mix in each well. 6) We performed 10 cycles of PCR during the second round of amplification as 98 °C for 20 sec, 67 °C for 20 sec, and 72 °C for 3 min.

### DroNc-seq

We performed the DroNc-seq experiment with single nuclei according to the published paper^23^ as described for Drop-seq (see above) with the following additional details. 1) We used a DroNc-seq device that generated droplets 75 µm in diameter. 2) We size selected beads <40 μm in diameter using a strainer (PluriSelect, #43-50040-03). 3) We used 300 nuclei/µl and 350 beads/µl. 4) We flowed cell and bead suspensions at 1.5 ml/hour, while flowing the oil at 16 ml/hour. 5) We collected the emulsion into a 50 ml Falcon tube for 22 min for each collection and did 2 collections for each experiment. We incubated the collections for up to 45 min at room temperature before breaking the emulsion.

### Seq-Well

We prepared the Seq-Well libraries following the published paper^10^ with the following details. 1) We loaded 12,000 cells onto each array to capture 2000 to 3000 cells. 2) We sealed the array with a hydroxylated polycarbonate membrane with pore sizes of 0.01 μm to allow buffer exchange while retaining biological molecules. 3) We used 1500 - 2000 beads per 50 μl PCR reaction volume and pooled 8 PCR products for each final library construction. 4) We purified PCR products with 0.6x AMPure XP SPRI beads followed by another 0.8x SPRI beads clean-up. 5) We prepared Illumina sequencing libraries with 800 pg of pooled cDNA PCR product from 12,000 - 16,000 single-cell transcriptomes attached to microparticles (STAMPs) and purified with 0.6x AMPure XP SPRI beads followed by another 0.8x SPRI beads clean-up.

### sci-RNA-Seq

For the mixed cell experiments, we started with 1 million cells following the published paper^12^ with these modifications and details. 1) We added RNase Inhibitor (Clontech/TaKaRa, #2313B) to all wash buffers in place of DEPC and SUPERaseIn RNase Inhibitor (Ambion). 2) We preloaded a 384-well plate with 1.25 μl of the mix containing 20 μM oligo-dT primer and 2.5 mM dNTP in each well. 3) We counted the fixed cells, re-suspended them at 500 cells/μl, and aliquoted 1000 cells into each well of a 384-well plate. 4) We used RNase Inhibitor in place of RNaseOUT Recombinant Ribonuclease Inhibitor (Invitrogen) in the first-strand RT reaction. 5) We pooled the cells after RT, then spun them down and re-suspended them in 1 ml PBS buffer containing 1% RNase Inhibitor and 1% BSA prior to staining with NucBlue Fixed Cell ReadyProbes Reagent (Invitrogen) at 2 drops/ml. 6) We sorted 40 cells into each well containing 5 μl TE buffer containing 10mM Tris pH 8.0 and 1 mM EDTA in 96-well plates and gated based on forward scatter plot, side scatter plot, and DAPI signal to avoid doublets. 7) For PCR, we used 1 μl each of the 10 μM P5 and P7 primers and amplified for 22 PCR cycles. 8) We did a two-sided size selection to remove the large genomic DNA (gDNA) fragments by adding 0.5x volume AMPure XP SPRI beads to the PCR products and recovered the supernatant after incubation followed by purifying the supernatant with another 0.5x SPRI clean up.

For the single nucleus experiment, we followed the published paper^12^ with similar details as for the mixed cell experiment with these additional changes. 1) We eliminated the methanol fixation step. 2) We used a final concentration of 300 nuclei/μl and aliquoted 600 nuclei/well across one 384-well plate. 3) We pooled the nuclei after RT and filtered through a 35 µm strainer (Falcon) prior to staining and sorting. 4) We gel-purified the final libraries after PCR to efficiently remove the large quantity of gDNA fragments following a procedure similar to a previously published paper^49^. Briefly, we ran the samples on a 10% Criterion TBE gel (Bio-Rad) at 100 V for 65-70 min prior to staining with 1/1000x SYBR Green (Thermo Fisher Scientific) for 10-20 min. We cut out the region from 300 to 600 bp from the gel, collected it into a 1.5 ml Eppendorf tube, crushed it with a disposable pestle (Kontes/Kimble Chase, #749520-0090), added 400 μl of 0.3M NaCl to the crushed gel pieces and incubated with rotation at room temperature for at least 4 h to overnight. We used a Spin-X cellulose acetate filter (Corning, #8163) to remove the gel particles by spinning for 3 min at 15,000 x g. We ethanol precipitated the flow through by adding 1 μl glycogen (20 μg/μl; Roche, #10901393001) and 2.5 volumes 100% ice cold ethanol followed by incubation at - 20 °C for at least 30 min and then 20 min of centrifugation at 15,000 x g and 4 °C. We washed the pellets with 1 ml 70% ethanol and air dried before resuspending the libraries in EB buffer (Qiagen).

### inDrops

We prepared inDrops libraries based on a previously described protocol^7, 50^ at the Single Cell Core at Harvard Medical School with 2000 to 2500 cells collected for each library. We made the following modifications in the primer sequences to eliminate the need for custom sequencing primers. 1) We changed the sequences of RT primers on hydrogel beads to: 5’-CGATTGATCAACGTAATACGACTCACTATAGGGTGTCGGGTGCAG[barcode1,8nt]GTC TCGTGGGCTCGGAGATGTGTATAAGAGACAG[barcode2,8nt]NNNNNNTTTTTTTTTTTT TTTTTTTV-3’. 2) We changed the PE2-N6 primer sequence in step 151 of the published protocol^50^ to: 5’-TCGTCGGCAGCGTCAGATGTGTATAAGAGACAGNNNNNN-3’. 3) We moved the indexing barcode from PE1 primer to PE2 in steps 157 and 160 of the protocol^50^ as the following: PE1-5’-CAAGCAGAAGACGGCATACGAGATGGGTGTCGGGTGCAG-3’, PE2-5’-AATGATACGGCGACCACCGAGATCTACACXXXXXXXXTCGTCGGCAGCGTC-3’, where XXXXXX is the indexing barcodes for multiplexing libraries.

### RNA-Seq from bulk samples

We pelleted the cultured cells, PBMCs, and isolated nuclei leftover from Tube A after taking out the aliquot for DroNc-seq (see Nuclei isolation and sorting section) by spinning at 500 x g for 5 min at 4 °C. After removing the supernatant, we resuspended them in 100 µl DNA/RNA Shield (Zymo Research) and stored them at −80 °C. We extracted total RNA using Quick-RNA Mini Prep (Zymo Research) and followed the vendor protocol with a DNase treatment step. We prepared the RNA-seq libraries using the TruSeq® Stranded mRNA Library Prep Kit (Illumina) following the manufacturer’s protocol except for the following modifications. 1) We did the elution, priming and fragmentation of mRNA at 85 °C for 4 min. 2) We used SuperScript III (Thermo Fisher Scientific) in place of SuperScript II and performed reverse transcription at 55 °C. 3) We used a different set of barcoding indices rather than those in the TruSeq kit for the ligation and final PCR steps.

### Sequencing

For each comparison experiment, we pooled together the single cell or nucleus libraries from different methods with normalized molar concentrations and loaded them onto a HiSeq2500 (Illumina) in the 8-lane high output or 2-lane rapid run mode to minimize the potential sequencing variation. For analysis, we trimmed the barcode reads (including inline barcodes from Read1 or Read2 and also the index read) back to the original length required for each library type, since the same run parameters had to be used throughout the sequencing run. Additional details are in **Supplementary Tables 10** and **11**. We also performed additional sequencing aiming to generate a total of 10^6^ reads per cell (or nucleus) for the plate-based Smart-seq2 and CEL-Seq2 methods, and 10^5^ reads per cell (or nucleus) for the other methods. We sequenced the bulk RNA-seq libraries on a NextSeq500 (Illumina) with 75 bases each for reads 1 and 2 and 8 bases each for index reads 1 and 2.

### Read demultiplexing

As methods using a single index read (e.g., 10x Chromium) or dual index reads (e.g., Smart-seq2) were pooled together for sequencing, we adopted a two-round demultiplexing strategy. We first demultiplexed the reads using both index reads to extract reads from Smart-seq2 and sci-RNA-seq. In the next round, we further demultiplexed the reads using a single index read. In the case of barcode collisions (i.e., a single index library index read matches that of a dual index library), we extracted the single index library reads from the unmatched reads after the first demultiplexing step. We only kept reads with one or no mismatches in the index reads.

### Annotating each cDNA read with its cell barcode and UMI

Different scRNA-seq protocols produce reads with different structures, especially the reads consisting of cell barcodes and/or UMI sequences. To address this issue, we used our scumi pipeline, which uses regular expressions (text strings defining search patterns) to extract the cell barcodes and UMI sequences from different FASTQ files and put them in the header of their corresponding cDNA reads. We started with the FASTQ files generated after de-multiplexing the BCL files from Illumina sequencers. For a typical 3’-tag based scRNA-seq experiment, a cDNA sequence fragment is in one read of a FASTQ file, and its corresponding cell barcode sequences and UMI sequences are in a paired read of a separate FASTQ file. For example, for the reads generated from the Drop-seq platform^8^, the cDNA reads are in read 2 and the cell barcodes (base 1 to base 12) and UMIs (base 13 to base 20) are in read 1. Details about the location of cell barcodes and UMIs can be found in **Supplementary Table 11**.

The scumi pipeline also corrects for sequence errors in the indices used by bead-based methods. We have implemented code similar to the standard Drop-seq pipeline that overcomes the problems observed for some batches of Drop-seq beads, in which up to 20% of the cell barcodes have errors in the last base (base 12), mostly because the beads (encoding the cell barcodes) only synthesized 11 (or fewer) bases^8^. In such cases, base 12 of the cell barcode is actually the first base of the UMI sequence, and the last base of the UMI sequence is from the poly(T) sequence. This bead synthesizing error can be detected by calculating the frequencies of T bases in the UMI sequences. The scumi pipeline first detects possible erroneous cell barcodes and then merges these cell barcodes that are the same in their first 11 bases but differ in the 12th base. If more than one base of the UMI sequences (with the same cell barcode) had a high-frequency of T bases (more than 80%), these cell barcodes were removed from further analyses.

### Mapping reads to a reference genome

We aligned the merged FASTQ files (each cDNA read with its cell barcode and UMI annotations) to a reference genome using STAR^51^ v 2.6.1a, except for Smart-seq2 and CEL-Seq2. For those libraries, we used HISAT2^52^ v2.0.5 as it is better suited than STAR to paired end read data such as Smart-seq2 because of the way it handles read pairs that do not both align to the same region of the genome – leading to more aligned reads and more detected genes per cell. Notably, HCA has adopted this aligner for its Smart-seq2 pipeline (https://staging.data.humancellatlas.org/learn/userguides/data-processing-pipelines/smart-seq2-workflow/). We also used it for CEL-Seq2 to facilitate better performance comparisons between the two low throughput methods. For mixture data, we used the STAR reference available in the hg19 and mm10 v2.1.0 Cell Ranger reference. For PBMC data, we used the STAR reference available in the GRCh38 v1.2.0 Cell Ranger reference. For cortex data, we used the STAR reference available in the mm10 v1.2.0 Cell Ranger reference. We downloaded Cell Ranger reference data from https://support.10xgenomics.com/genome-exome/software/pipelines/latest/advanced/references. For each sample type, we also generated a HISAT2 reference with the associated GTF and FASTA files.

### Annotating each alignment with a gene tag

We use featureCounts^53^ from the Subread package, v1.6.2, to add a gene tag to each alignment. To count reads overlapping with introns for single nucleus RNA-seq data, we used a two-step approach to first count the reads overlapping with exons. In the second step, the reads not overlapping with exons were recounted if they overlapped with introns. We only included reads aligning in the sense orientation with the genome annotation, except for Smart-seq2, which does not generate strand-specific data.

### Counting transcripts of each gene in each cell

For the UMI-based methods, we used scumi to generate a cell x gene UMI count matrix. We included a multi-mapped read if all its alignments overlapped with a single gene, similar to the Cell Ranger pipeline^9^. We collapsed UMIs in reads from the same gene from the same cell based on a Hamming distance of one. To prevent over-collapsing UMIs^42^, we did not collapse two UMIs – in the same gene in the same cell – if they each had more than five reads support. For Smart-seq2, we used a similar procedure to generate the count matrix used for the sensitivity and technical precision metrics, except we created a cell x gene read count matrix. For clustering Smart-seq2 data and downstream analysis, we used RSEM^54^ v1.3.0 to generate a cell x gene transcripts per million (TPM) matrix, which was used instead of the UMI count matrix. We generated the RSEM reference using the FASTA and GTF files used for creating the STAR and HISAT2 references (see Mapping reads to a reference genome). When generating the RSEM reference for cortex data, we modified the GTF to include one unspliced transcript per gene that included all introns and exons in that gene. This allowed us to count reads that mapped to introns.

### Selecting the number of cells

For a scRNA-seq experiment, we have a rough estimate of the number of cells *N* that can be recovered. A simple yet robust empirical method (used by Cell Ranger of the 10x Chromium pipeline) for cell barcode selection is to first estimate the library size *m* (in reads or UMIs) by the 99th percentile of the top *N* cell barcodes in terms of the number of reads (or UMIs). The cell barcodes with reads (or UMIs) greater than 0.1*m* are considered as ‘cells’.

For the cell line mixture experiments, we used different filtering approaches depending on the dataset. For 10x Chromium, Drop-seq, Seq-Well, inDrops, and sci-RNA-seq, we used this empirical rule for cell barcode selection. For Smart-seq2 and CEL-Seq2, we had a better estimation of the number of cells as we sorted individual cells into wells. We used a mixture of two Student’s *t* distribution model^55^ to model the read or UMI (log_10_ transformed) count distributions of each cell, and removed the cells that were likely from the mixture component with fewer reads or UMIs (posterior probability ≥0.5). The parameters of the Student’s *t* mixture model were estimated by maximizing the posterior distribution using the Expectation-Maximization algorithm. For sci-RNA-seq and inDrops, the empirical rule tended to select low-quality cell barcodes. We therefore used this mixture model on the cell barcodes selected by the empirical rule to further filter out likely low-quality cells.

For all the high-throughput PBMC datasets, we extracted two times the number of expected cell barcodes for each method, choosing the cells with the most reads. We removed cells with a high fraction of reads aligning to mitochondrial genes (names starting with ‘mt-’ for mouse and ‘MT-’ for human) – greater than 75th percentile + 3 * IQR of the mitochondrial ratios across the top returned cell barcodes, where IQR stands for interquartile range. For each cell, its UMI counts were divided by the total number of UMIs from that cell and then scaled by multiplying 10,000 to get transcripts per 10,000 (TP10K). We then added 1 to these TP10K and log transformed by the natural log. We then performed PCA using all genes, did clustering analysis (Louvain clustering^56, 57^ of the *k*-nearest neighbor (*k*-NN) graph built from the first 50 principal components of each single-cell dataset with parameter *k*=30 and a resolution parameter used for Louvain optimization of 1.0, implemented in the Seurat package^58^ v2.3.4 (see Parameter selection for clustering analysis section for more details), followed by differential gene expression analysis with the FindAllMarkers command in Seurat to find cluster specific (up-regulated) marker genes. To filter out clusters of cells likely derived from low quality cells or empty droplets, we removed clusters with insufficient markers genes, as follows. First, we identified marker genes for each cluster as genes expressed in ≥25% of cells in that cluster and with FDR <0.01 (significantly highly expressed in the cluster compared to the cells not in that cluster). Second, we excluded ribosomal protein coding genes, MALAT1, and genes starting with MTRNR, as they could be erroneously identified as marker genes after normalization and scaling because for cells with a small number of UMIs, the UMIs of highly expressed genes will be weighted more than those from cells with a large number of UMIs based on the scaling formula x_j_/sum_j_ x_j_, where x_j_ is the expression of gene j of a cell. Third, we only kept the clusters in which > 70% of the top 15 marker genes (or 10 out of all markers genes in a cluster that had <15 marker genes) were not mitochondrial protein coding genes as high expression of mitochondrial genes can indicate stressed cells or empty droplets. This process was repeated twice.

We used a modified strategy for Smart-seq2 and CEL-Seq2 because there were fewer cells to cluster, which potentially could have led to the low-quality cells not forming distinct clusters. The assumptions for these cell selections were that (1) there were enough low quality cells to form distinct clusters and (2) the clustering algorithm did not split high-quality cells of the same cell type into many distinct clusters because this could have led to some sub-clusters having too few marker genes. We therefore set *k*=5 (the number of neighbors in building the *k*-NN graph) to detect small clusters with a low solution parameter of 0.5 to prevent splitting cells of the same type into many clusters. We also only used the top 25 principal components as we did not expect to identify as many cell types from a smaller number of cells.

For each cortex dataset, we observed that the total numbers of UMIs from individual cell barcodes returned by scumi followed a bimodal distribution. We therefore first used a mixture of two Student’s *t* distribution models to filter nuclei with few UMIs (with probability ≥0.5 likelihood for the mixture component with relatively few UMIs compared to the cell barcodes from the other mixture component) and then applied the filtering approach used for PBMC data. We used the top 25 principal components and set the number of nearest neighbors *k* to 10 in clustering analyses (the choice of parameters was to help recover rarer cell types with lower numbers of UMI per cell, such as the oligodendrocyte precursors). To prevent splitting up big clusters due to the small *k*, we lowered the resolution to 0.8. For these nuclei libraries, the mitochondrial ratios were very low compared to those from cells, so that it was less likely to see low quality nuclei with mitochondrial protein coding genes as the top marker genes. Therefore, in addition to using mitochondrial protein coding genes to remove poor quality nuclei, we removed cluster-specific marker genes that were also expressed in >70% of the nuclei from the other clusters in a given library. We also used the top 20 marker genes in each cluster for filtering. For Smart-seq2 data, we only used the mixture model for cell selection as the remaining clusters could be assigned known mouse cortex cell types and applying the cluster-based filtering removed many cells that could be easily identified as known cell types.

### Multiplet Detection

In the mixture experiments, we considered cells multiplets if they had >15% of UMIs from mouse and >15% of UMIs from human. For Smart-seq2, this calculation was instead based on the percent reads, though with the same cutoffs. We report the observed multiplet rate, though the actual multiplet rate was likely to be about twice this frequency due to the presence of undetected multiplets that are all from cells of the same species (assuming about equal numbers of human and mouse cells).

### Detection of both human and mouse genes expressed in single cells

In the Mixture experiments, for each human cell, we computed the percentage of detected mouse genes in that cell. Similarly, we repeated this analysis to detect human genes from mouse cells. To show the relationship between sequencing depth and the number of detected genes from the other species, we drew scatter plots for the detected human cells and mouse cells where the *x*-axis is the number of detected human genes and the *y*-axis is the number of detected mouse genes. Generally, the number of detected genes *y* from the other species increases linearly with the number of detected genes *x* from the called species. Therefore, we used robust linear regression (Huber M-estimator, implemented in the MASS R package) to fit *y* = a*x* + b, for human cells and mouse cells, independently. The fitted line and its slope (a) were plotted on the scatter plots.

### Pseudo-bulk expression profile comparisons

We built pseudo-bulks by combining the expression vectors from all the human cells (or mouse cells) in the mixture experiments and computed the Pearson correlation coefficients between the pseudo-bulks from different methods. The pseudo-bulk gene expression profile for a set of cells was computed by first taking the sum of the gene expression vectors from these cells, where the gene expression vector of a cell was the raw UMI count vector, one element for a gene. The dimensionality of a gene expression vector was the number of genes. To make the bulk gene expression vectors from different sets of cells comparable, a bulk gene expression vector was normalized by dividing the total number of UMIs from all the cells used in computing that bulk gene expression vector, and further multiplying by 10^4^ and finally taking the natural log transform (adding one before the log transformation to make all the elements of the bulk vector positive). For Smart-seq2 data, we summed the read counts from cells and then transformed the counts to transcripts per 10K (TP10K) values (10^4^ (N_i_ / L_i_) / sum_i_ (N_i_ / L_i_), where N_i_ is the total number of counts for gene *i* and L_i_ is the length of gene *i*) to get the pseudo-bulk expression.

### Bulk RNA-seq processing and analysis

Expression in bulk RNA-seq data was quantified with RSEM^54^. We extracted TPM values and divided by 100 to transform the data into TP10K. For PBMCs, the log of the resulting TP10K was then compared to the pseudo-bulk data from each of the single cell methods (see Pseudo-bulk expression profile comparisons section) using Pearson correlation, with the exception of Smart-seq2. For Smart-seq2, we instead combined all FASTQ files into one joint FASTQ, and quantified TPM using RSEM using the same pipeline as for the bulk data. The percent mitochondrial transcripts for bulk data was calculated by taking the sum of TPM values over all mitochondrial genes (defined above) and dividing through by 1,000,000.

In addition to expression, *de novo* transcript construction was performed on the bulk RNA-seq data in the hope of correcting mistakes in the standard annotation. To achieve this reannotation, bulk RNA-seq data was mapped to the genome with HISAT2^52^ v 2.0.5, using the –dta flag and otherwise default parameters. *De novo* transcript construction was then performed with StringTie^59^ v0.1.18 with the parameters -v --rf -c 10 and –G, using the GTF from the scumi pipeline.

In order to quantify the improvement in performance by using the new annotation, we extracted reads from the annotated BAM file output by scumi that did not overlap any exons in the standard annotation (using the equally sampled data). We used featureCounts^53^ to count the number of these reads that overlapped an exon in the reannotated GTF output, to quantify how many more reads per cell we recovered by using the reannotated reference.

### Technical precision

We computed the extra Poisson variations that include both biological variations (assumed to be small and the same across methods in our data) and technical variations that cannot be explained by simple Poisson models^16^. High extra Poisson variations indicate either high technical variation or mispecification of the Poisson model (as seen with Smart-seq2). Specifically, let the vector 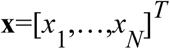 denote the expression of a gene *x* across *N* cells, where *x_i_* is the expression of that gene in cell *i*. We assume that *x_i_* follows a Poisson distribution with parameter λ*e_i_*, where λ corresponds to the number of transcripts present in this specific gene and *e_i_* accounts for the sequencing depth variation of cell *i*. For simplicity, we assume that *e_i_* is the same for all cells after median normalizing the UMI counts, i.e., 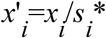 median(**s**) follows a Poisson distribution with parameter λ’. Here 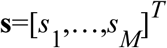, where *s*_*i*_ denotes the total number of UMIs across genes in cell *i*, and *M* is the total number of cells from a method. We then computed the extra Poisson variations defined as the empirical coefficient of variations (CV) subtracted from the Poisson CV. High extra Poisson CV indicates high technical noise. We used only genes expressed in more than 5% of the cells.

### Sampling reads

In order to correct for differences in sequencing depth between methods, we used seqtk v1.0 (https://github.com/lh3/seqtk) to sample the sequencing data, so that for each method we could analyze nearly the same average number of reads per cell. Low- and high-throughput methods were sampled independently. For a given experiment, we first decided on the average number reads per cell to sample. We usually set this equal to the lowest average number of reads per cell for a method in that experiment, except for PBMC1 Seq-Well, which had a lower average number of reads per cell, so that we chose the library with the next lowest average number of reads per cell. For each library in a given experiment, we then derived a sampling ratio by dividing this sampling target by the original average number of reads per cell in that library. We sampled the FASTQ file for each library with the ‘seqtk sample’ command, using the sampling ratio calculated above and the random seed set to 100 (after combining FASTQ files from different sequencing runs). Although we aimed to have the exact same average number of reads per cell for each library, there were some small deviations from this in practice because the number of cells we identified in each library was not always the same before and after sampling, as well as due to the random nature of the sampling. Details of the sampling for each experiment are in **Supplementary Table 5**.

### Effects of sequencing depth on gene detection

To estimate the sensitivity of each method at different sequencing depths, we sampled datasets to a given numbers of reads per cell. To efficiently sample without running scumi starting over using FASTQ files as inputs, we instead sampled the ‘molecular_info.h5’ matrix output from running scumi. The molecular_info.h5 matrix recorded the number of reads for each UMI of a gene in each selected cell. Given *N*, the total number of reads in the input FASTQ files, *C*, the number of detected cells, *M*, the total number of reads recorded in the ‘molecular_info.h5’ matrix, and *R*, the expected number of reads per cell after sampling, we needed to draw *C*\**M*\**R*/*N* reads recorded in the ‘molecular_info.h5’ matrix. This sampled molecular_info.h5 matrix was converted to the final UMI matrix after collapsing UMIs. The code for sampling is part of the scumi package.

### Automatically assigning cell types to clusters

We followed common practices for scRNA-seq data clustering. Specifically, cells were divided into non-overlapping clusters by using the Louvain community detection algorithm^56, 57^. For each cell from a dataset, we computed its *k*-nearest neighbors in that dataset, and then built a directed *k*-NN graph using all the cells from that dataset. This directed *k*-NN graph was further converted to an undirected weighed graph by using shared neighbors. The Louvain algorithm was used to partition the undirected weighted *k*-NN graph into non-overlapping clusters.

We used marker genes for each cell type to compute a score for each cell and automatically assign cell types to clusters. Both human PBMC and mouse cortex have well-annotated cell types and marker genes for each cell type^32, 60, 61^. We generated lists of marker genes for each tissue with manual curation (**Supplementary Tables 12 and 13**). The score of cell *i* for cell type *m* is a normalized version of the percentage of total counts from marker genes from cell type *m*. Assuming that there were *N_m_* marker genes for cell type *m*, we considered these *N_m_* genes combined as a ‘meta-gene’ with counts 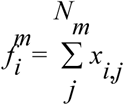 in cell *i*, where *x_i,j_* was the expression (UMI count) of marker gene *j* for cell type *m*. The meta-gene relative expression in cell *i* was its count divided by the total count *C_i_* in cell *i*. We obtained the score of cell *i* for cell type *m* as 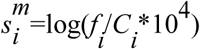.

Based on the scores, we assigned cell types to clusters using the area under the receiver operating characteristic curves (AUCs). Stated another way, for a given cluster and a given cell type *c*, a cell *i* in that cluster is a true positive if the score s_i_,_c_ is above a given threshold and a false negative otherwise. On the other hand, a cell not in that cluster is a false positive if it has a score above the threshold and a true negative otherwise. A receiver operating characteristic (ROC) curve plots the true positive rate against the false positive rate at different score thresholds. The AUC is 1.0 for perfectly assigning a cell type to a cluster (we can find a threshed score perfectly separating a cluster from the rest), and around 0.5 for randomly assigning a score to a cell. Specifically, for each cluster, the cell type with the maximum AUC was assigned to that cluster. As the same type of cells can be split into several clusters, after initial assignment of cell types to clusters, we recomputed the AUC of a cluster for a cell type by excluding other clusters of cells that were assigned to that cell type. This process was repeated until no changes in the cluster assignment. We then calculated the AUC for a cell type by merging the cluster of cells that were assigned to that cell type.

### Parameter selection for clustering analysis

We considered the following parameters for the Louvain clustering: 1) We tested different numbers of principal components (20, 30, 50, 100). 2) We varied the number of neighbors *k* used in building the *k*-NN graph. This parameter was dependent on the number of cells from each cell type and was either 15 or 30 for high-throughput datasets from PBMCs. The only exception was the Seq-Well dataset from PBMC2 which had only 551 cells and *k* was of 10, 15, or 30. For the data from low-throughput methods Smart-seq2 (from PBMC1, PBMC2, Cortex1, and Cortex2) and CEL-Seq2 (from both PBMC1 and PBMC2), the parameter was either 5, 10, or 15. 3) We varied whether we used only variable or all genes for clustering. 4) We varied the resolution parameter that could influence the number of clusters (high-throughput methods: 0.8, 1.0, 1.2, 1.5; low-throughput methods: 0.5, 0.8, 1.0, 1.2, 1.5). For the cortex datasets, the search space of parameters was the same as for the PBMC2 Seq-Well dataset with the exception of DroNc-seq data from Cortex2, which had only 892 cells so its parameter search space was the same as for Smart-seq2 data. We then scored each clustering by the formula 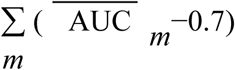, where 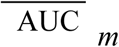 is the average AUC for cell type *m* (the average was over the clusters assigned to cell type *m*). Therefore, if cells of a single type, e.g., the B cells were partitioned into several clusters, their AUC average was smaller compared to the case where all B cells were grouped into one cluster. For this reason, the average AUC penalized partitioning cells of the same type across several clusters. The summation over all cell types in the formula favored the recovery of more distinct cell types (with AUC greater than 0.7). The actual parameters used for clustering are listed in **Supplementary Table 14**.

### Scoring each dataset to compare scRNA-seq methods

We used the AUCs to quantify the quality of each recovered cell type. After clustering analysis and assigning cell types to clusters, we also calculated the AUC of each cell type (from its cell type marker gene scores in separating that cluster of cells from the rest).

### Standard, method-specific computational pipelines

We processed each of the PBMC datasets through the standard pipeline for that scRNA-seq method and compared the output statistics to those from scumi. If there were multiple sequencing runs for a given library, we combined the FASTQ files before further analysis. Additionally, files were renamed to match the naming schemes required for a given pipeline.

For 10x Chromium v2 data, we used the Cell Ranger pipeline^9^ (v2.0.0) with the same reference genome (stored in a FASTA file) and GTF file that were used for scumi. We truncated index reads to 8 bases if the original sequence was longer. We then ran the Cell Ranger ‘count’ command with the –nosecondary flag and with –cells set to the expected number of cells. The gene by cell UMI count matrix was then extracted, with no additional cell filtering. For 10x Chromium (v3), we used a similar pipeline, except we used the Cell Ranger (v.3.0.2) pipeline with the –expect-cells flag instead of the –cells flag, and did not truncate the index reads.

For Drop-seq and Seq-Well data, we used the Drop-seq pipeline^8^ (v1.3, available at https://github.com/broadinstitute/Drop-seq). We created Drop-seq references from the GTF and FASTA files used in the scumi reference with the make_resource_bundle.py script in the Drop-seq pipeline. We processed the reads with the Drop-seq pipeline with the number of core barcodes set to the expected number of cells and using the reference we generated, though otherwise the same parameters as in Shekhar et al.^48^. To select the number of cells, we used three steps: first, we applied the cell_selection.R script in the Drop-seq pipeline to the output of the standard Drop-seq pipeline, feeding in the expected number of cells. Second, we used the DigitalExpression command from the Drop-seq pipeline to create a digital expression matrix using the cells selected. Finally, we chose a cutoff by visual inspection of the histogram of genes per.

We used the inDrops pipeline from the initial publication^7^ (available at https://github.com/indrops/indrops). We first constructed references according to the inDrops README using RSEM^54^ (v1.3) with the GTF and FASTA files used to construct our scumi references. We trimmed the reads to be the same length in all runs (read 1 50 bases, read 2 19 bases, index reads 8 bases). We then ran the pipeline using the default YAML configuration, modified to work with Bowtie^62^ (v1.1.1). We then extracted the gene by cell UMI count matrix. For cell selection, we chose the nGene cutoff used by visual inspection of the histogram of nGenes. For CEL-Seq2 data, we used the pipeline from the original publication^6^ (available at https://github.com/yanailab/CEL-Seq-pipeline). We created a Bowtie 2 reference using the bowtie2-build command in Bowtie 2^63^ (v2.2.1), using the same FASTA file as for the other pipelines. We processed the reads with the CEL-Seq2 pipeline with the default configuration file, modified to include our updated reference. We extracted and combined the gene by cell UMI count matrix. For cell selection, we chose the nGene cutoff used by visual inspection of the histogram of nGenes.

For Smart-seq2, we used the same pipeline as in the scumi results – HISAT2^52^ to map the FASTQ file, featureCounts^53^ to annotate the BAM file, followed by counting reads that overlap the exons of a unique gene – except for cell selection we chose the nGene cutoff by visual inspection of the histogram of nGenes.

### Clustering combined datasets

For the PBMC and cortex datasets, to find cell types that were missed in analyzing each dataset independently, we took the count data for each method (after sampling the same number of reads and cells as describe above) in each experiment, and merged them into one count matrix. For Smart-seq2 we used reads counts, while for the other methods we used UMI counts. We loaded the results into R using Seurat, normalized to log counts per 10,000, and scaled the data. We identified variable genes separately for each library using the ‘FindVariableGenes’ function (with x.low.cutoff=.1 and x.high.cutoff=5), and keeping genes that were variable in at least 5 libraries for the PBMCs or at least 3 libraries for the Cortex. We then performed PCA with those genes, and used the results for Harmony^33^ with parameters, theta = 4 (for PBMCs) or theta=6 (for cortex), nclust = 50, and max.iter.cluster = 100. The resulting 20 Harmony components were used for clustering and t-SNE visualization with default parameters. Cell type annotations were based on the PBMC and cortex marker genes. For cortex, we also manually identified two clusters, one that seemed to correspond to the unidentified cells in sci-RNA-seq, and the other that seemed to be fibroblast-like cells (not considered in our analysis), both of which we labelled as unassigned.

To check the Harmony cell type results, the data for each library were extracted from the Harmony object one by one, and the cell type clusters identified in the combined Harmony analysis were reannotated using the AUC cell type identification method with only cells from that one library (excluding unassigned cells). This updated annotation was compared to the combined Harmony annotation to determine whether the same cell types were identified.

### Data and software availability

RNA-Seq data generated in this project are available from Gene Expression Omnibus with accession number TBD and the Single Cell Portal (https://portals.broadinstitute.org/single_cell). The scumi Python package is available freely from bitbucket repository https://bitbucket.org/jerry00/scumi-dev/src/master/. The R scripts (used to assign cell types to clusters based on a set of marker genes, for parameter selecting for clustering analysis, and for filtering low-quality cells) are available from bitbucket repository https://bitbucket.org/jerry00/scumi-dev/src/master/.

## Supplementary Figures

**Supplementary Figure 1.** Description of scRNA-seq methods evaluated.

Salient details for seven protocols tested in this paper.

**Supplementary Figure 2.** Flowchart detailing computational analysis.

(**a**) scumi workflow, (**b**) removing low quality cell barcodes, (**c**) profiling samples, (**d**) bulk data workflow.

**Supplementary Figure 3.** Characterization of genome alignments for sequence reads.

(**a**) Mixture, (**b**) PBMCs, (**c**) Cortex. For each pair of bar graphs, experiment 1 is on the left and experiment 2 is on the right. For Smart-seq2, there were no poly(T) reads due to the full transcriptome coverage and the library construction using transposase-based Nextera reagents to attach adapters to both ends of cDNA fragments. Reads were assigned in the following order: no poly(T), unmapped or multi-mapped, ambiguous (mapping to a single location that overlaps 2 or more genes), and then one of the remaining categories. Reads were assigned as antisense only for the cortex datasets (**c**). % of reads may not sum to 100 due to rounding and numbers not shown for fraction of reads in categories with <2%.

**Supplementary Figure 4.** Mitochondrial ratios.

(**a**) human and mouse cell line mixtures, (**b**) PBMCs, and (**c**) cortex. The mitochondrial ratio distributions in cells from each method. Each box represents the median (labeled on the right of each box) and the first and the third quantiles per method.

**Supplementary Figure 5.** Sensitivity relative to number of cells.

(**a**) human cells and mouse cells from Mixture experiments. Multiplet cells are not shown in this plot. (**b**) PBMC. (**c**) cortex. The means of the number of detected genes at given number of cells for each method. For each curve, the cells were ordered based on the total number of genes from most to least. The right most point at the end of each curve shows the average number of detected genes for the final selected number of cells.

**Supplementary Figure 6.** Rarefaction curves for PBMC datasets.

(**a**) PBMC1, (**b**) PBMC2. The effect of sequencing depth on the median number of detected genes per cell. The far right point of each curve represents the median number of detected genes per cell at full sequencing depth.

**Supplementary Figure 7.** Fraction of reads from each species in Mixture experiments.

(**a**) Mixture1, (**b**) Mixture2 – showing % UMIs (or reads for Smart-seq2) aligned to either mouse or human. Fraction of genes detected from the “wrong” species increases with sequencing depth. Each dot represents a cell. Each line is the robust linear regression fitted line, and the number along each line is the slope of the regression line.

**Supplementary Figure 8.** Correlation of pseudo-bulk expression profiles.

The Pearson correlation coefficients among pseudo-bulk profiles of (**a**) human and (**b**) mouse cells obtained from different methods. High correlations indicate high reproducibility. The number in each cell represents the Pearson correlation coefficient between two pseudo-bulks. The Pearson correlation coefficient matrix were clustered using complete-linkage hierarchical clustering. The expression vectors of cells from a method were added together to form a pseudo-bulk profile. Human cells and mouse cells were analyzed independently.

**Supplementary Figure 9.** Technical precision plots for mixture experiments.

Distributions of the extra Poisson coefficients of variations (“Extra Poisson CV”, *y* axis) from each method (*x* axis). Each box represents the median (labeled above each box) and the first and the third quantiles per method. (**a**,**b**) Human cells, (**c**,**d**) mouse cells – from Mixture1 (left) and Mixture2 (right).

**Supplementary Figure 10.** Correlation of bulk and scRNA-seq PBMC datasets

(**a**) PBMC1, (**b**) PBMC2. For each library, we compared the log transcripts per 10K (log TP10K) for bulk and pseudo-bulk from scRNA-seq datasets on a gene by gene basis, plotted as a 2 dimensional histogram. The color of each box in the plane is calculated from the log count of genes in that square. In all cases, the majority of the weight is along the diagonal, suggesting high concordance between bulk and single cell datasets. The Pearson correlation is indicated for each library.

**Supplementary Figure 11.** t-SNE plots for PBMCs.

(**a**) PBMC1, (**b**) PBMC2

**Supplementary Figure 12.** t-SNE plots for cortex nuclei, experiment 2.

**Supplementary Figure 13.** Analysis of combined PBMC datasets.

(**a**) t-SNE plot generated with Harmony clustering all PBMC cells in this study. (**b**) All libraries contain cells of every cell type, according to this joint annotation. This differs from the individual level clustering results, in which many libraries are missing particular cell types. (**c**) PBMC1 and (**d**) PBMC2. For each annotated cell type and method in the jointly clustered dataset (*y* axis), we calculated the percentage of cells from that cell type that come from each cell type in the individual level clustering results (*x* axis). This is denoted by the color of the corresponding boxes.

**Supplementary Figure 14.** Analysis of combined cortex datasets.

(**a**) t-SNE plot generated with Harmony clustering all cortex nuclei in this study. (**b**) All libraries contain nuclei of every cell type, according to this joint annotation. (**c**) For each annotated cell type and method in the jointly clustered dataset (*y* axis), we calculated the percentage of nuclei from that cell type that come from each cell type in the individual level clustering results (*x* axis).

**Supplementary Figure 15.** Benchmarking scumi against method-specific pipelines.

(**a**) and (**b**) UMIs per cell, except reads per cell for Smart-seq2. (**c**) and (**d**) genes per cell. *r* is the Pearson correlation coefficient. A line indicates the case where the detected number of genes (or UMIs) from both methods are the same. Color of points encodes the density (two-dimensional kernel density) estimate at each point.

## Supplementary Tables

**Supplementary Table 1.** Mixture experiment sequencing metrics

**Supplementary Table 2.** PBMC experiment sequencing metrics

**Supplementary Table 3.** Cortex experiment sequencing metrics

**Supplementary Table 4.** Additional reads mapped using transcriptome derived from bulk RNA-seq

**Supplementary Table 5.** Read sampling.

**Supplementary Table 6.** PBMC cell types from Harmony not confirmed in separate clustering

**Supplementary Table 7.** Cost and time for each method

**Supplementary Table 8.** Cell preparation for mixture experiment

**Supplementary Table 9.** Cell passage information for mixture experiment

**Supplementary Table 10.** Sequencing runs.

**Supplementary Table 11.** Read processing for each library type

**Supplementary Table 12.** Gene classifiers for PBMCs

**Supplementary Table 13.** Gene classifiers for cortex

**Supplementary Table 14.** Optimized clustering parameters for PBMC and cortex datasets

