## Supplementary Figures for "Systematic comparative analysis of single cell RNA-sequencing methods"

Supp. Fig. 1

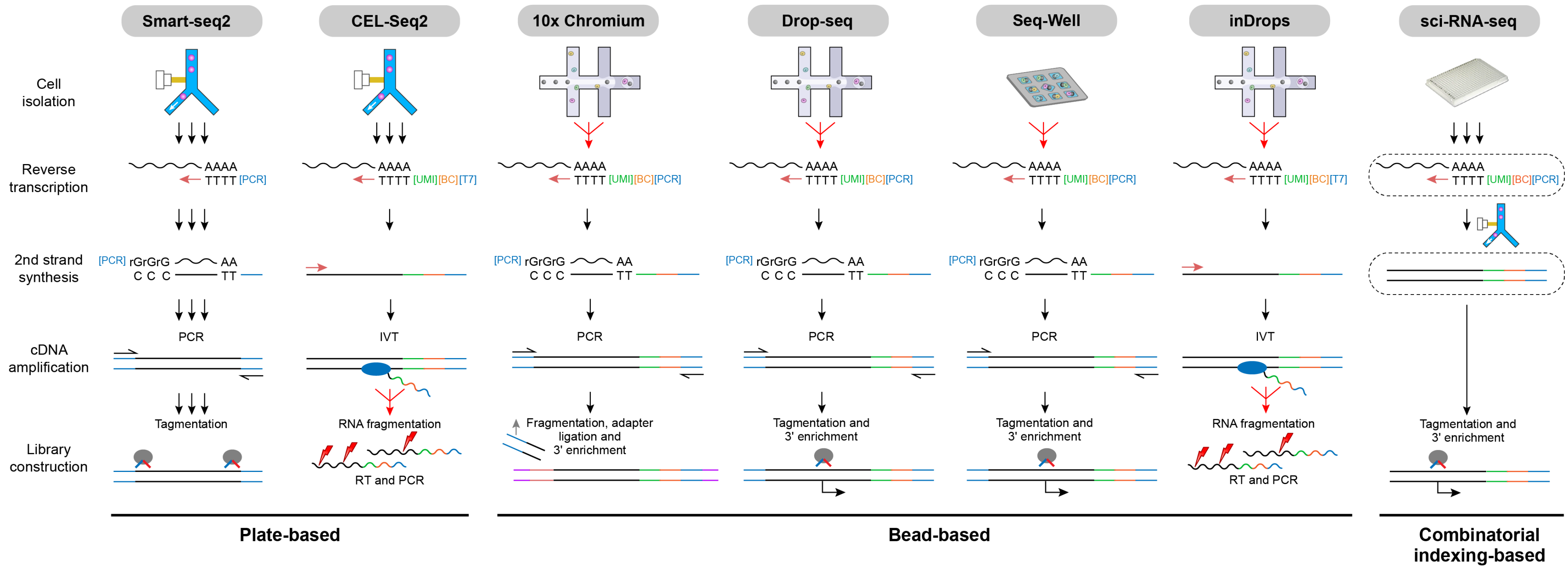
Smart-seq2
CEL-Seq2
10x Chromium
Drop-seq
Seq-Well
inDrops
sci-RNA-seq
Cell isolation
Reverse transcription
2nd strand synthesis
cDNA amplification
Library construction
AAAA
TTTT [PCR]
TTTT [UMI][BC][T7]
TTTT [UMI][BC][PCR]
[PCR] rGrGrG AA
C C C TT
PCR
IVT
Tagmentation
RNA fragmentation
Fragmentation, adapter ligation and 3' enrichment
Tagmentation and 3' enrichment
RT and PCR
Plate-based
Bead-based
Combinatorial indexing-based

Supp. Fig. 2

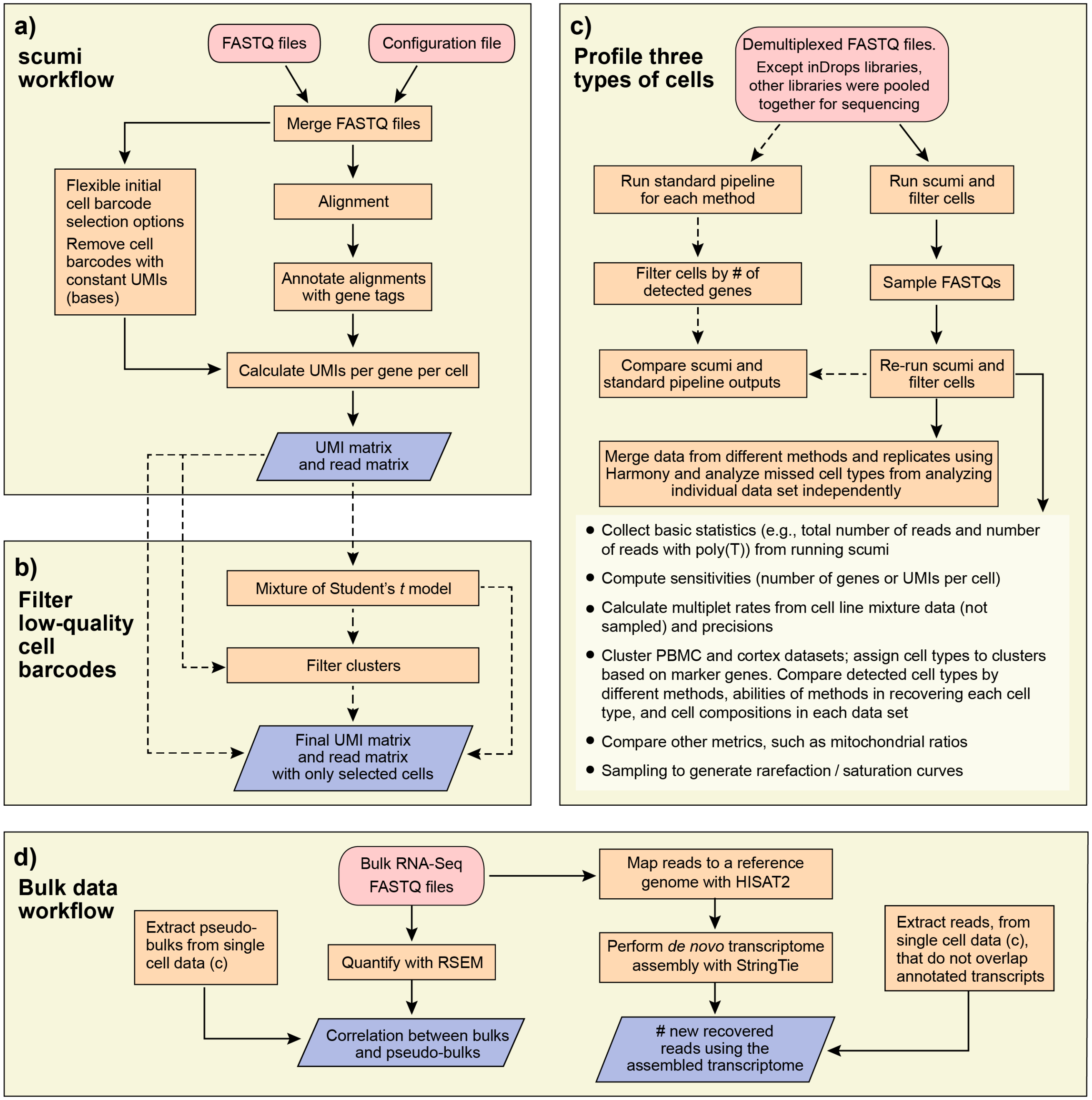

a)
scumi workflow
FASTQ files
Configuration file
Merge FASTQ files
Flexible initial cell barcode selection options
Remove cell barcodes with constant UMIs (bases)
Alignment
Annotate alignments with gene tags
Calculate UMIs per gene per cell
UMI matrix and read matrix
b)
Filter low-quality cell barcodes
Mixture of Student's t model
Filter clusters
Final UMI matrix and read matrix with only selected cells
c)
Profile three types of cells
Demultiplexed FASTQ files. Except inDrops libraries, other libraries were pooled together for sequencing
Run standard pipeline for each method
Run scumi and filter cells
Filter cells by # of detected genes
Sample FASTQs
Compare scumi and standard pipeline outputs
Re-run scumi and filter cells
Merge data from different methods and replicates using Harmony and analyze missed cell types from analyzing individual data set independently
• Collect basic statistics (e.g., total number of reads and number of reads with poly(T)) from running scumi
• Compute sensitivities (number of genes or UMIs per cell)
• Calculate multiplet rates from cell line mixture data (not sampled) and precisions
• Cluster PBMC and cortex datasets; assign cell types to clusters based on marker genes. Compare detected cell types by different methods, abilities of methods in recovering each cell type, and cell compositions in each data set
• Compare other metrics, such as mitochondrial ratios
• Sampling to generate rarefaction / saturation curves
d)
Bulk data workflow
Bulk RNA-Seq FASTQ files
Map reads to a reference genome with HISAT2
Extract pseudo-bulks from single cell data (c)
Quantify with RSEM
Perform de novo transcriptome assembly with StringTie
Extract reads, from single cell data (c), that do not overlap annotated transcripts
Correlation between bulks and pseudo-bulks
### new recovered reads using the assembled transcriptome

Supp. Fig. 3

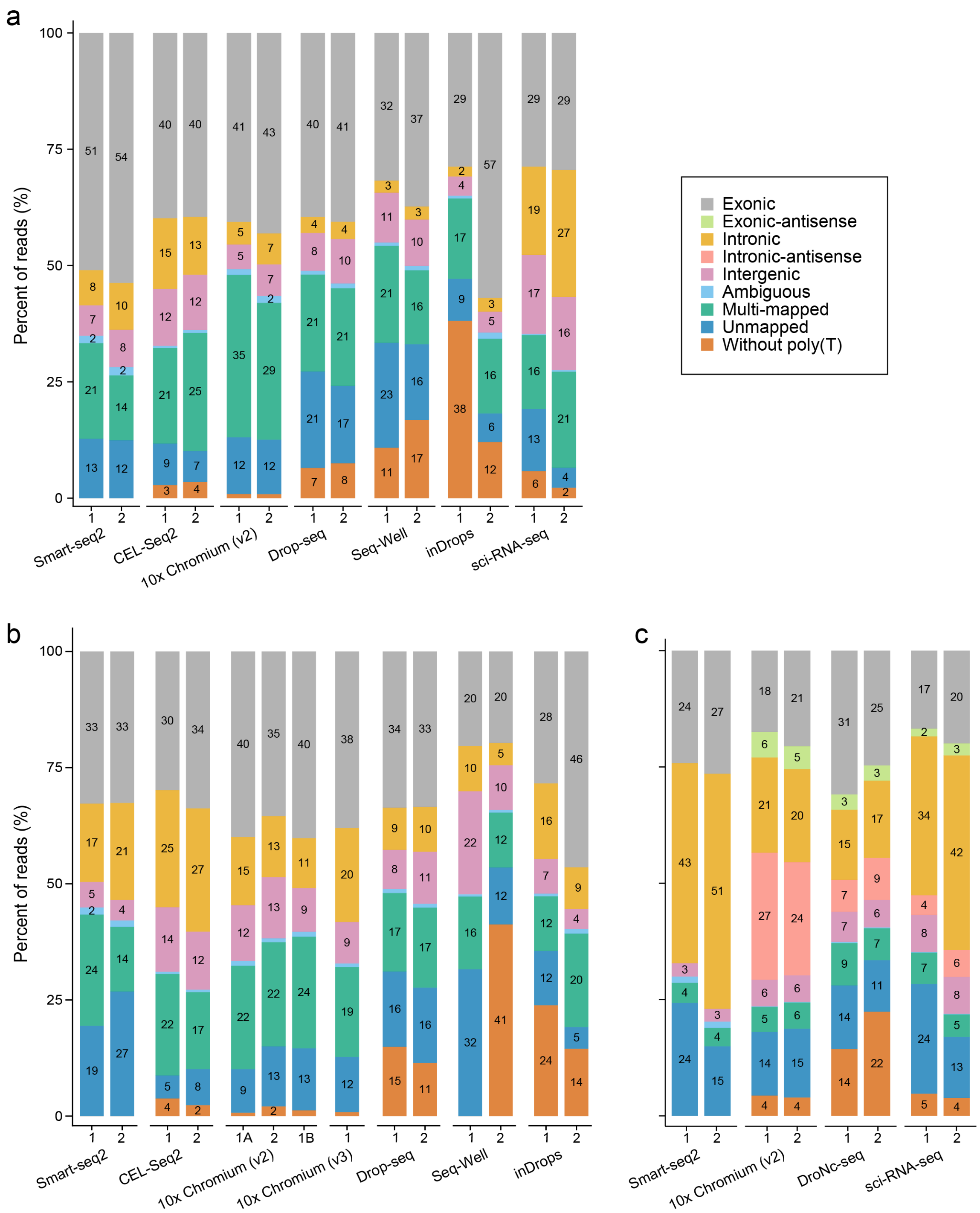
a
b
c
Percent of reads (%)
0
25
50
75
100
Exonic
Exonic-antisense
Intronic
Intronic-antisense
Intergenic
Ambiguous
Multi-mapped
Unmapped
Without poly(T)
Smart-seq2
CEL-Seq2
10x Chromium (v2)
10x Chromium (v3)
Drop-seq
Seq-Well
inDrops
sci-RNA-seq
DroNc-seq
1A
1B

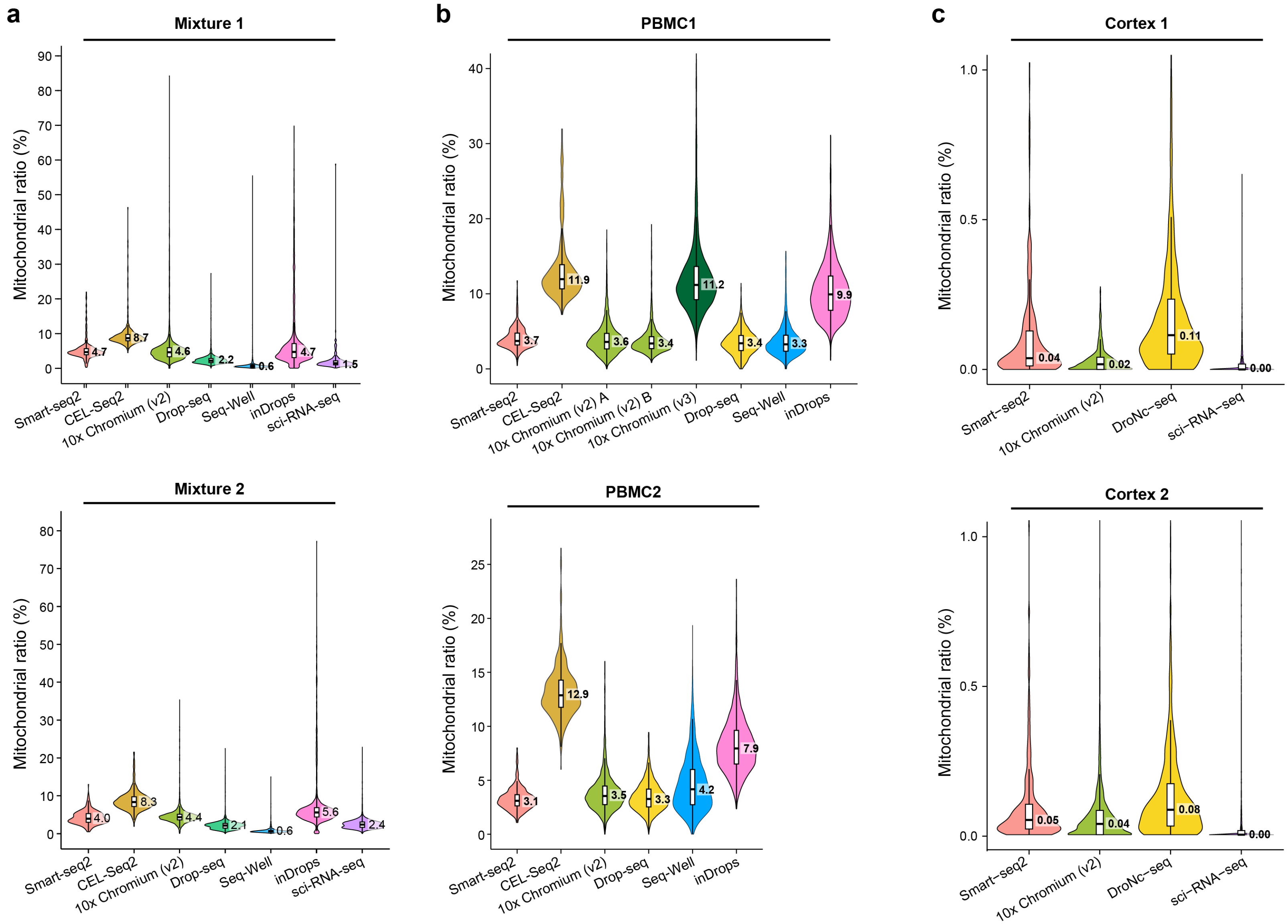
a
Mixture 1
Mitochondrial ratio (%)
Smart-seq2
CEL-Seq2
10x Chromium (v2)
Drop-seq
Seq-Well
inDrops
sci-RNA-seq
4.7
8.7
4.6
2.2
0.6
4.7
1.5
Mixture 2
4.0
8.3
4.4
2.1
0.6
5.6
2.4
b
PBMC1
10x Chromium (v2) A
10x Chromium (v2) B
10x Chromium (v3)
3.7
11.9
3.6
3.4
11.2
3.4
3.3
9.9
PBMC2
3.1
12.9
3.5
3.3
4.2
7.9
c
Cortex 1
DroNc−seq
sci−RNA−seq
0.04
0.02
0.11
0.00
Cortex 2
0.05
0.04
0.08
0.00

Supp. Fig. 5

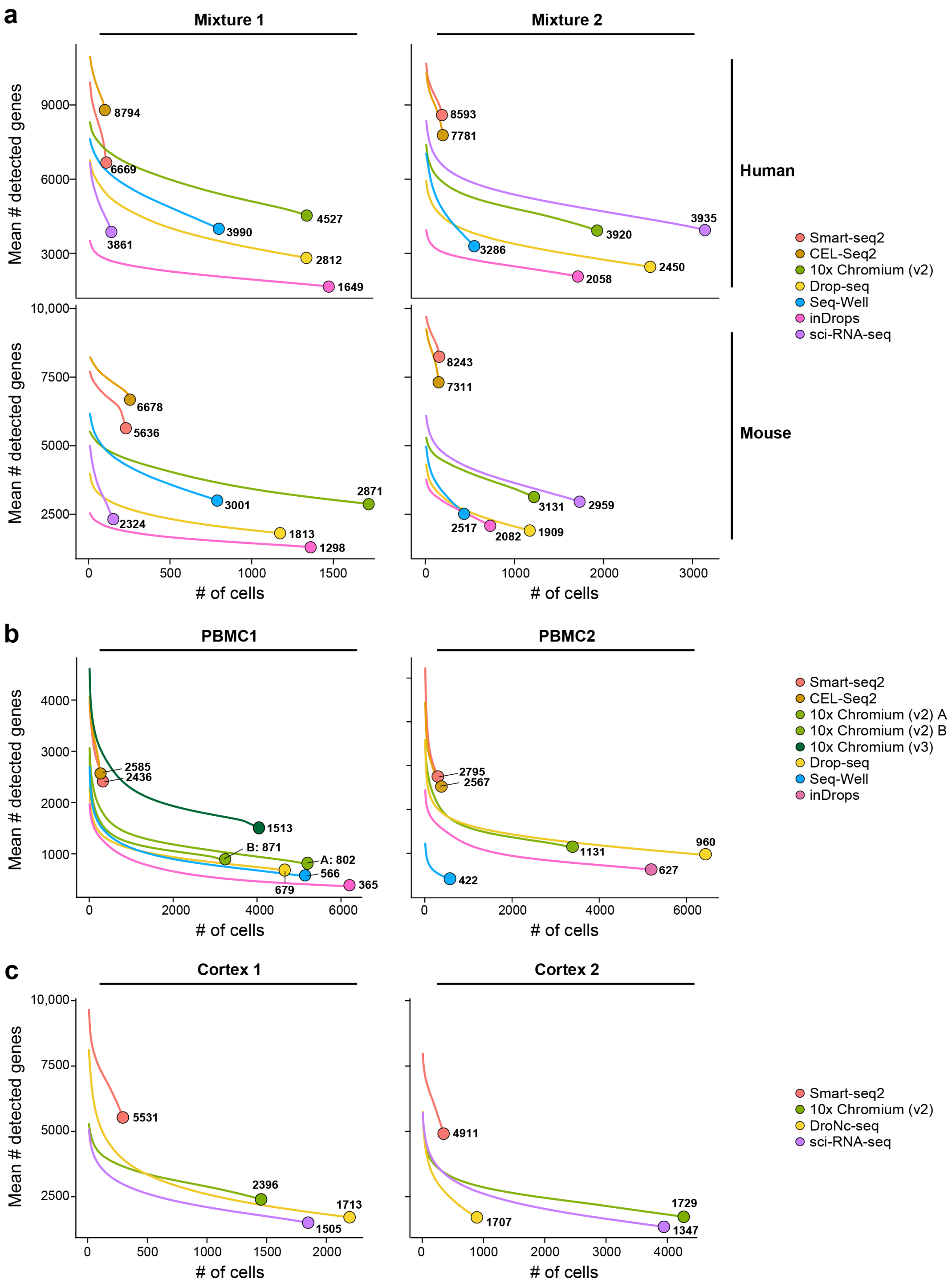

a
Mixture 1
Mixture 2
Human
Mouse
Mean # detected genes
9000
6000
3000
10,000
7500
5000
2500
8794
6669
4527
3990
3861
2812
1649
8593
7781
3935
3920
3286
2450
2058
6678
5636
2871
3001
2324
1813
1298
8243
7311
3131
2959
2517
2082
1909
0
500
1000
1500
2000
3000
### of cells
Smart-seq2
CEL-Seq2
10x Chromium (v2)
Drop-seq
Seq-Well
inDrops
sci-RNA-seq
b
PBMC1
PBMC2
4000
3000
2000
1000
2585
2436
1513
B: 871
A: 802
566
679
365
2795
2567
1131
960
627
422
6000
10x Chromium (v2) A
10x Chromium (v2) B
10x Chromium (v3)
c
Cortex 1
Cortex 2
5531
2396
1713
1505
4911
1729
1707
1347
DroNc-seq

Supp. Fig. 6

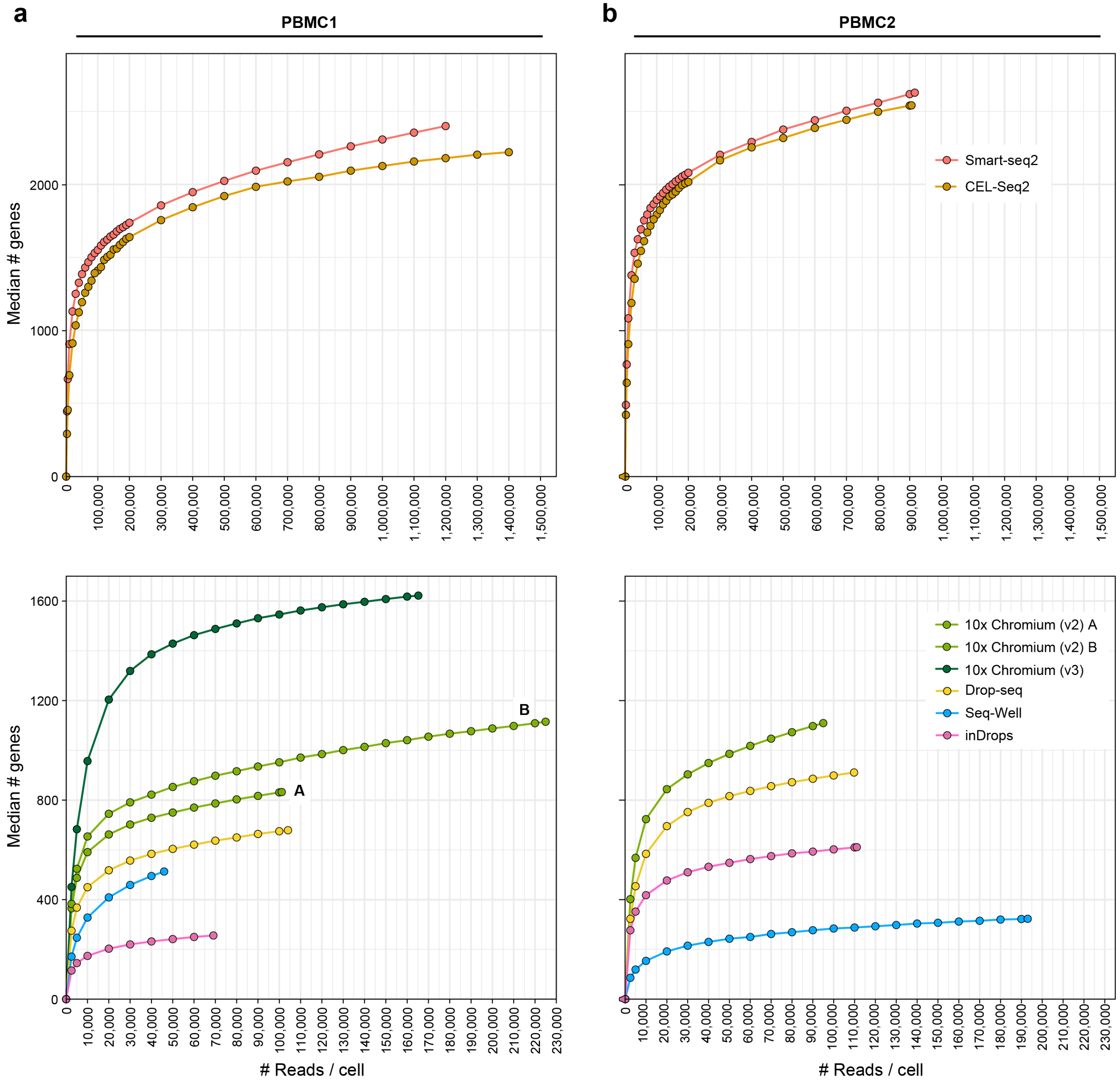

a
b
PBMC1
PBMC2
Median # genes
### Reads / cell
Smart-seq2
CEL-Seq2
10x Chromium (v2) A
10x Chromium (v2) B
10x Chromium (v3)
Drop-seq
Seq-Well
inDrops
A
B

Supp. Fig. 7

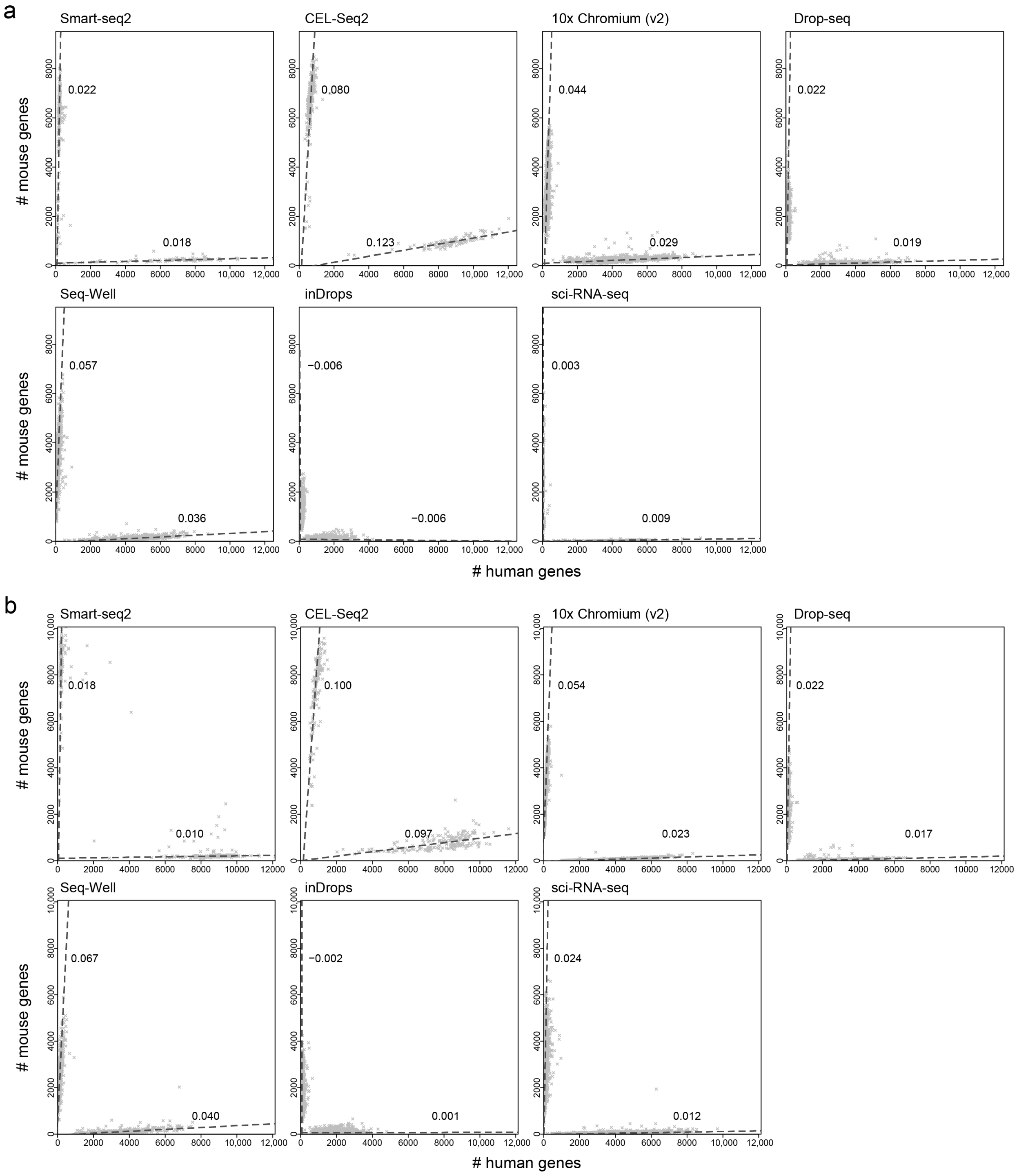
a
Smart-seq2
0.022
0.018
CEL-Seq2
0.080
0.123
10x Chromium (v2)
0.044
0.029
Drop-seq
0.022
0.019
Seq-Well
0.057
0.036
inDrops
−0.006
−0.006
sci-RNA-seq
0.003
0.009
### mouse genes
### human genes
b
Smart-seq2
0.018
0.010
CEL-Seq2
0.100
0.097
10x Chromium (v2)
0.054
0.023
Drop-seq
0.022
0.017
Seq-Well
0.067
0.040
inDrops
−0.002
0.001
sci-RNA-seq
0.024
0.012
### mouse genes
### human genes

Supp. Fig. 8

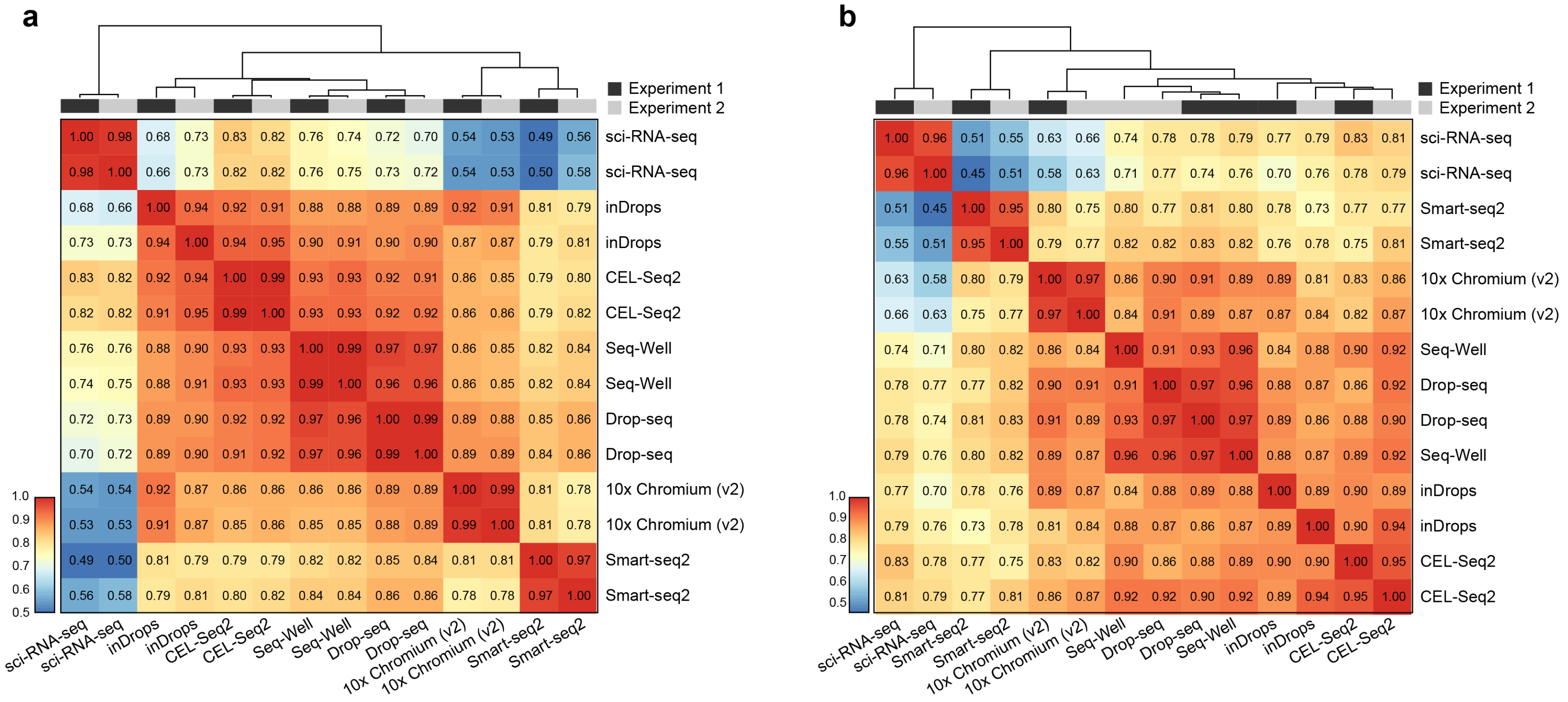
a
Experiment 1
Experiment 2
1.00 0.98 0.68 0.73 0.83 0.82 0.76 0.74 0.72 0.70 0.54 0.53 0.49 0.56 sci-RNA-seq
0.98 1.00 0.66 0.73 0.82 0.82 0.76 0.75 0.73 0.72 0.54 0.53 0.50 0.58 sci-RNA-seq
0.68 0.66 1.00 0.94 0.92 0.91 0.88 0.88 0.89 0.89 0.92 0.91 0.81 0.79 inDrops
0.73 0.73 0.94 1.00 0.94 0.95 0.90 0.91 0.90 0.90 0.87 0.87 0.79 0.81 inDrops
0.83 0.82 0.92 0.94 1.00 0.99 0.93 0.93 0.92 0.91 0.86 0.85 0.79 0.80 CEL-Seq2
0.82 0.82 0.91 0.95 0.99 1.00 0.93 0.93 0.92 0.92 0.86 0.86 0.79 0.82 CEL-Seq2
0.76 0.76 0.88 0.90 0.93 0.93 1.00 0.99 0.97 0.97 0.86 0.85 0.82 0.84 Seq-Well
0.74 0.75 0.88 0.91 0.93 0.93 0.99 1.00 0.96 0.96 0.86 0.85 0.82 0.84 Seq-Well
0.72 0.73 0.89 0.90 0.92 0.92 0.97 0.96 1.00 0.99 0.89 0.88 0.85 0.86 Drop-seq
0.70 0.72 0.89 0.90 0.91 0.92 0.97 0.96 0.99 1.00 0.89 0.89 0.84 0.86 Drop-seq
0.54 0.54 0.92 0.87 0.86 0.86 0.86 0.86 0.89 0.89 1.00 0.99 0.81 0.78 10x Chromium (v2)
0.53 0.53 0.91 0.87 0.85 0.86 0.85 0.85 0.88 0.89 0.99 1.00 0.81 0.78 10x Chromium (v2)
0.49 0.50 0.81 0.79 0.79 0.79 0.82 0.82 0.85 0.84 0.81 0.81 1.00 0.97 Smart-seq2
0.56 0.58 0.79 0.81 0.80 0.82 0.84 0.84 0.86 0.86 0.78 0.78 0.97 1.00 Smart-seq2
1.0 0.9 0.8 0.7 0.6 0.5
sci-RNA-seq sci-RNA-seq inDrops inDrops CEL-Seq2 CEL-Seq2 Seq-Well Seq-Well Drop-seq Drop-seq 10x Chromium (v2) 10x Chromium (v2) Smart-seq2 Smart-seq2
b
Experiment 1
Experiment 2
1.00 0.96 0.51 0.55 0.63 0.66 0.74 0.78 0.78 0.79 0.77 0.79 0.83 0.81 sci-RNA-seq
0.96 1.00 0.45 0.51 0.58 0.63 0.71 0.77 0.74 0.76 0.70 0.76 0.78 0.79 sci-RNA-seq
0.51 0.45 1.00 0.95 0.80 0.75 0.80 0.77 0.81 0.80 0.78 0.73 0.77 0.77 Smart-seq2
0.55 0.51 0.95 1.00 0.79 0.77 0.82 0.82 0.83 0.82 0.76 0.78 0.75 0.81 Smart-seq2
0.63 0.58 0.80 0.79 1.00 0.97 0.86 0.90 0.91 0.89 0.89 0.81 0.83 0.86 10x Chromium (v2)
0.66 0.63 0.75 0.77 0.97 1.00 0.84 0.91 0.89 0.87 0.87 0.84 0.82 0.87 10x Chromium (v2)
0.74 0.71 0.80 0.82 0.86 0.84 1.00 0.91 0.93 0.96 0.84 0.88 0.90 0.92 Seq-Well
0.78 0.77 0.77 0.82 0.90 0.91 0.91 1.00 0.97 0.96 0.88 0.87 0.86 0.92 Drop-seq
0.78 0.74 0.81 0.83 0.91 0.89 0.93 0.97 1.00 0.97 0.89 0.86 0.88 0.90 Drop-seq
0.79 0.76 0.80 0.82 0.89 0.87 0.96 0.96 0.97 1.00 0.88 0.87 0.89 0.92 Seq-Well
0.77 0.70 0.78 0.76 0.89 0.87 0.84 0.88 0.89 0.88 1.00 0.89 0.90 0.89 inDrops
0.79 0.76 0.73 0.78 0.81 0.84 0.88 0.87 0.86 0.87 0.89 1.00 0.90 0.94 inDrops
0.83 0.78 0.77 0.75 0.83 0.82 0.90 0.86 0.88 0.89 0.90 0.90 1.00 0.95 CEL-Seq2
0.81 0.79 0.77 0.81 0.86 0.87 0.92 0.92 0.90 0.92 0.89 0.94 0.95 1.00 CEL-Seq2
1.0 0.9 0.8 0.7 0.6 0.5
sci-RNA-seq sci-RNA-seq Smart-seq2 Smart-seq2 10x Chromium (v2) 10x Chromium (v2) Seq-Well Drop-seq Drop-seq Seq-Well inDrops inDrops CEL-Seq2 CEL-Seq2

Supp. Fig. 9

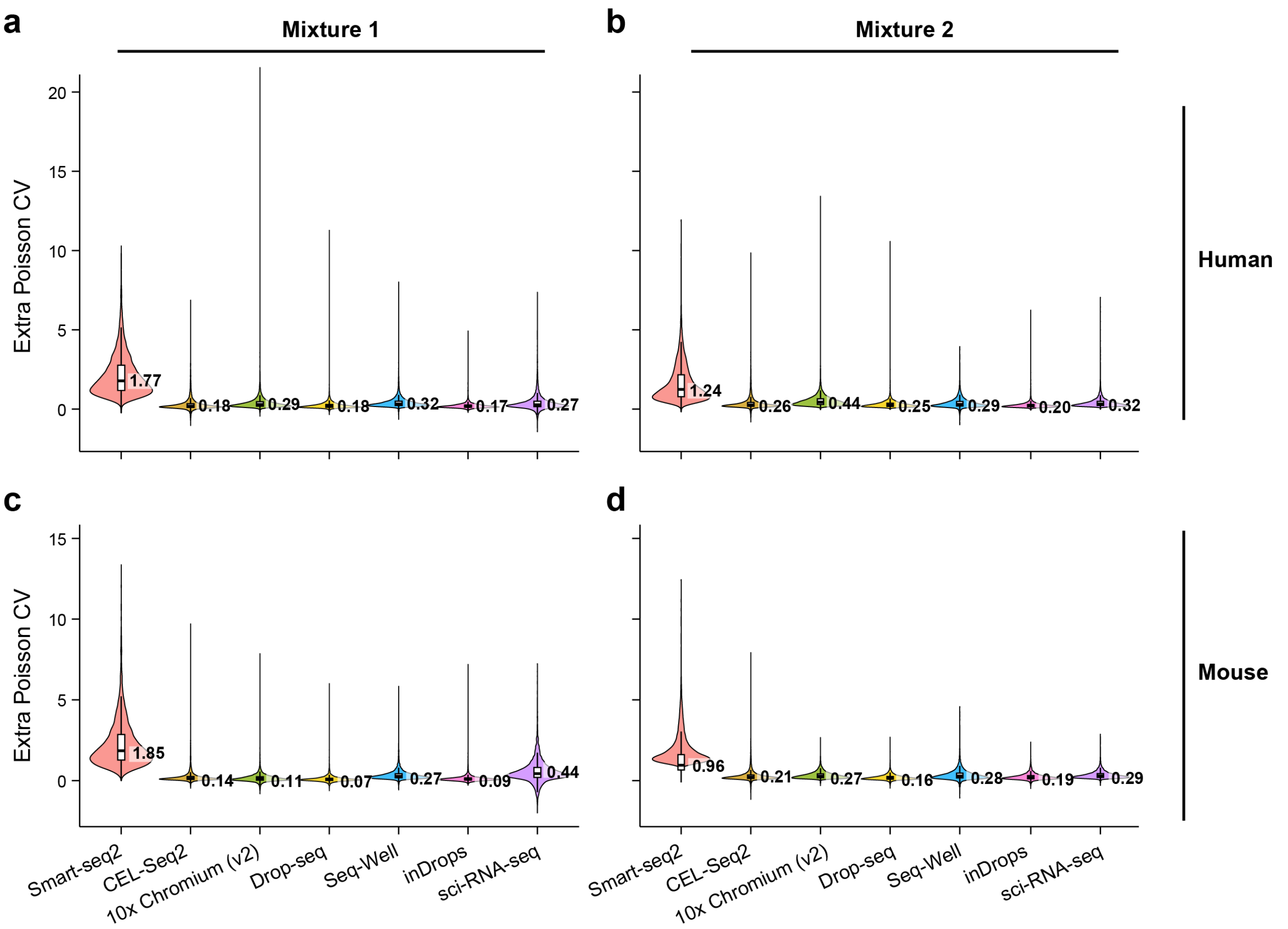

a
Mixture 1
b
Mixture 2
Human
Mouse
c
d
Extra Poisson CV
20
15
10
5
0
1.77
0.18
0.29
0.18
0.32
0.17
0.27
1.24
0.26
0.44
0.25
0.29
0.20
0.32
1.85
0.14
0.11
0.07
0.27
0.09
0.44
0.96
0.21
0.27
0.16
0.28
0.19
0.29
Smart-seq2
CEL-Seq2
10x Chromium (v2)
Drop-seq
Seq-Well
inDrops
sci-RNA-seq

### Supp. Fig. 10

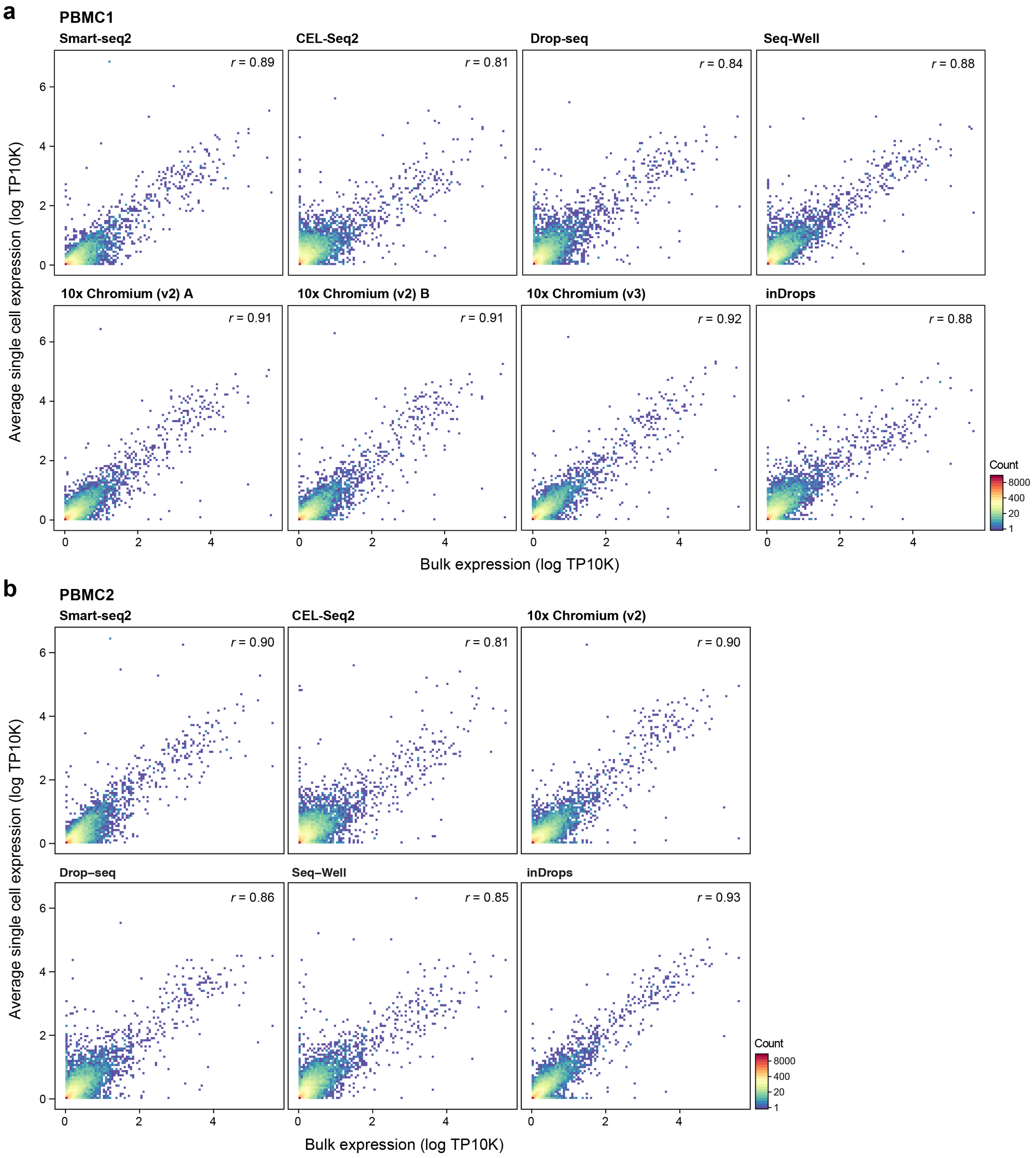
a
PBMC1
Smart-seq2
r = 0.89
CEL-Seq2
r = 0.81
Drop-seq
r = 0.84
Seq-Well
r = 0.88
10x Chromium (v2) A
r = 0.91
10x Chromium (v2) B
r = 0.91
10x Chromium (v3)
r = 0.92
inDrops
r = 0.88
Average single cell expression (log TP10K)
Bulk expression (log TP10K)
Count
8000
400
20
1
b
PBMC2
Smart-seq2
r = 0.90
CEL-Seq2
r = 0.81
10x Chromium (v2)
r = 0.90
Drop–seq
r = 0.86
Seq–Well
r = 0.85
inDrops
r = 0.93
Average single cell expression (log TP10K)
Bulk expression (log TP10K)
Count
8000
400
20
1

Supp. Fig. 11

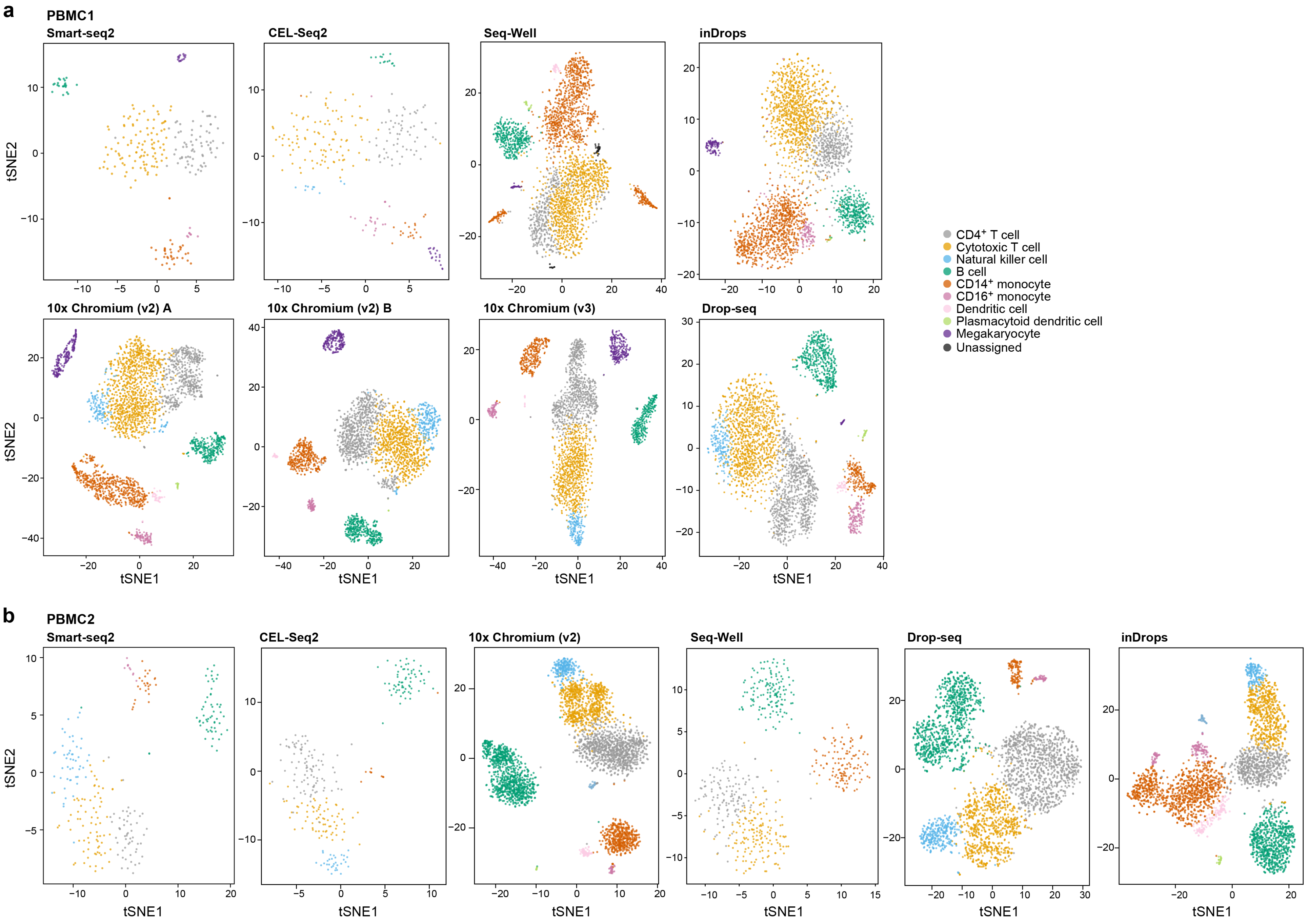
a
PBMC1
Smart-seq2
CEL-Seq2
Seq-Well
inDrops
10x Chromium (v2) A
10x Chromium (v2) B
10x Chromium (v3)
Drop-seq
tSNE1
tSNE2
CD4+ T cell
Cytotoxic T cell
Natural killer cell
B cell
CD14+ monocyte
CD16+ monocyte
Dendritic cell
Plasmacytoid dendritic cell
Megakaryocyte
Unassigned
b
PBMC2
Smart-seq2
CEL-Seq2
10x Chromium (v2)
Seq-Well
Drop-seq
inDrops
tSNE1
tSNE2

Supp. Fig. 12

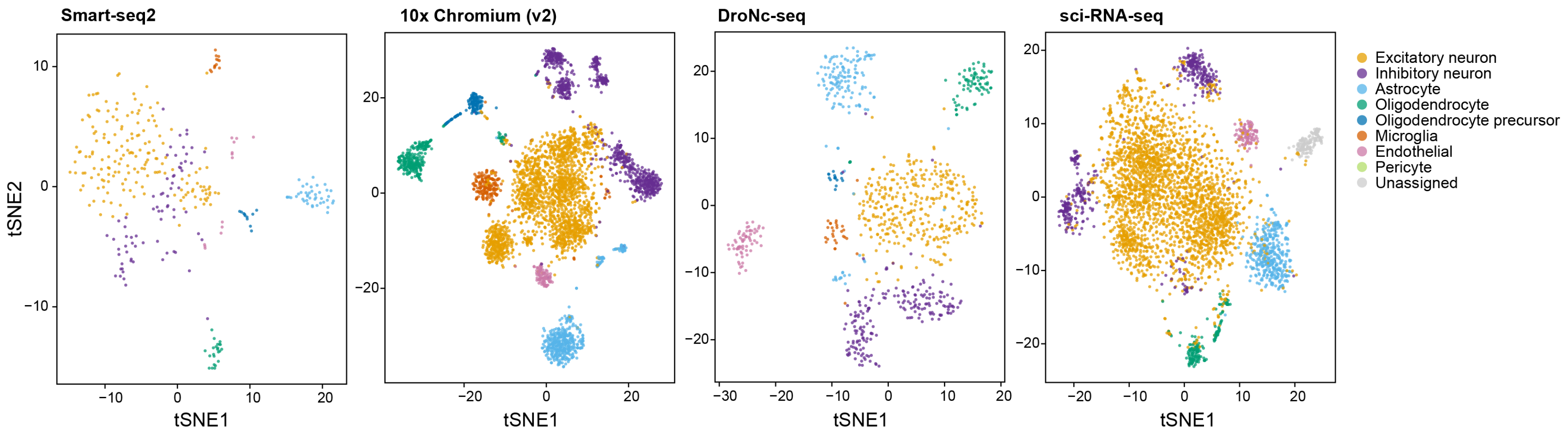
Smart-seq2
10x Chromium (v2)
DroNc-seq
sci-RNA-seq
tSNE2
tSNE1
tSNE1
tSNE1
tSNE1
Excitatory neuron
Inhibitory neuron
Astrocyte
Oligodendrocyte
Oligodendrocyte precursor
Microglia
Endothelial
Pericyte
Unassigned

Supp. Fig. 13

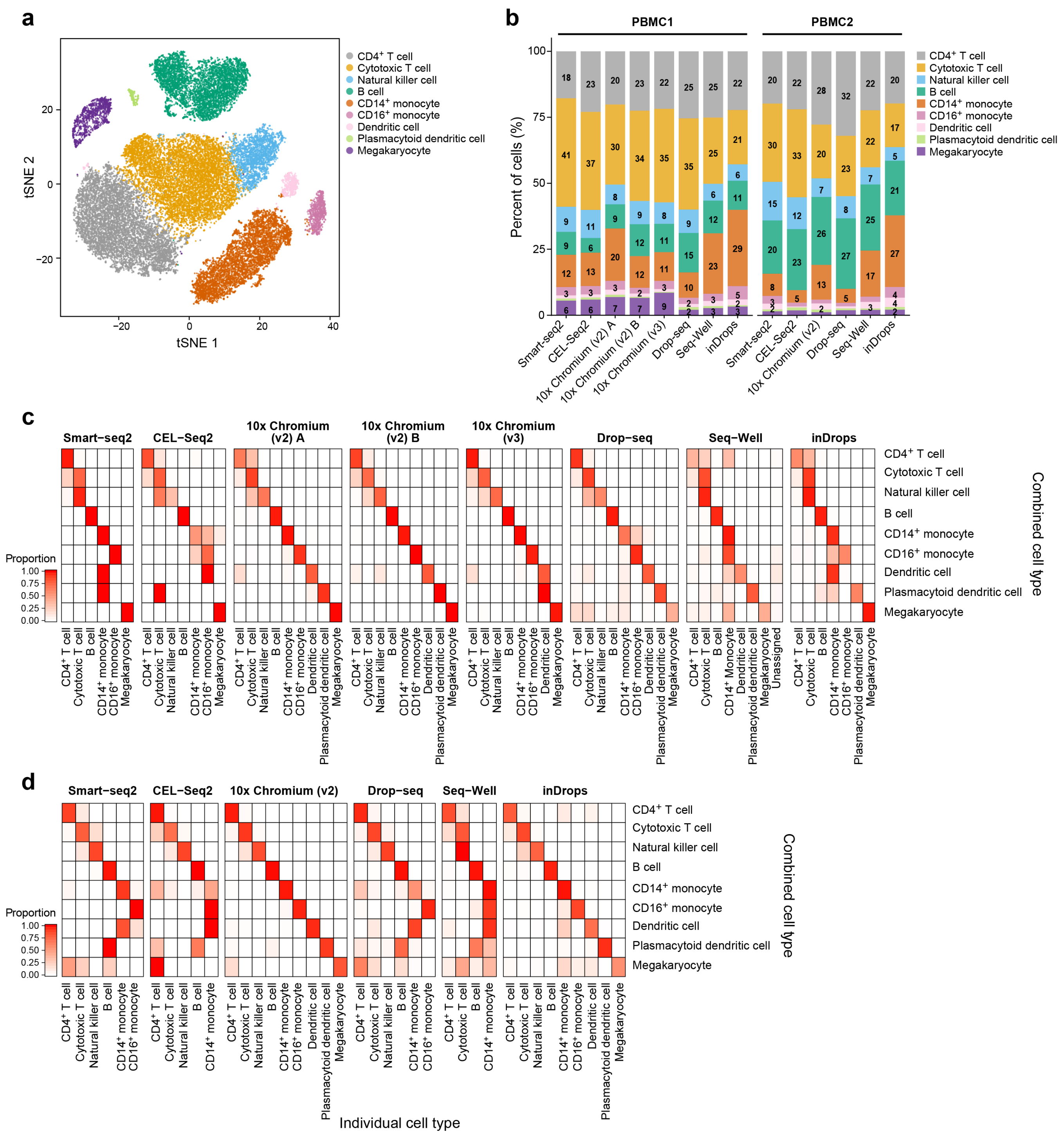
a
CD4+ T cell
Cytotoxic T cell
Natural killer cell
B cell
CD14+ monocyte
CD16+ monocyte
Dendritic cell
Plasmacytoid dendritic cell
Megakaryocyte
tSNE 1
tSNE 2
b
PBMC1
PBMC2
Percent of cells (%)
Smart-seq2
CEL-Seq2
10x Chromium (v2) A
10x Chromium (v2) B
10x Chromium (v3)
Drop-seq
Seq-Well
inDrops
10x Chromium (v2)
c
Proportion
Combined cell type
Unassigned
d
Individual cell type

Supp. Fig. 14

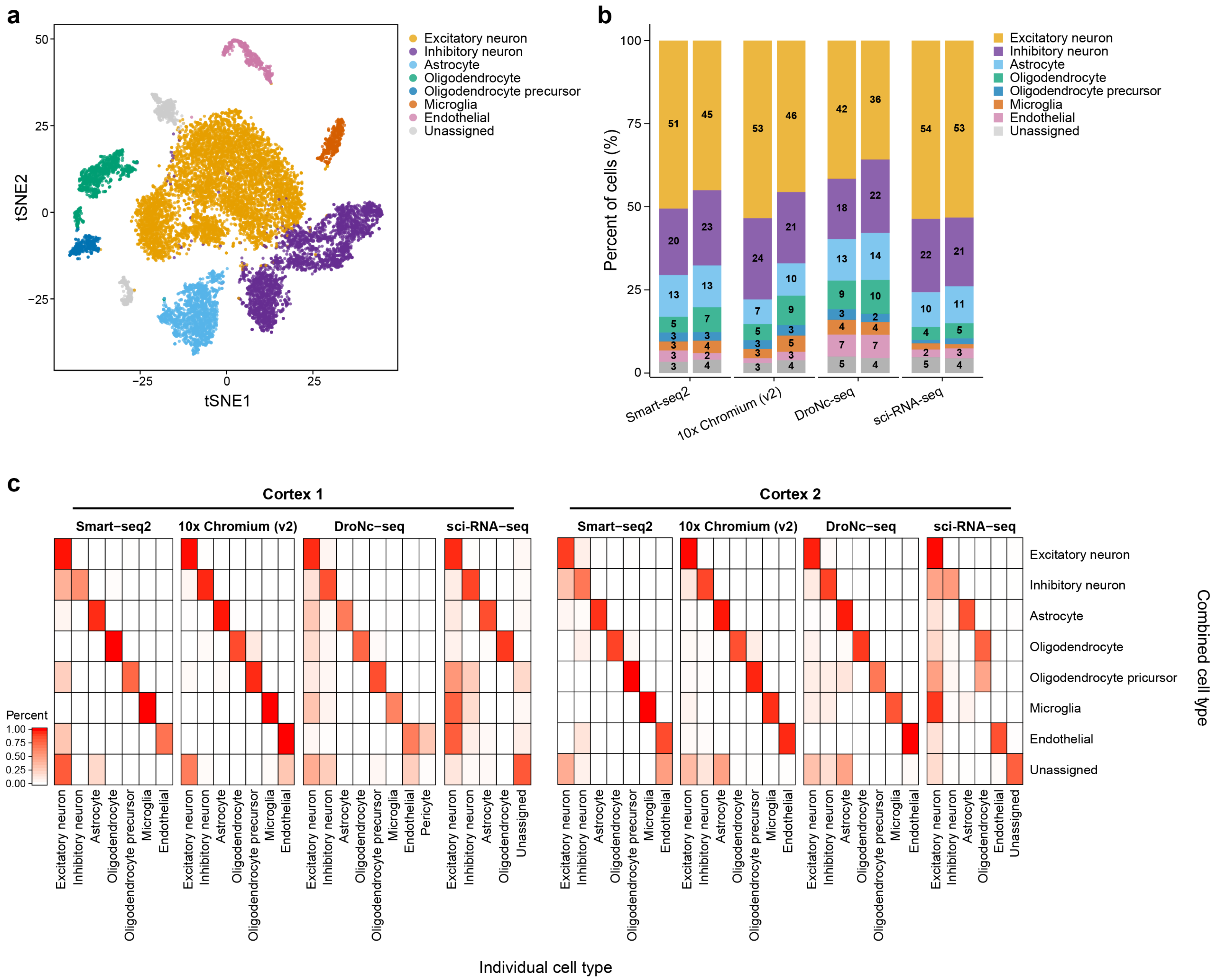

a
tSNE1
tSNE2
Excitatory neuron
Inhibitory neuron
Astrocyte
Oligodendrocyte
Oligodendrocyte precursor
Microglia
Endothelial
Unassigned
b
Percent of cells (%)
Smart-seq2
10x Chromium (v2)
DroNc-seq
sci-RNA-seq
c
Cortex 1
Cortex 2
Percent
Combined cell type
Oligodendrocyte pricursor
Pericyte
Individual cell type

### Supp. Fig. 15

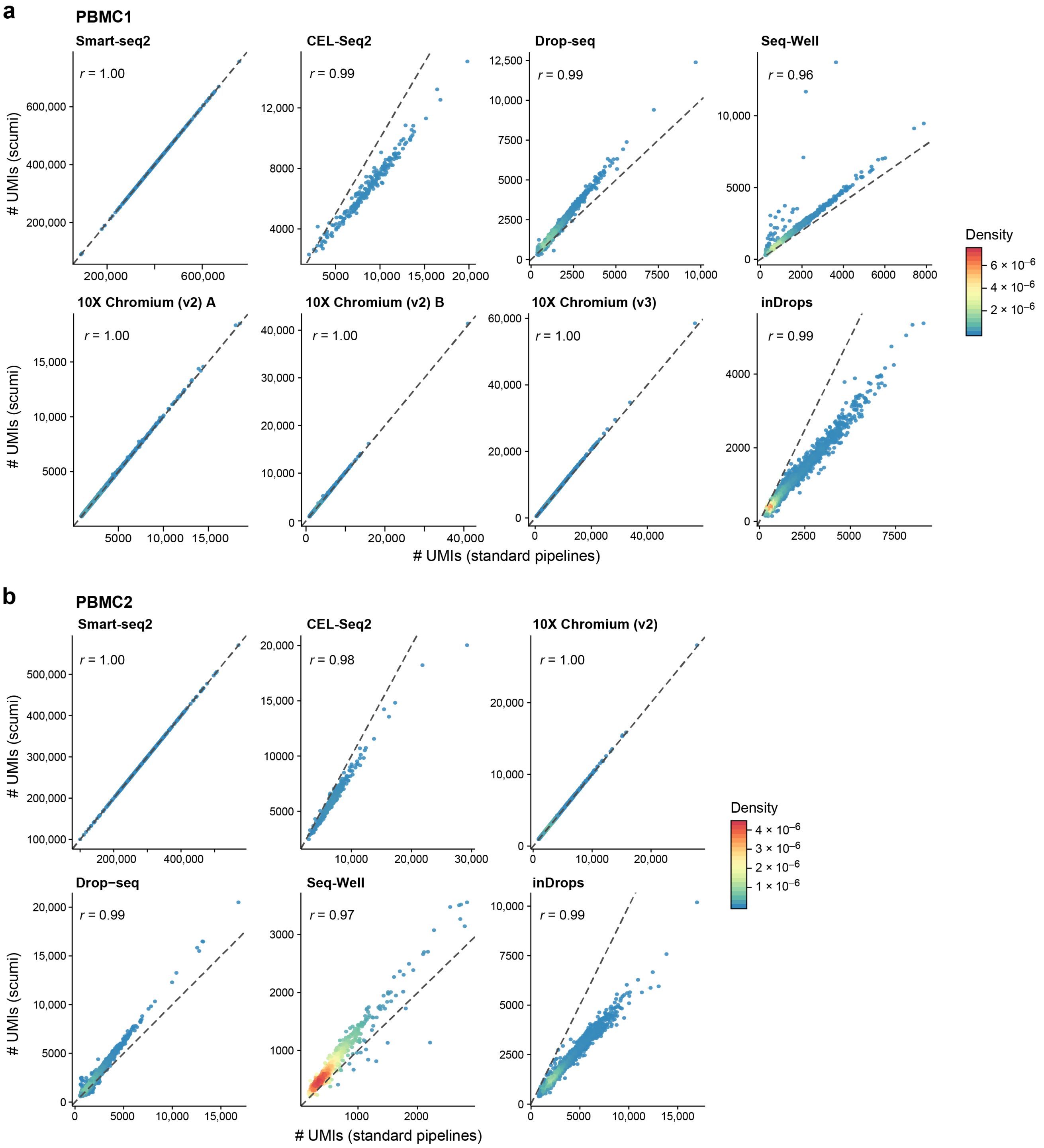
a
PBMC1
Smart-seq2
r = 1.00
CEL-Seq2
r = 0.99
Drop-seq
r = 0.99
Seq-Well
r = 0.96
10X Chromium (v2) A
r = 1.00
10X Chromium (v2) B
r = 1.00
10X Chromium (v3)
r = 1.00
inDrops
r = 0.99
### UMIs (scumi)
### UMIs (standard pipelines)
Density
b
PBMC2
Smart-seq2
r = 1.00
CEL-Seq2
r = 0.98
10X Chromium (v2)
r = 1.00
Drop−seq
r = 0.99
Seq-Well
r = 0.97
inDrops
r = 0.99
### UMIs (scumi)
### UMIs (standard pipelines)
Density

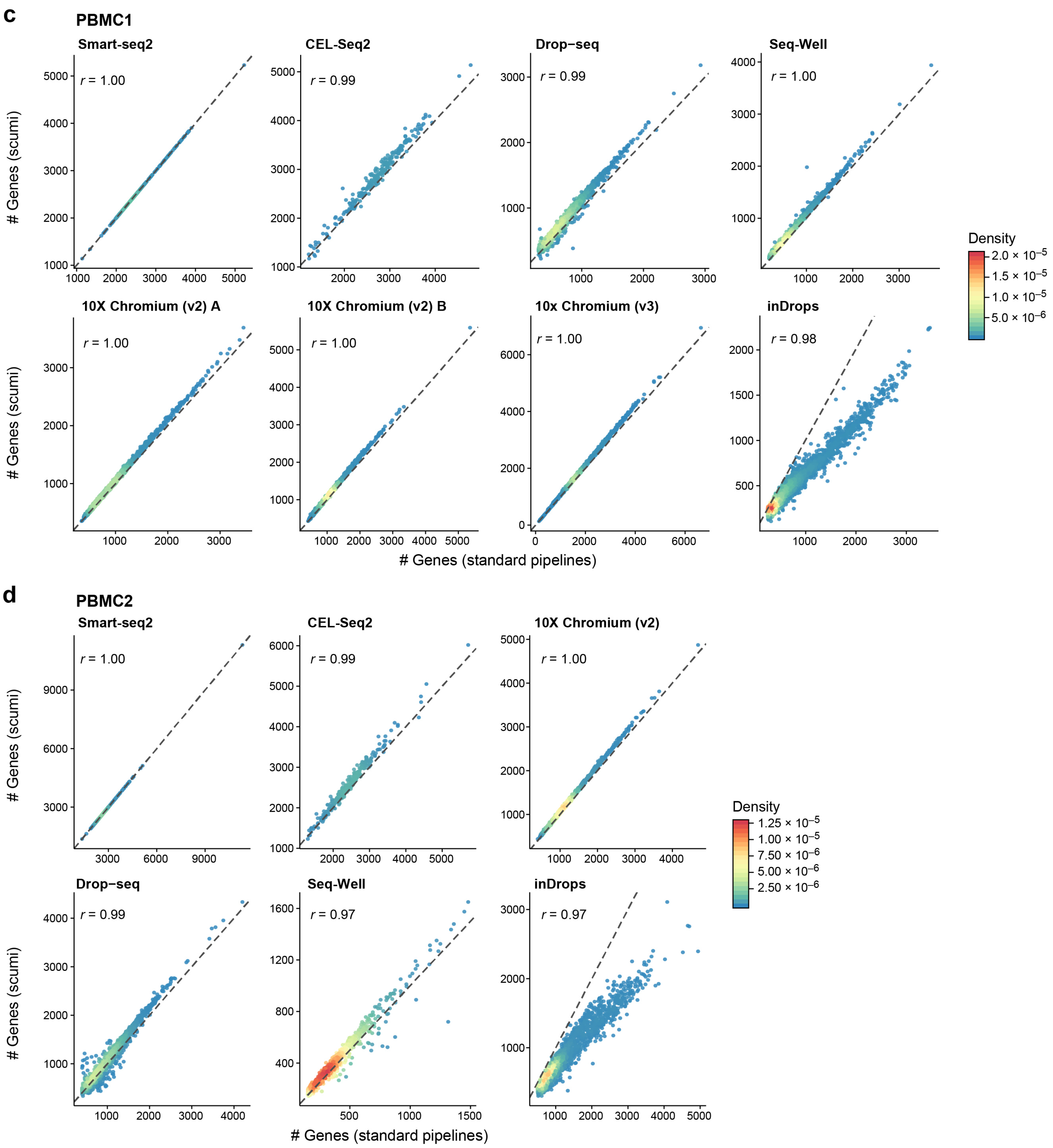
c
PBMC1
Smart-seq2
r = 1.00
CEL-Seq2
r = 0.99
Drop−seq
r = 0.99
Seq-Well
r = 1.00
10X Chromium (v2) A
r = 1.00
10X Chromium (v2) B
r = 1.00
10x Chromium (v3)
r = 1.00
inDrops
r = 0.98
### Genes (scumi)
### Genes (standard pipelines)
Density
2.0 × 10−5
1.5 × 10−5
1.0 × 10−5
5.0 × 10−6
d
PBMC2
Smart-seq2
r = 1.00
CEL-Seq2
r = 0.99
10X Chromium (v2)
r = 1.00
Drop−seq
r = 0.99
Seq-Well
r = 0.97
inDrops
r = 0.97
### Genes (scumi)
### Genes (standard pipelines)
Density
1.25 × 10−5
1.00 × 10−5
7.50 × 10−6
5.00 × 10−6
2.50 × 10−6
