## Supplementary Tables for "Systematic comparative analysis of single cell RNA-sequencing methods"

Table s1

| Method | Replicate | Total # reads | # reads with poly (T) | % reads without poly (T) | # cells | Total # reads / cell | # human cells | # mouse cells | human / mouse cell ratio | Median # human genes | Median # human UMIs | Median # mouse genes | Median # mouse UMIs | Multiplet ratio | |
|---|---|---|---|---|---|---|---|---|---|---|---|---|---|---|---|
| Smart-seq2 | Mixture 1 | 326,789,097 | NA | NA | 342 | 955,524 | 110 | 230 | 0.48 | 7,325 | NA | 6,230 | NA | 0.6% | . |
| CEL-Seq2 | Mixture 1 | 313,910,931 | 305,042,735 | 2.8% | 359 | 874,404 | 101 | 255 | 0.40 | 8,852 | 52,928 | 6,819 | 34,291 | 0.8% | |
| 10x Chromium (v2) | Mixture 1 | 385,003,972 | 381,578,888 | 0.9% | 3,159 | 121,875 | 1,336 | 1,714 | 0.78 | 4,886 | 23,445 | 2,973 | 9,510 | 3.5% | |
| Drop-seq | Mixture 1 | 294,051,172 | 274,892,045 | 6.5% | 2,594 | 113,358 | 1,333 | 1,175 | 1.13 | 2,701 | 5,503 | 1,749 | 3,075 | 3.3% | |
| Seq-Well | Mixture 1 | 241,611,646 | 215,415,794 | 10.8% | 1,627 | 148,501 | 797 | 788 | 1.01 | 4,399 | 12,539 | 3,255 | 8,385 | 2.6% | |
| inDrops | Mixture 1 | 275,471,818 | 142,447,608 | 38.1%* | 3,081 | 89,410 | 1,472 | 1,361 | 1.08 | 1,632 | 3,011 | 1,263 | 2,177 | 8.0% | |
| sci-RNA-seq | Mixture 1 | 302,408,305 | 284,775,866 | 5.8% | 299 | 1,011,399 | 141 | 151 | 0.93 | 4,555 | 11,490 | 2,932 | 6,677 | 2.3% | |
| Smart-seq2 | Mixture 2 | 328,694,640 | NA | NA | 343 | 958,293 | 187 | 156 | 1.20 | 8,729 | NA | 8,476 | NA | 0.0% | |
| CEL-Seq2 | Mixture 2 | 360,590,379 | 348,088,920 | 3.5% | 347 | 1,039,165 | 196 | 149 | 1.32 | 8,334 | 43,683 | 7,786 | 52,892 | 0.6% | |
| 10x Chromium (v2) | Mixture 2 | 354,232,761 | 351,169,692 | 0.9% | 3,187 | 111,149 | 1,929 | 1,221 | 1.58 | 4,940 | 23,027 | 3,905 | 13,702 | 1.2% | |
| Drop-seq | Mixture 2 | 287,143,635 | 265,627,539 | 7.5% | 3,812 | 75,326 | 2,522 | 1,155 | 2.18 | 2,505 | 5,100 | 1,938 | 3,373 | 3.5% | |
| Seq-Well | Mixture 2 | 287,200,476 | 239,060,154 | 16.8% | 1,012 | 283,795 | 547 | 433 | 1.26 | 4,543 | 14,338 | 3,569 | 12,033 | 3.2% | |
| inDrops | Mixture 2 | 133,347,729 | 110,816,672 | 12.1%* | 2,529 | 52,727 | 1,713 | 729 | 2.35 | 2,096 | 3,870 | 2,092 | 4,258 | 3.4% | |
| sci-RNA-seq | Mixture 2 | 459,027,161 | 448,663,261 | 2.3% | 5,023 | 91,385 | 3,135 | 1,733 | 1.81 | 4,593 | 11,025 | 3,401 | 8,256 | 3.1% | |

*For inDrops, 230,135,105 & 126,031,619 reads matched the known sample barcodes in Mixture1 & 2, respectively. Only these reads were used to calculate % reads without poly (T).

All results with scumi computational pipeline.

Table s2

| Method | Replicate | Total # reads | # reads with poly (T) | % reads without poly (T) | # cells | Total # reads/cell | Median # UMIs | Median # genes | | # cells* | Median # UMIs* | Median # genes* | Threshold (genes/cell)* |
|---|---|---|---|---|---|---|---|---|---|---|---|---|---|
| Smart-seq2 | PBMC 1 | 375,056,049 | NA | NA | 311 | 1,205,968 | NA | 2,406 | | 311 | NA | 2,406 | 1,000 |
| CEL-Seq2 | PBMC 1 | 384,742,228 | 370,155,021 | 3.8% | 257 | 1,497,051 | 6,383 | 2,798 | | 250 | 8,470 | 2,696 | 1,200 |
| 10x Chromium (v2) A | PBMC 1 | 523,224,661 | 519,300,975 | 0.7% | 5,172 | 101,165 | 2,029 | 833 | | 4,368 | 2,309 | 864 | none |
| 10x Chromium (v2) B | PBMC 1 | 687,784,556 | 679,229,257 | 1.2% | 3,057 | 224,987 | 3,050 | 1,115 | | 3,035 | 2,978 | 1,042 | none |
| 10x Chromium (v3) | PBMC 1 | 666,523,933 | 660,891,029 | 0.8% | 4,033 | 165,268 | 4,926 | 1,622 | | 3,795 | 4,851 | 1,555 | none |
| Drop-seq | PBMC 1 | 487,402,716 | 414,658,200 | 14.9% | 4,683 | 104,079 | 1,199 | 679 | | 4,864 | 938 | 580 | 300 |
| Seq-Well | PBMC 1 | 235,605,948 | NA | NA | 5,125 | 45,972 | 844 | 513 | | 5,006 | 696 | 471 | 200 |
| inDrops | PBMC 1 | 427,146,724 | 325,157,955 | 23.9% | 6,184 | 69,073 | 366 | 256 | | 5,946 | 599 | 373 | 250 |
| Smart-seq2 | PMBC 2 | 250,081,663 | NA | NA | 273 | 916,050 | NA | 2,632 | | 303 | NA | 2,589 | 1,000 |
| CEL-Seq2 | PMBC 2 | 278,052,915 | 271,465,989 | 2.4% | 307 | 905,710 | 5,787 | 2,545 | | 299 | 6,685 | 2,447 | 1,200 |
| 10x Chromium (v2) | PMBC 2 | 319,398,613 | 312,697,214 | 2.1% | 3,362 | 95,003 | 2,627 | 1,110 | | 3,188 | 2,627 | 1,060 | none |
| Drop-seq | PMBC 2 | 703,840,624 | 623,619,962 | 11.4% | 6,412 | 109,769 | 1,851 | 911 | | 6,323 | 1,483 | 818 | 400 |
| Seq-Well | PMBC 2 | 176,233,532 | 103,664,673 | 41.2% | 913 | 193,027 | 519 | 323 | | 1,087 | 363 | 280 | 150 |
| inDrops | PMBC 2 | 502,795,162 | 430,102,035 | 14.5% | 4,526 | 111,090 | 1,243 | 611 | | 4,708 | 2,021 | 782 | 500 |

All results are from scumi computational pipeline, except those with an * are from standard, method-specific pipelines.

For standard pipelines, only cells above the threshold were included.

NA = Not applicable

Table s3

| Method | Replicate | Total # reads | # reads with poly (T) | % reads without poly (T) | # cells | Total # reads / cell | Median # UMIs | Median # genes |
|---|---|---|---|---|---|---|---|---|
| Smart-seq2 | Cortex 1 | 478,693,321 | NA | NA | 295 | 1,622,689 | NA | 5,774 |
| 10x Chromium (v2) | Cortex 1 | 561,087,440 | 536,520,270 | 4.4% | 1,480 | 379,113 | 6,994 | 3,221 |
| DroNc-seq | Cortex 1 | 216,550,548 | 185,384,312 | 14.4% | 2,195 | 98,656 | 2,092 | 1,401 |
| sci-RNA-seq | Cortex 1 | 315,124,882 | 300,100,752 | 4.8% | 1,886 | 167,086 | 3,524 | 1,591 |
| Smart-seq2 | Cortex 2 | 570,270,279 | NA | NA | 349 | 1,634,012 | NA | 5,014 |
| 10x Chromium (v2) | Cortex 2 | 590,837,576 | 567,439,605 | 4.0% | 4,091 | 144,424 | 3,527 | 1,931 |
| DroNc-seq | Cortex 2 | 250,573,006 | 194,428,927 | 22.4% | 935 | 267,993 | 3,094 | 1,879 |
| sci-RNA-seq | Cortex 2 | 382,131,662 | 367,386,555 | 3.9% | 3,944 | 96,889 | 1,895 | 1,163 |
| All results with scumi computational pipeline. | | | | | | | | |

Table s4

| Method | Experiment | Additional reads | Number Cells | Additional reads per cell | Average reads per cell | Increase in reads per cell |
|---|---|---|---|---|---|---|
| Smart-seq2 | PBMC1 | 7,291,199 | 311 | 23,444 | 1,205,968 | 1.94% |
| Smart-seq2 | PBMC2 | 4,508,783 | 273 | 16,516 | 916,050 | 1.80% |
| CEL-Seq2 | PBMC1 | 1,828,337 | 257 | 7,114 | 1,197,690 | 0.59% |
| CEL-Seq2 | PBMC2 | 1,113,041 | 307 | 3,626 | 905,710 | 0.40% |
| 10X Chromium (v2) A | PBMC1 | 2,810,044 | 5,184 | 542 | 68,635 | 0.79% |
| 10X Chromium (v2) B | PBMC1 | 1,714,342 | 3,222 | 532 | 66,173 | 0.80% |
| 10X Chromium (v3) | PBMC1 | 2,577,057 | 4,027 | 640 | 69,516 | 0.92% |
| 10X Chromium (v2) | PBMC2 | 2,138,732 | 3,362 | 636 | 95,003 | 0.67% |
| Drop-seq | PBMC1 | 2,846,555 | 4,640 | 613 | 69,331 | 0.88% |
| Drop-seq | PBMC2 | 5,754,844 | 6,412 | 898 | 95,501 | 0.94% |
| Seq-Well | PBMC1 | 2,948,993 | 5,125 | 575 | 45,972 | 1.25% |
| Seq-Well | PBMC2 | 415,842 | 551 | 755 | 156,699 | 0.48% |
| inDrops | PBMC1 | 4,559,763 | 6,184 | 737 | 69,073 | 1.07% |
| inDrops | PBMC2 | 4,086,940 | 5,166 | 791 | 83,705 | 0.95% |

Table s5

| Experiment | Method | # Cells (after sampling) | # Cells (before sampling) | Average reads per cell (Using # cells after sampling) | Average reads per cell (Using # cells before sampling) | Total Reads (after sampling) | Notes (read sampling) | # cells (after sampling cells) |
|---|---|---|---|---|---|---|---|---|
| Mixture1 | Smart-seq2 | 342 | 342 | 879,194 | 879,194 | 300,684,518 | | NA |
| Mixture1 | CEL-Seq2 | 359 | 359 | 874,404 | 874,404 | 313,910,931 | Sampled other to this one | NA |
| Mixture1 | 10X Chromium | 3,162 | 3,159 | 88,883 | 88,968 | 281,049,304 | | NA |
| Mixture1 | Drop-seq | 2,594 | 2,594 | 89,551 | 89,551 | 232,296,106 | | NA |
| Mixture1 | Seq-Well | 1,630 | 1,627 | 88,936 | 89,100 | 144,965,823 | | NA |
| Mixture1 | inDrops | 3,081 | 3,081 | 89,410 | 89,410 | 275,471,818 | Sampled others to this one | NA |
| Mixture1 | sci-RNA-seq | 300 | 299 | 90,703 | 91,006 | 27,210,842 | | NA |
| Mixture2 | Smart-seq2 | 343 | 343 | 958,293 | 958,293 | 328,694,640 | Sampled other to this one | NA |
| Mixture2 | CEL-Seq2 | 347 | 347 | 956,044 | 956,044 | 331,747,198 | | NA |
| Mixture2 | 10X Chromium | 3,186 | 3,187 | 52,254 | 52,238 | 166,480,959 | | NA |
| Mixture2 | Drop-seq | 3,830 | 3,812 | 52,480 | 52,728 | 201,000,212 | | NA |
| Mixture2 | Seq-Well | 1,015 | 1,012 | 53,756 | 53,916 | 54,562,643 | | NA |
| Mixture2 | inDrops | 2,529 | 2,529 | 52,727 | 52,727 | 133,347,729 | Sampled others to this one | NA |
| Mixture2 | sci-RNA-seq | 5,026 | 5,023 | 52,982 | 53,013 | 266,286,458 | | NA |
| PBMC1 | Smart-seq2 | 311 | 311 | 1,205,968 | 1,205,968 | 375,056,048 | Sampled other to this one | 253 |
| PBMC1 | CEL-Seq2 | 253 | 257 | 1,216,626 | 1,197,690 | 307,806,288 | | 253 |
| PBMC1 | 10X Chromium (v2) A | 5,184 | 5,172 | 68,635 | 68,794 | 355,803,563 | | 3,222 |
| PBMC1 | 10X Chromium (v2) B | 3,222 | 3,057 | 66,173 | 69,744 | 213,207,949 | | 3,222 |
| PBMC1 | 10X Chromium (v3) | 4,027 | 4,033 | 69,516 | 69,413 | 279,941,482 | | 3,222 |
| PBMC1 | Drop-seq | 4,640 | 4,683 | 69,331 | 68,694 | 321,693,781 | | 3,222 |
| PBMC1 | Seq-Well | 5,125 | 5,125 | 45,972 | 45,972 | 235,605,948 | Used all reads | 3,222 |
| PBMC1 | inDrops | 6,184 | 6,184 | 69,073 | 69,073 | 427,147,432 | Sampled others to this one | 3,222 |
| PBMC2 | Smart-seq2 | 273 | 273 | 916,050 | 916,050 | 250,081,650 | Used all reads | 273 |
| PBMC2 | CEL-Seq2 | 307 | 307 | 905,710 | 905,710 | 278,052,970 | Used all reads | 273 |
| PBMC2 | 10X Chromium | 3,362 | 3,362 | 95,003 | 95,003 | 319,400,086 | Sampled others to this one | 3,362 |
| PBMC2 | Drop-seq | 6,412 | 6,412 | 95,501 | 95,501 | 612,349,900 | | 3,362 |
| PBMC2 | Seq-Well | 551 | 913 | 156,699 | 94,569 | 86,341,083 | | 551 |
| PBMC2 | inDrops | 5,166 | 4,526 | 83,705 | 95,541 | 432,418,556 | | 3,362 |
| Cortex1 | Smart-seq2 | 295 | 295 | 1,622,689 | 1,622,689 | 478,693,321 | Used all reads | 295 |
| Cortex1 | 10X Chromium | 1,453 | 1,480 | 100,404 | 98,572 | 145,886,949 | | 1,453 |
| Cortex1 | DroNc-seq | 2,195 | 2,195 | 98,656 | 98,656 | 216,550,548 | Sampled others to this one | 1,453 |
| Cortex1 | sci-RNA-seq | 1,848 | 1,886 | 100,617 | 98,589 | 185,939,664 | | 1,453 |
| Cortex2 | Smart-seq2 | 349 | 349 | 1,634,012 | 1,634,012 | 570,270,279 | Used all reads | 349 |
| Cortex2 | 10X Chromium | 4,267 | 4,091 | 92,775 | 96,766 | 395,869,815 | | 3,944 |
| Cortex2 | DroNc-seq | 892 | 935 | 101,130 | 96,479 | 90,208,024 | | 892 |
| Cortex2 | sci-RNA-seq | 3,944 | 3,944 | 96,889 | 96,889 | 382,131,662 | Sampled others to this one | 3,944 |
| NA: not applicable | | | | | | | | |

Table s6

| Expt. | Method | Cell Type (Harmony) | Cell Type (Cells separated by method) | # cells in cluster | % cells in cluster for this expt. |
|---|---|---|---|---|---|
| PBMC1 | CEL-Seq2 | Plasmacytoid dendritic cell | B cell | 2 | 0.79% |
| PBMC2 | Smart-seq2 | Megakaryocyte | B cell | 4 | 1.47% |
| PBMC2 | CEL-Seq2 | Megakaryocyte | $CD4^+$ T cell | 5 | 1.83% |
| PBMC2 | CEL-Seq2 | Plasmacytoid dendritic cell | B cell | 3 | 1.10% |
| PBMC2 | Drop-seq | Megakaryocyte | $CD4^+$ T cell | 63 | 1.87% |

Table s7

| Method | Cost/cell | | # Cells | Time (hours) |
|---|---|---|---|---|
| Smart-seq2 | $ | 10.59 | 384 | 25.67 |
| CEL-Seq2 | $ | 3.56 | 384 | 25.17 |
| 10X Chromium (v2) | $ | 0.32 | 4,000 | 9.00 |
| 10X Chromium (v3) | $ | 0.33 | 4,000 | 9.00 |
| Drop-Seq/DroNc-Seq | $ | 0.10 | 6,000 | 10.00 |
| Seq-Well | $ | 0.09 | 2,500 | 10.17 |
| inDrops | $ | 0.07 | 3,000 | 24.00 |
| sci-RNA-seq | $ | 0.28 | 7,680 | 17.42 |
| Time is for entire process | | | | |
| FACS costs not included for plate-based methods or for nuclei | | | | |

Table s8

| RNA-Seq Method | Experiment | Concentration (cells/ml) | Total Number of Cells | Resuspension Buffer | NIH3T3:HEK293 Mixed at 1:1 Ratio? |
|---|---|---|---|---|---|
| Smart-seq2 | Mixture1 & Mixture2 for sorting | 1,000,000 | 1,000,000 | 1X PBS | Yes |
| CEL-Seq2 | | | | | |
| 10x Chromium (v2) | Mixture1 & Mixture2 | 1,000,000 | 10,000 | 1X PBS + 0.04% BSA | Yes |
| Drop-Seq | Mixture1 & Mixture2 | 100,000 | 400,000 | 1X PBS + 0.01% BSA | Yes |
| Seq-Well | Mixture1 & Mixture2 | 50,000 | 50,000 | RPMI + 10% FBS | Yes |
| inDrops | Mixture1 & Mixture2 | 100,000 | 50,000 | 1X PBS + 15% OptiPrep | Yes |
| sci-RNA-seq | Mixture1 & Mixture2 | 1,000,000 | 2,500,000 | 1X PBS | Yes |
| TruSeq (bulk) | Mixture1 & Mixture2 | | 5,000,000 | DNA/RNA Shield | No, each separate |
| Smart-seq2 | PBMC1 for sorting | 890,000 | 1,000,000 | RPMI (without phenol) + 2% human serum | NA |
| CEL-Seq2 | | | | | |
| Smart-seq2 | PBMC2 for sorting | 2,100,000 | 1,800,000 | RPMI (without phenol) + 2% human serum | NA |
| CEL-Seq2 | | | | | |
| 10x Chromium (v2) | PBMC1 (A) & PBMC2 | 1,000,000 | 10,000 | 1X PBS + 0.04% BSA | NA |
| Drop-Seq | PBMC1 & PBMC2 | 100,000 | 400,000 | 1X PBS + 0.01% BSA | NA |
| Seq-Well | PBMC1 & PBMC2 | 50,000 | 50,000 | RPMI + 10% FBS | NA |
| inDrops | PBMC1 & PBMC2 | 100,000 | 50,000 | 1X PBS + 15% OptiPrep | NA |
| sci-RNA-seq | PBMC1 & PBMC2 | 1,000,000 | 2,500,000 | 1X PBS | NA |
| 10x Chromium (v2) and (v3) | PBMC1 (B) | 1,000,000 | 20,000 | 1X PBS + 0.04% BSA | NA |
| TruSeq (bulk) | PBMC1 | | 1,000,000 | DNA/RNA Shield | NA |
| TruSeq (bulk) | PBMC2 | | 1,800,000 | DNA/RNA Shield | NA |
| Smart-seq2 | Cortex1 for sorting | 7,000,000 | 6,510,000 | 1X PBS + 1% BSA | NA |
| 10x Chromium (v2) | | | | | |
| Smart-seq2 | Cortex2 for sorting | 5,400,000 | 4,590,000 | 1X PBS + 1% BSA | NA |
| 10x Chromium (v2) | | | | | |
| DroNc-Seq | Cortex1 | 300,000 | 750,000 | 1X PBS + 0.01% BSA | NA |
| DroNc-Seq | Cortex2 | 360,000 | 900,000 | 1X PBS + 0.01% BSA | NA |
| sci-RNA-seq | Cortex1 | 300,000 | 450,000 | 1X PBS + 1% BSA | NA |
| sci-RNA-seq | Cortex2 | 540,000 | 810,000 | 1X PBS + 1% BSA | NA |
| TruSeq (bulk) | Cortex1 | | 1,000,000 | DNA/RNA Shield | NA |
| TruSeq (bulk) | Cortex2 | | 720,000 | DNA/RNA Shield | NA |
| NA: Not applicable | | | | | |
| We list the number of cells actually used, although in many cases, we may not have needed all of the cells to complete the experiment. | | | | | |

Table s9

| Experiment # | Cell Source | Date Thawed | RNA-Seq Date | Time in Culture from Thaw -> RNA-Seq | Total time in culture | Passage Number |
|---|---|---|---|---|---|---|
| 1 | fresh frozen NIH3T3 & HEK293 ATCC vials | 6/9/2017 | 6/26/2017 | 18 days | 18 days | Both P4 (thawed fresh ATCC cells - considered as P0) |
| 2 | 2x NIH3T3 & HEK293 vials (2.5e6 cells, both P3) frozen on 6/17/17 (after 9 days in culture) | 10/19/2017 | 10/31/2017 | 13 days | 22 days | NIH3T3: P7; HEK293: P6 |

Table s10

| Sequencing Platform | Flow Cell | Sample | Library Types | Number of lanes | Read 1 (bases) | Read 2 (bases) | Index 1 (bases) | Index 2 (bases) | Custom Sequencing Primer | PhiX added |
|---|---|---|---|---|---|---|---|---|---|---|
| HiSeq2500 high capacity | CAPTEANXX | Mixture1 | Smart-seq2, CEL-Seq2, 10x (v2), Drop-Seq, Seq-Well, sci-RNA-Seq | 7 | 50 | 50 | 10 | 10 | for Read 1 | none |
| HiSeq2500 high capacity | CAPTEANXX | Mixture1 | inDrops | 1 | 50 | 50 | 10 | 10 | | none |
| HiSeq2500 high capacity | CBU1CANXX | Mixture2 | Smart-seq2, CEL-Seq2, 10x (v2), Drop-Seq, Seq-Well, sci-RNA-Seq | 8 | 50 | 50 | 10 | 10 | for Read 1 | none |
| HiSeq2500 rapid run | HYK72BCXY | Mixture2 | inDrops | 1 | 100 | 94 | 8 | 8 | | 5% |
| HiSeq2500 rapid run | H5VV5BCX2 | Mixed2 | Seq-Well | 1 | 100 | 94 | 8 | 8 | for Read 1 | 5% |
| HiSeq2500 high capacity | CC7W2ANXX | PBMC1 | Smart-seq2, CEL-Seq2, 10x (v2), Drop-Seq, sci-RNA-Seq | 8 | 50 | 50 | 10 | 10 | for Read 1 | none |
| HiSeq2500 rapid run | H5VV5BCX2 | PBMC1 | inDrops | 1 | 100 | 94 | 8 | 8 | | 5% |
| HiSeq2500 high capacity | CC86JANXX | PBMC2 | Smart-seq2, CEL-Seq2, 10x (v2), Drop-Seq, Seq-Well, sci-RNA-Seq | 7 | 50 | 50 | 10 | 10 | for Read 1 | none |
| HiSeq2500 high capacity | CC86JANXX | PBMC2 | inDrops | 1 | 50 | 50 | 10 | 10 | | 5% |
| HiSeq2500 high capacity | CCJ15ANXX | Cortex1 | Smart-seq2, 10x (v2), sci-RNA-Seq | 4 | 50 | 50 | 10 | 10 | | none |
| HiSeq2500 high capacity | CCJ15ANXX | Cortex2 | Smart-seq2, 10x (v2), sci-RNA-Seq | 4 | 50 | 50 | 10 | 10 | | none |
| HiSeq2500 high capacity | CCKVLANXX | Cortex1 | Smart-seq2, 10x (v2), DroNc-Seq, sci-RNA-Seq | 4 | 50 | 50 | 10 | 10 | for Read 1 | none |
| HiSeq2500 high capacity | CCKVLANXX | Cortex2 | Smart-seq2, 10x (v2), DroNc-Seq, sci-RNA-Seq | 4 | 50 | 50 | 10 | 10 | for Read 1 | none |
| HiSeq2500 rapid run | HJTJGBCX2 | Mixture1, Mixture2, PBMC1, PBMC2 | inDrops | 2 | 50 | 19 | 8 | 8 | | 10% |
| HiSeq2500 rapid run | HKGWKBCX2 | PBMC1 | Seq-Well | 2 | 20 | 50 | 8 | 0 | for Read 1 | |
| HiSeq2500 rapid run | HKGWFBCX2 | PBMC1, PBMC2 | inDrops | 2 | 50 | 19 | 8 | 8 | | 10% |
| HiSeq2500 rapid run | HKH3FBCX2 | PBMC1, PBMC2 | inDrops | 2 | 50 | 19 | 8 | 8 | | 10% |
| HiSeq2500 high capacity | CCLBDANXX | PBMC2 | Drop-Seq | 2 | 31 | 50 | 8 | 0 | for Read 1 | 10% |
| HiSeq2500 high capacity | CCLBDANXX | PBMC1, Cortex1 | Drop-Seq (P1), Seq-Well (P1), DroNc-Seq (C1) | 3 | 31 | 50 | 8 | 0 | for Read 1 | 10% |
| HiSeq2500 high capacity | CCLBDANXX | PBMC1, PBMC2, Cortex2 | 10x (v2) (P1), Drop-Seq (P1&P2), CEL-Seq2 (P2), DroNc-Seq (C2) | 3 | 31 | 50 | 8 | 0 | for Read 1 | 10% |
| HiSeq2500 high capacity | CD370ANXX | PBMC1 | 10x (v2), 10x (v3) | 8 | 33 | 50 | 8 | 0 | for Read 1 | 10% |
| Other libraries not included in this study were also sequenced in some lanes. | | | | | | | | | | |

Table s11

| Library Types | Flow Cell(s) | Sample | Read 1 length | Read 2 length | Index 1 length | Index 2 length | Read 1 trimmed | Read 2 trimmed | Index 1 trimmed | Index 2 trimmed | In-line index position | UMI position | PolyT start position | Cell barcodes |
|---|---|---|---|---|---|---|---|---|---|---|---|---|---|---|
| Smart-seq2 | CAPTEANXX, CBU1CANXX, CC7W2ANXX, CC86JANXX, CCKVLANXX, CCJ15ANXX | Mixture1, Mixture2, PBMC1, PBMC2, Cortex1, Cortex2 | 50 | 50 | 10 | 10 | 0 | 0 | 2 | 2 | none | none | none | index 1, index 2 |
| CEL-Seq2 | CAPTEANXX, CBU1CANXX, CC7W2ANXX, CC86JANXX | Mixture1, Mixture2, PBMC1, PBMC2 | 50 | 50 | 10 | 10 | 38 | 0 | 2 | 10 | Read 1, #7-12 | Read 1, #1-6 | Read 1, #13 | In-line |
| CEL-Seq2 | CCLBDANXX | PBMC2 | 31 | 50 | 8 | 0 | 19 | 0 | 0 | 0 | Read 1, #7-12 | Read 1, #1-6 | Read 1, #13 | In-line |
| 10x Chromium (v2) | CAPTEANXX, CBU1CANXX, CC7W2ANXX, CC86JANXX, CCKVLANXX, CCJ15ANXX | Mixture1, Mixture2, PBMC1, PBMC2, Cortex1, Cortex2 | 50 | 50 | 10 | 10 | 24 | 0 | 2 | 10 | Read 1, #1-16 | Read 1, #17-26 | Read 1, #27 | In-line |
| 10x Chromium (v2) | CCLBDANXX | PBMC1, PBMC2 | 31 | 50 | 8 | 0 | 5 | 0 | 0 | 0 | Read 1, #1-16 | Read 1, #17-26 | Read 1, #27 | In-line |
| 10x Chromium (v2) | CD370ANXX | PBMC1 | 33 | 50 | 8 | 0 | 7 | 0 | 0 | 0 | Read 1, #1-16 | Read 1, #17-26 | Read 1, #27 | In-line |
| 10x Chromium (v3) | CD370ANXX | PBMC1 | 33 | 50 | 8 | 0 | 5 | 0 | 0 | 0 | Read 1, #1-16 | Read 1, #17-28 | Read 1, #29 | In-line |
| Drop-seq | CAPTEANXX, CBU1CANXX, CC7W2ANXX, CC86JANXX | Mixture1, Mixture2, PBMC1, PBMC2 | 50 | 50 | 10 | 10 | 30 | 0 | 2 | 10 | Read 1, #1-12 | Read 1, #13-20 | Read 1, #21 | In-line |
| Drop-seq | CCLBDANXX | PBMC1, PBMC2 | 31 | 50 | 8 | 0 | 11 | 0 | 0 | 0 | Read 1, #1-12 | Read 1, #13-20 | Read 1, #21 | In-line |
| DroNc-seq | CCKVLANXX | Cortex1, Cortex2 | 50 | 50 | 10 | 10 | 30 | 0 | 2 | 10 | Read 1, #1-12 | Read 1, #13-20 | Read 1, #21 | In-line |
| DroNc-seq | CCLBDANXX | Cortex1, Cortex2 | 31 | 50 | 8 | 0 | 11 | 0 | 0 | 0 | Read 1, #1-12 | Read 1, #13-20 | Read 1, #21 | In-line |
| Seq-Well | CAPTEANXX, CBU1CANXX | Mixture1, Mixture2 | 50 | 50 | 10 | 10 | 30 | 0 | 2 | 10 | Read 1, #1-12 | Read 1, #13-20 | Read 1, #21 | In-line |
| Seq-Well | H5VV5BCX2 | Mixture2 | 100 | 94 | 8 | 8 | 80 | 0 | 0 | 8 | Read 1, #1-12 | Read 1, #13-20 | Read 1, #21 | In-line |
| Seq-Well | HKGWKBCX2 | PBMC1 | 20 | 50 | 8 | 0 | 0 | 0 | 0 | 0 | Read 1, #1-12 | Read 1, #13-20 | Not sequenced | In-line |
| Seq-Well | CCLBDANXX | PBMC1 | 31 | 50 | 8 | 0 | 11 | 0 | 0 | 0 | Read 1, #1-12 | Read 1, #13-20 | Read 1, #21 | In-line |
| Seq-Well | CC86JANXX | PBMC2 | 50 | 50 | 10 | 10 | 30 | 0 | 2 | 10 | Read 1, #1-12 | Read 1, #13-20 | Read 1, #21 | In-line |
| inDrops | CAPTEANXX, CC86JANXX | Mixture1, PBMC2 | 50 | 50 | 10 | 10 | 0 | 36 | 2 | 2 | Read 2, #1-8 | Read 2, #9-14 | Read 2, #15 | In-line, index 1, index 2 |
| inDrops | HJTJGBCX2 | Mixture1, Mixture2 | 50 | 19 | 8 | 8 | 0 | 5 | 0 | 0 | Read 2, #1-8 | Read 2, #9-14 | Read 2, #15 | In-line, index 1, index 2 |
| inDrops | HYK72BCXY, H5VV5BCX2 | Mixture2, PBMC1 | 100 | 94 | 8 | 8 | 0 | 80 | 0 | 0 | Read 2, #1-8 | Read 2, #9-14 | Read 2, #15 | In-line, index 1, index 2 |
| inDrops | HJTJGBCX2, HKGWFBCX2, HKH3FBCX2 | PBMC1 | 50 | 19 | 8 | 8 | 0 | 5 | 0 | 0 | Read 2, #1-8 | Read 2, #9-14 | Read 2, #15 | In-line, index 1, index 2 |
| inDrops | HJTJGBCX2, HKGWFBCX2, HKH3FBCX2 | PBMC2 | 50 | 19 | 8 | 8 | 0 | 5 | 0 | 0 | Read 2, #1-8 | Read 2, #9-14 | Read 2, #15 | In-line, index 1, index 2 |
| sci-RNA-seq | CAPTEANXX, CBU1CANXX, CCKVLANXX, CCJ15ANXX | Mixture1, Mixture2, Cortex1, Cortex2 | 50 | 50 | 10 | 10 | 32 | 0 | 0 | 0 | Read 1, #9-18 | Read 1, #1-8 | Read 1, #19 | In-line, index 1, index 2 |

Bases listed for total and trimmed read lengths

sci-RNA-seq reads were demultiplexed such that the cells from the same well in the RT step were in the same fastq file; we therefore only used the in-line RT cell barcodes for down-stream analysis

Table s12

| Cell type | Gene | Positive or Negative |
|---|---|---|
| CD4+ T cell | CD3D | + |
| CD4+ T cell | CD3E | + |
| CD4+ T cell | CD3G | + |
| CD4+ T cell | TRAC | + |
| CD4+ T cell | CD4 | + |
| CD4+ T cell | TCF7 | + |
| CD4+ T cell | CD27 | + |
| CD4+ T cell | IL7R | + |
| CD4+ T cell | CD8A | - |
| CD4+ T cell | CD8B | - |
| CD4+ T cell | GNLY | - |
| CD4+ T cell | NKG7 | - |
| CD4+ T cell | CST7 | - |
| Cytotoxic T cell | CD3D | + |
| Cytotoxic T cell | CD3E | + |
| Cytotoxic T cell | CD3G | + |
| Cytotoxic T cell | TRAC | + |
| Cytotoxic T cell | CD8A | + |
| Cytotoxic T cell | CD8B | + |
| Cytotoxic T cell | GZMK | + |
| Cytotoxic T cell | CCL5 | + |
| Cytotoxic T cell | NKG7 | + |
| Cytotoxic T cell | CD4 | - |
| Cytotoxic T cell | FCER1G | - |
| B cell | CD19 | + |
| B cell | MS4A1 | + |
| B cell | CD79A | + |
| B cell | CD79B | + |
| B cell | MZB1 | + |
| B cell | IGHD | + |
| B cell | IGHM | + |
| Natural killer cell | NCAM1 | + |
| Natural killer cell | NKG7 | + |
| Natural killer cell | KLRB1 | + |
| Natural killer cell | KLRD1 | + |
| Natural killer cell | KLRF1 | + |
| Natural killer cell | KLRC1 | + |
| Natural killer cell | KLRC2 | + |
| Natural killer cell | KLRC3 | + |
| Natural killer cell | KLRC4 | + |
| Natural killer cell | CD3D | - |
| Natural killer cell | CD3E | - |
| Natural killer cell | CD3G | - |
| Natural killer cell | CD14 | - |
| Natural killer cell | FCGR3A | + |
| Natural killer cell | FCGR3B | + |

Table s12

| | | |
|---|---|---|
| Natural killer cell | ITGAL | + |
| Natural killer cell | ITGAM | + |
| Natural killer cell | FCER1G | + |
| Natural killer cell | TRAC | - |
| CD14+ monocyte | VCAN | + |
| CD14+ monocyte | FCN1 | + |
| CD14+ monocyte | S100A8 | + |
| CD14+ monocyte | S100A9 | + |
| CD14+ monocyte | CD14 | + |
| CD14+ monocyte | ITGAL | + |
| CD14+ monocyte | ITGAM | + |
| CD14+ monocyte | CSF3R | + |
| CD14+ monocyte | CSF1R | + |
| CD14+ monocyte | CX3CR1 | + |
| CD14+ monocyte | FCGR3A | - |
| CD14+ monocyte | FCGR3B | - |
| CD14+ monocyte | TYROBP | + |
| CD14+ monocyte | LYZ | + |
| CD14+ monocyte | S100A12 | + |
| CD14+ monocyte | CD3D | - |
| CD14+ monocyte | CD3E | - |
| CD14+ monocyte | CD3G | - |
| CD14+ monocyte | TRAC | - |
| CD14+ monocyte | NKG7 | - |
| CD14+ monocyte | KLRB1 | - |
| CD14+ monocyte | KLRD1 | - |
| CD16+ monocyte | FCN1 | + |
| CD16+ monocyte | FCGR3A | + |
| CD16+ monocyte | FCGR3B | + |
| CD16+ monocyte | ITGAL | + |
| CD16+ monocyte | ITGAM | + |
| CD16+ monocyte | CSF3R | + |
| CD16+ monocyte | CSF1R | + |
| CD16+ monocyte | CX3CR1 | + |
| CD16+ monocyte | CDKN1C | + |
| CD16+ monocyte | MS4A7 | + |
| CD16+ monocyte | S100A8 | - |
| CD16+ monocyte | S100A9 | - |
| CD16+ monocyte | S100A12 | - |
| CD16+ monocyte | CD14 | - |
| CD16+ monocyte | CD3D | - |
| CD16+ monocyte | CD3E | - |
| CD16+ monocyte | CD3G | - |
| CD16+ monocyte | TRAC | - |
| CD16+ monocyte | NKG7 | - |
| CD16+ monocyte | KLRB1 | - |
| CD16+ monocyte | KLRD1 | - |

Table s12

| | | |
|---|---|---|
| Dendritic cell | HLA-DPB1 | + |
| Dendritic cell | HLA-DPA1 | + |
| Dendritic cell | HLA-DQA1 | + |
| Dendritic cell | ITGAX | + |
| Dendritic cell | CD3D | - |
| Dendritic cell | CD3E | - |
| Dendritic cell | CD3G | - |
| Dendritic cell | NCAM1 | - |
| Dendritic cell | CD19 | - |
| Dendritic cell | CD14 | - |
| Dendritic cell | CD1C | + |
| Dendritic cell | CD1E | + |
| Dendritic cell | FCER1A | + |
| Dendritic cell | CLEC10A | + |
| Dendritic cell | FCGR2B | + |
| Dendritic cell | MS4A1 | - |
| Dendritic cell | CD79A | - |
| Dendritic cell | CD79B | - |
| Plasmacytoid dendritic cell | IL3RA | + |
| Plasmacytoid dendritic cell | GZMB | + |
| Plasmacytoid dendritic cell | JCHAIN | + |
| Plasmacytoid dendritic cell | IRF7 | + |
| Plasmacytoid dendritic cell | TCF4 | + |
| Plasmacytoid dendritic cell | LILRA4 | + |
| Plasmacytoid dendritic cell | CLEC4C | + |
| Plasmacytoid dendritic cell | ITGAX | - |
| Plasmacytoid dendritic cell | CD3D | - |
| Plasmacytoid dendritic cell | CD3E | - |
| Plasmacytoid dendritic cell | CD3G | - |
| Plasmacytoid dendritic cell | NCAM1 | - |
| Plasmacytoid dendritic cell | CD19 | - |
| Plasmacytoid dendritic cell | CD14 | - |
| Plasmacytoid dendritic cell | MS4A1 | - |
| Plasmacytoid dendritic cell | CD79A | - |
| Plasmacytoid dendritic cell | CD79B | - |
| Plasma cell | CD38 | + |
| Plasma cell | XBP1 | + |
| Plasma cell | CD27 | + |
| Plasma cell | SLAMF7 | + |
| Plasma cell | CD19 | - |
| Plasma cell | MS4A1 | - |
| Plasma cell | CD3D | - |
| Plasma cell | CD3E | - |
| Plasma cell | CD3G | - |
| Plasma cell | IGHA1 | + |
| Plasma cell | IGHA2 | + |
| Plasma cell | IGHG1 | + |

Table s12

| | | |
|---|---|---|
| Plasma cell | IGHG2 | + |
| Plasma cell | IGHG3 | + |
| Plasma cell | IGHG4 | + |
| Megakaryocyte | PF4 | + |
| Megakaryocyte | PPBP | + |
| Megakaryocyte | GP5 | + |
| Megakaryocyte | ITGA2B | + |
| Megakaryocyte | NRGN | + |
| Megakaryocyte | TUBB1 | + |
| Megakaryocyte | SPARC | + |
| Megakaryocyte | RGS18 | + |
| Megakaryocyte | MYL9 | + |
| Megakaryocyte | GNG11 | + |

Table s13

| Cell type | Gene | Positive or Negative |
|---|---|---|
| Astrocyte | Slc1a3 | + |
| Astrocyte | Aqp4 | + |
| Astrocyte | Gja1 | + |
| Astrocyte | F3 | + |
| Astrocyte | Aldoc | + |
| Astrocyte | Fgfr3 | + |
| Excitatory neuron | Slc17a7 | + |
| Excitatory neuron | Neurod6 | + |
| Excitatory neuron | Neurod2 | + |
| Excitatory neuron | Tbr1 | + |
| Excitatory neuron | Satb2 | + |
| Excitatory neuron | Cbln2 | + |
| Excitatory neuron | Slc17a6 | + |
| Inhibitory neuron | Gad1 | + |
| Inhibitory neuron | Gad2 | + |
| Inhibitory neuron | Dlx1 | + |
| Inhibitory neuron | Dlx2 | + |
| Inhibitory neuron | Slc32a1 | + |
| Inhibitory neuron | Erbb4 | + |
| Microglia | Csf1r | + |
| Microglia | C1qa | + |
| Microglia | Aif1 | + |
| Microglia | Tmem119 | + |
| Microglia | Ctss | + |
| Microglia | C1qb | + |
| Microglia | Tyrobp | + |
| Microglia | Laptm5 | + |
| Oligodendrocyte | Olig1 | + |
| Oligodendrocyte | Olig2 | + |
| Oligodendrocyte | Mog | + |
| Oligodendrocyte | Mbp | + |
| Oligodendrocyte | Mobp | + |
| Oligodendrocyte | Plp1 | + |
| Oligodendrocyte | Sox10 | + |
| Oligodendrocyte | Gpr37 | + |
| Oligodendrocyte | Mag | + |
| Oligodendrocyte | Cnp | + |
| Oligodendrocyte Precursor | Pdgfra | + |
| Oligodendrocyte Precursor | Cspg4 | + |
| Oligodendrocyte Precursor | Olig1 | + |
| Oligodendrocyte Precursor | Olig2 | + |
| Oligodendrocyte Precursor | Rlbp1 | + |
| Oligodendrocyte Precursor | C1ql1 | + |
| Endothelial cell | Flt1 | + |
| Endothelial cell | Id1 | + |
| Endothelial cell | Foxf2 | + |

Table s13

| | | |
|---|---|---|
| Endothelial cell | Foxq1 | + |
| Endothelial cell | Lef1 | + |
| Endothelial cell | Cldn5 | + |
| Endothelial cell | Zic3 | + |
| Endothelial cell | Ocln | + |
| Pericyte | Vtn | + |
| Pericyte | Pdgfrb | + |
| Pericyte | Atp13a5 | + |
| Pericyte | Rgs5 | + |
| Pericyte | Kcnj8 | + |
| Pericyte | Des | + |

Table s14

| Method | Replicate | # nearest neighbors | variable.gene | resolution | # PCs |
|---|---|---|---|---|---|
| Smart-seq2 | PBMC1 | 5 | FALSE | 0.5 | 20 |
| CEL-Seq2 | PBMC1 | 5 | FALSE | 0.5 | 20 |
| 10x Chromium (v2) A | PBMC1 | 30 | FALSE | 1.5 | 30 |
| 10x Chromium (v2) B | PBMC1 | 15 | TRUE | 0.8 | 20 |
| 10x Chromium (v3) | PBMC1 | 30 | TRUE | 0.8 | 30 |
| Drop-seq | PBMC1 | 15 | FALSE | 1 | 30 |
| Seq-Well | PBMC1 | 15 | TRUE | 1.5 | 30 |
| inDrops | PBMC1 | 30 | TRUE | 1.2 | 30 |
| Smart-seq2 | PBMC2 | 5 | FALSE | 0.5 | 20 |
| CEL-Seq2 | PBMC2 | 5 | TRUE | 0.5 | 20 |
| 10x Chromium (v2) | PBMC2 | 30 | TRUE | 1.5 | 50 |
| Drop-seq | PBMC2 | 30 | TRUE | 0.8 | 20 |
| Seq-Well | PBMC2 | 15 | FALSE | 1.2 | 30 |
| inDrops | PBMC2 | 15 | TRUE | 1.5 | 20 |
| Smart-seq2 | Cortex1 | 5 | TRUE | 1.5 | 30 |
| 10x Chromium (v2) | Cortex1 | 30 | FALSE | 1.2 | 20 |
| DroNc-seq | Cortex1 | 10 | FALSE | 1.5 | 50 |
| sci-RNA-seq | Cortex1 | 30 | FALSE | 1.2 | 20 |
| Smart-seq2 | Cortex2 | 5 | TRUE | 1 | 20 |
| 10x Chromium (v2) | Cortex2 | 30 | TRUE | 0.8 | 100 |
| DroNc-seq | Cortex2 | 5 | TRUE | 0.5 | 30 |
| sci-RNA-seq | Cortex2 | 30 | TRUE | 1 | 100 |
